# A single-cell transcriptional timelapse of mouse embryonic development, from gastrula to pup

**DOI:** 10.1101/2023.04.05.535726

**Authors:** Chengxiang Qiu, Beth K. Martin, Ian C. Welsh, Riza M. Daza, Truc-Mai Le, Xingfan Huang, Eva K. Nichols, Megan L. Taylor, Olivia Fulton, Diana R. O’Day, Anne Roshella Gomes, Saskia Ilcisin, Sanjay Srivatsan, Xinxian Deng, Christine M. Disteche, William Stafford Noble, Nobuhiko Hamazaki, Via B. Moens, David Kimelman, Junyue Cao, Alexander F. Schier, Malte Spielmann, Stephen A. Murray, Cole Trapnell, Jay Shendure

## Abstract

The house mouse, *Mus musculus*, is an exceptional model system, combining genetic tractability with close homology to human biology. Gestation in mouse development lasts just under three weeks, a period during which its genome orchestrates the astonishing transformation of a single cell zygote into a free-living pup composed of >500 million cells. Towards a global framework for exploring mammalian development, we applied single cell combinatorial indexing (sci-*) to profile the transcriptional states of 12.4 million nuclei from 83 precisely staged embryos spanning late gastrulation (embryonic day 8 or E8) to birth (postnatal day 0 or P0), with 2-hr temporal resolution during somitogenesis, 6-hr resolution through to birth, and 20-min resolution during the immediate postpartum period. From these data (E8 to P0), we annotate dozens of trajectories and hundreds of cell types and perform deeper analyses of the unfolding of the posterior embryo during somitogenesis as well as the ontogenesis of the kidney, mesenchyme, retina, and early neurons. Finally, we leverage the depth and temporal resolution of these whole embryo snapshots, together with other published data, to construct and curate a rooted tree of cell type relationships that spans mouse development from zygote to pup. Throughout this tree, we systematically nominate sets of transcription factors (TFs) and other genes as candidate drivers of the *in vivo* differentiation of hundreds of mammalian cell types. Remarkably, the most dramatic shifts in transcriptional state are observed in a restricted set of cell types in the hours immediately following birth, and presumably underlie the massive changes in physiology that must accompany the successful transition of a placental mammal to extrauterine life.

## Introduction

The last ten years have witnessed the development and application of single-cell molecular profiling technologies to characterize biological development at the scale of the whole organism^1–10^. Most of these studies comprise time series, wherein each embryo profiled with single-cell RNA-seq (scRNA-seq) or ATAC-seq (scATAC-seq) captures one moment in developmental time, yielding “snapshots” that must be pieced together analogous to a timelapse movie. Inevitably, there are tradeoffs between the span of development studied and both the frame rate and resolution of the snapshots taken. For example, in the mouse, two studies intensely profiled gastrulating embryos, together quantifying gene expression in 150,000 cells from over 500 embryos spanning E6.5 to E8.5^6, 10^, while a study from our group profiled 2 million nuclei from 61 embryos staged at roughly 24-hr intervals spanning E9.5 to E13.5^7^. We systematically integrated these and other scRNA-seq snapshots of whole mouse embryos, overcoming various batch effects to obtain a coarse tree relating mammalian transcriptional states from E3.5 to E13.5^11^. However, these studies did not examine later stages of prenatal development, which are more difficult to subject to whole embryo single cell profiling due to the sheer number of cells as well as the emergence of bone, which is difficult to pulverize.

Here we set out to deeply profile the molecular states of single nuclei derived from precisely staged mouse embryos spanning late gastrulation (E8) to birth (P0). The resulting dataset greatly improves upon our previously reported single cell atlas of early organogenesis in the mouse^7^ with respect to: (a) the number of single nucleus profiles obtained (2 million → 12.4 million single nuclei), (b) the depth of profiling (median 671 → 2,545 unique molecular identifiers [UMIs] per nucleus), (c) temporal resolution (24-hr → 20-min, 2-hr or 6-hr intervals), and (d) the span of development covered (E9.5 to E13.5 → E8 to P0).

We focus here on describing the dataset and our early analyses, which include the preliminary annotation of hundreds of cell types and deeper dives into the ontogenesis of selected systems. Improving upon our previously described approach^11^, we also leverage these and other published scRNA-seq snapshots of whole mouse embryos to curate a data-driven tree of mouse development that relates cell types throughout prenatal development, as well as to nominate sets of TFs and other genes as candidate drivers of the *in vivo* differentiation of each mammalian cell type. Surprisingly, we find that birth (E18.75 → P0) is associated with a dramatic shift in transcriptional state in a restricted subset of cell types. A more detailed time-course of additional embryos at 20-min intervals in the immediate postpartum period confirmed the rapidity of these transitions and highlighted specific ways in which rapidly activated gene expression programs in specific cell types may support the adaptation from fetal to extrauterine life. Altogether, our studies provide a framework for exploring the entirety of mammalian embryogenesis, from single cell zygote to newborn pup.

## Ontogenetic Staging and Embryo Selection

Towards a more continuous view of single cell transcriptional dynamics throughout embryonic development, we sought to profile whole mouse embryos at a higher “frame rate” than the 24-hr intervals of our previous study of early organogenesis^7^. To enable rigorous comparisons between embryos within our sample set, we distinguish between gestational stage and absolute developmental stage of individual embryos harvested. Mouse gestational age, based upon the observation of a vaginal plug for which noon on that day is counted as E0.5, provides a loose approximation of the time elapsed from conception to the time a litter is harvested. Importantly, stochastic differences in timing of mating or fertilization, and genetic factors such as strain/species-dependent gestational length and litter size, can result in significant intra- and inter-litter variation of embryos of an identical gestational stage^12^. Conversely, embryonic morphogenesis is highly ordered, reproducible, and inherently reflective of an embryo’s developmental age with respect to absolute position within a morphogenetic trajectory and the dynamic progression of underlying cellular transcriptional states^1, 13, 14^. Therefore, we sought to stage individual embryos based on well-defined morphological criteria. Each embryo was assigned to one of 45 temporal bins at 6-hr increments from E8 to P0 (Fig. 1a; Supplementary Figs. 1-2). For the earlier stages (before E10), embryos were staged based on morphological features (Supplementary Fig. 1) and somites counted at the time of harvest, facilitating temporal binning at roughly 2-hr increments. For E10.25 to E14.75 embryos, developmental age was determined by measurements of hindlimb bud development using the Embryonic Mouse Ontogenetic Staging System (eMOSS), which leverages dynamic changes in hindlimb bud morphology and a pattern recognition algorithm to estimate the absolute stage of a sample^15, 16^. Due to the increased complexity of limb morphology at later stages, automated staging beyond E15 is not possible. Therefore, harvests for all remaining embryonic samples (E15 to E18.75) were performed precisely at 00:00, 06:00, 12:00, and 18:00 on the targeted gestational day. From close inspection of limbs in this set, we defined additional dynamics related to digit morphogenesis that allowed further binning of samples collected on days 15 and 16 (Supplementary Fig. 2). The remaining timepoints (E17 and beyond) were staged based on hour of harvest and gestational age.

**Figure 1.**
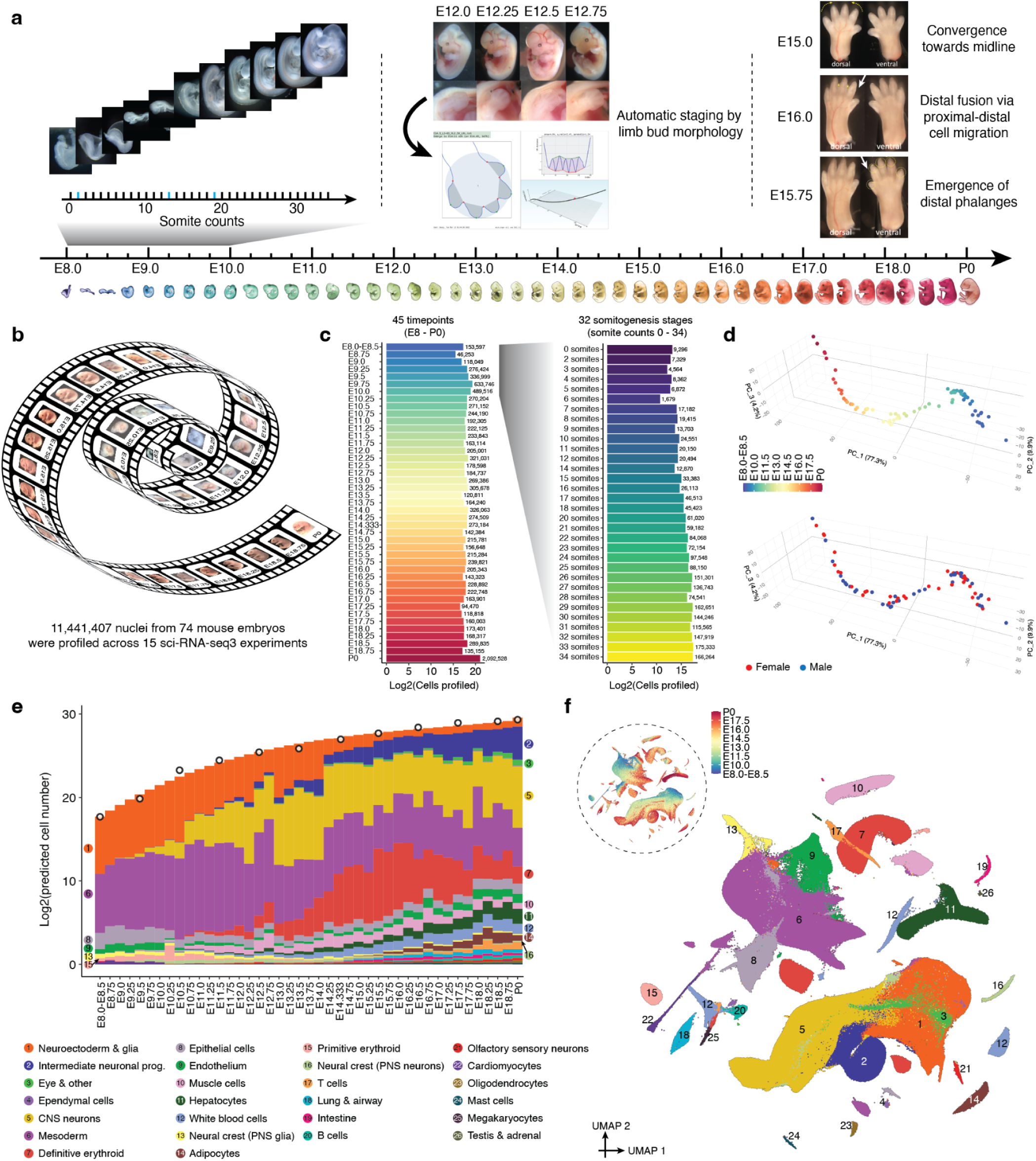
A single cell transcriptional timelapse of mouse development, from gastrula to pup. **a,** Embryos were collected and precisely staged based on morphological features, including by counting somite numbers (up to E10) and an automated process that leverages limb bud geometry (E10-E15) (**Methods**). Each embryo was assigned to one of 45 temporal bins at 6-hr increments from E8 to P0, and to more highly resolved 2-hr bins at earlier timepoints based on somite counts. The first three bins (E8.0, E8.25, E8.5) are combined for presentation purposes, and this is also the subset of the data that was previously reported^11^. Embryos with somite counts 1, 3, 19 are missing from the series (blue ticks in sub-axis at left). See Supplementary Figs. 1-2 and **Supplementary Table 1** for further details. **b,** Across 15 sci-RNA-seq3 experiments, we generated single nucleus profiles for 11.4M cells from 74 embryos spanning mouse development from late gastrulation to birth, with around 2-hr temporal resolution from 0 to 34 somites, and 6-hr resolution through to birth. **c,** The number (log2 scale) of nuclei profiled at each timepoint, shown for 6-hr bins at left and for 2-hr bins (somitogenesis) at right. **d,** Embeddings of pseudo-bulk RNA-seq profiles of 74 mouse embryos in PCA space with visualization of top three PCs. Briefly, single nucleus data from each embryo was aggregated to create 74 profiles, with which we performed dimensionality reduction via PCA. Embryos are colored by either developmental stage (top) or data-inferred sex (bottom). **e,** Composition of embryos from each 6-hr bin by major cell cluster. The y-axis is scaled to the estimated cell number (log2 scale) at each timepoint (**Methods**). Briefly, we isolated and quantified total genomic DNA from whole embryos to estimate the total cell number at each of 12 stages, spanning from E8.5 to P0 (1-day bins, highlighted by black circles), and then applied polynomial regression to predict log2-scaled cell number of the whole embryo at each of the 43 timepoints (P0 was treated as E19.5 in the regression). CNS: central nervous system. PNS: peripheral nervous system. **f,** 2D UMAP visualization of the whole dataset. Colors and numbers correspond to 26 major cell cluster annotations as listed in the key of panel e. Dashed circle inset at top left shows the same UMAP colored by developmental stage (plotting a uniform number of cells per timepoint).

From a total of 523 staged embryos, we selected 75 for whole embryo scRNA-seq, aiming for one embryo for every somite count from 0 to 34, and then one embryo for every 6-hr bin from E10 to P0 (**Supplementary Table 1**). Embryos with somite counts 1, 13, and 19 are missing from the series, while a handful of post-E10 timepoints are represented by 2-3 individual embryos. Wherever possible, we sought to alternate between male and female embryos in neighboring bins; for the final P0 timepoint, both sexes were deeply profiled.

## Single Nucleus Transcriptional Profiling of Whole Embryos

Frozen embryos were processed via a recently optimized protocol for three-level single cell transcriptional profiling by combinatorial indexing (sci-RNA-seq3)^17^. In each of 15 sci-RNA-seq3 experiments, we generally sought to include embryos from adjacent stages of development (**Supplementary Table 1**). Sequencing data was generated across 21 runs of the Illumina NovaSeq instrument, with all libraries sequenced to either a PCR duplication rate of >50% or a median UMI count of >2,500 per nucleus (**Supplementary Table 2**). Reads from each sci-RNA-seq3 experiment were demultiplexed, trimmed and mapped to the mouse reference genome (mm10). This was followed by the removal of PCR duplicates. Finally, poor quality nuclei and potential doublets were aggressively filtered (**Methods**)^7, 11^. As nuclei from only one embryo are deposited to each well during the first indexing round, we retain the identity of the embryo from which each single nucleus profile is derived^7^.

The cell-by-gene count matrix that we carried forward combines profiles from all 15 experiments, and includes transcriptional profiles for 11,441,407 nuclei from 74 embryos spanning E8 to P0 (Fig. 1b). Of note, ∼1% of these single nucleus profiles (data from embryos with somite counts 0-12) were reported in a previous publication from our group^11^. On average, 154,614 nuclei were profiled per embryo, with recovery generally correlated with stage (Fig. 1c; **Supplementary Table 1**; median 142,854; range = 1,679 to 1,613,834). The median UMI count obtained per nucleus was 2,700 and the median number of genes detected was 1,574 (Supplementary Fig. 3a). Further analyses suggested that batch effects were relatively minimal (Supplementary Fig. 3b). Consistent with this, a principal component analysis (PCA) of pseudo-bulked RNA-seq profiles corresponding to each timepoint resulted in a major first component (PC1; 77.3%) strongly correlated with developmental time (Fig. 1d).

What kind of “coverage” of all cells in the mouse embryo are we achieving here? As the total number of cells in developing mouse embryos is not well documented, we sought to experimentally estimate this by quantifying total DNA content across a series of staged embryos. In brief, we estimate that the embryo grows 3,000-fold between E8.5 and P0 (210K → 670M cells), with the cellular ‘doubling time’ of the embryo slowing from around 6 hours to 1.5 days across the same interval (Fig. 1e; **Supplementary Table 3**; **Methods**). Thus, even with the large number of nuclei profiled here, our “cellular coverage” of the embryo remains modest, ranging from 0.5-fold for early stages (summing across 6 embryos with somite counts 7 to 12) to 0.002-fold immediately before birth (summing across 6 embryos staged E17.5 to E18.75).

## Preliminary Annotation of Major Cell Clusters and Cell Types

To get our bearings on this very large cell-by-gene count matrix, we used *Scanpy*^18^ to generate a global embedding of the 11,441,407 nuclei (hereafter referred to as cells) from all experiments and timepoints, and then annotated 26 major cell clusters based on marker genes (Fig. 1e-f; **Supplementary Table 4**). The major cell clusters assigned the most cells were the mesoderm (n = 3,267,338), central nervous system (CNS) neurons (n = 2,106,206), neuroectoderm & glia (n = 1,733,663) and definitive erythroid (n = 1,033,409) lineages.

We emphasize that the resolution of these major cell clusters is somewhat arbitrary and impacted by abundance, *e.g.* we have separate major cell clusters for B, T and mast cells, but the remainder of blood cells, including hematopoietic stem cells (HSCs), are lumped into ‘white blood cells’; intermediate neuronal progenitors and their derivatives are a major cell cluster, while nearly all other CNS neurons are lumped into a separate major cell cluster. Although most major cell clusters were very straightforward to annotate based on the literature and our previous experience with mouse scRNA-seq datasets up to E13.5^7, 11^, a few first appear only late in organogenesis, including adipocytes, testes & adrenal cells (Fig. 1e). As expected, cell clusters whose proportions decline over time either stream towards derivatives (*e.g.* neuroectoderm & glia → CNS neurons, intermediate neuronal progenitors) or are displaced by a functionally analogous but developmentally distinct lineage (*e.g.* primitive erythroid → definitive erythroid).

We next performed sub-clustering of cells within each of the 26 major cell clusters, followed by annotation of each cluster based on marker genes, resulting in 190 labeled cell types (Supplementary Fig. 4; **Supplementary Table 5**). These annotations are preliminary, and subject to refinement or correction as we and others further explore the data^7, 11^.

We next sought to leverage this single cell timelapse of mouse development from gastrula to pup to perform deeper dives into the unfolding of the posterior embryo during somitogenesis, as well as the ontogenesis of the kidney, mesenchyme, retina, and early neurons.

## The Posterior Embryo During Somitogenesis

Neuromesodermal progenitors (NMPs) are a fascinating population of bipotent cells that give rise to the spinal cord as well as trunk and tail somites^19^. We previously profiled late-gastrulation embryos staged in one-somite increments (0 to 12 somites) and investigated transcriptional heterogeneity among early NMPs^11^. With the goal of extending this analysis to later time points and also relating NMPs to other physically coincident cell types in the posterior embryo, we extracted and re-embedded cells from all somite-staged embryos (0 to 34 somites) annotated as NMPs & spinal cord progenitors, mesodermal progenitors (*Tbx6*+), notochord or gut (Fig. 2a-b).

**Figure 2.**
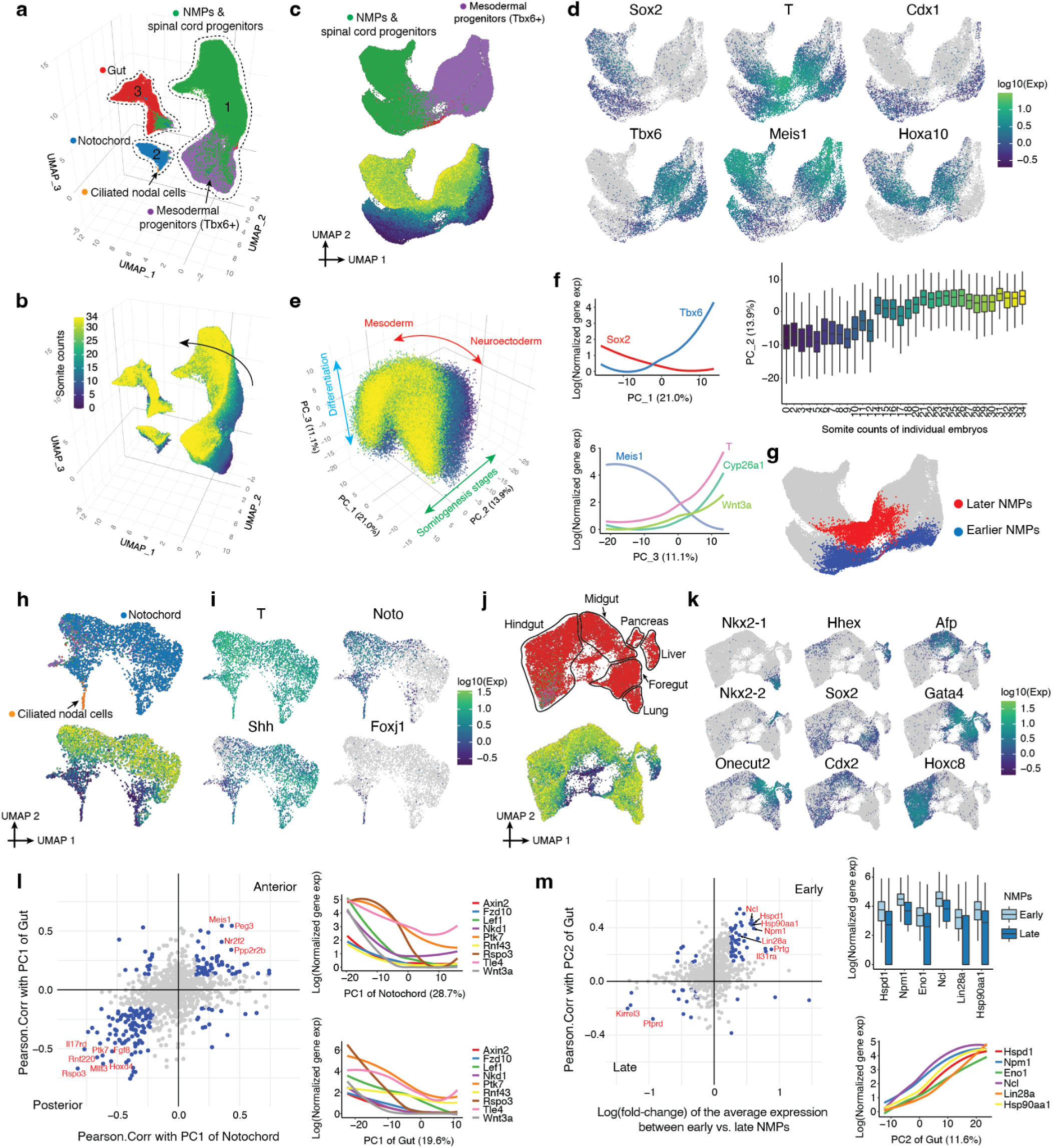
Transcriptional heterogeneity in the posterior embryo during the early somitogenesis. **a,** Re-embedded 3D UMAP of 121,118 cells which were initially annotated as neuromesodermal progenitors (NMPs) & spinal cord progenitors, mesodermal progenitors (*Tbx6*+), notochord, ciliated nodal cells, or gut, from embryos during the early somitogenesis (somite counts 0-34; E8-E10). Cells are colored by their initial annotations. Three clusters are identified and highlighted, with cluster 1 dominated by NMPs and their derivatives, cluster 2 dominated by notochord and ciliated nodal cells, and cluster 3 dominated by gut. **b,** The same UMAP as in panel a, colored by somite counts. Arrow highlights the increment of somite counts. **c,** Re-embedded 2D UMAP of cells from cluster 1 in panel a. Cells are colored by either their initial annotations (top) or somite counts (bottom). **d,** The same UMAP as in panel c, colored by gene expression of marker genes which appear specific to different subpopulations of NMPs: column 1: differences between neuroectodermal (*Sox2*+) vs. mesodermal (*Tbx6*+) fates^35^; column 2: the differentiation of bipotential NMPs (*T*+, *Meis1*-) towards either fate^36, 37^; column 3: earlier (*Cdx1*+) vs. later (*Hoxa10*+) NMPs^24^. **e,** 3D visualization of the top three principal components (PCs) of gene expression variation in cells from cluster 1, calculated on the basis of the 2,500 most highly variable genes. Cells are colored by the somite count of the embryo from which they derived. As further shown in panel f, PC1 correlates with mesodermal vs. neuroectodermal fates (red arrow), PC2 with somite stage (green arrow), and PC3 with the progression of differentiation of bipotential NMPs towards either fate (blue arrow). **f,** Correlations between top three PCs and the normalized expression of selected genes (*Sox2* & *Tbx6* for PC1; *T*, *Cyp26a1*, *Wnt3a* and *Meis1* for PC3) or somite counts (for PC2). Gene expression values were calculated from original UMI counts normalized to total UMIs per cell, followed by natural-log transformation. The line of gene expression was plotted by the *geom_smooth* function in ggplot2. In the boxplot, the center lines show the medians; the box limits indicate the 25th and 75th percentiles. The genes significantly correlated with each PC are shown in **Supplementary Table 6**. **g,** The same UMAP as in panel c, with earlier (blue, *n* = 4,949 cells) and later (red, *n* = 3,910 cells) NMPs highlighted. Here cells are labeled as NMPs if they are both strongly *T*+ (raw count >= 5) & *Meis1*- (raw count = 0). **h,** Re-embedded 2D UMAP of cells from cluster 2 in panel a. Cells are colored by either the initial annotation (top) or somite counts (bottom). **i,** The same UMAP as in panel h, colored by gene expression of marker genes which appear specific to notochord (*T*+, *Noto*+, *Shh*+)^38, 39^ or ciliated nodal cells (*Foxj1*+)^38^. **j,** Re-embedded 2D UMAP of cells from cluster 3 in panel a. Cells are colored by either the initial annotations (top) or somite counts (bottom). Different subpopulations of gut cells are highlighted by black circles. **k,** The same UMAP as in panel j, colored by gene expression of marker genes which appear specific to different subpopulations of gut cells, including lung (*Nkx2-1*+)^40^, hepatocytes (*Hhex*+, *Afp*+)^41, 42^, pancreas (*Nkx2-2*+)^43^, foregut (*Gata4*+, *Sox2*+)^44, 45^, midgut (*Gata4*+, *Onecut2*+)^45, 46^, and hindgut (*Cdx2*+, *Hoxc8*+)^47–49^. **l,** On the left, the Pearson correlation coefficient between gene expression for the top highly variable genes and either PC1 of notochord (x-axis) or PC1 of gut (y-axis). For each cell cluster, the top 2,500 highly variable genes were identified and their gene expression values were calculated from original UMI counts normalized to total UMIs per cell, followed by natural-log transformation and scaling. After performing Pearson correlation with the selected PC, significant genes were identified if their correlation coefficients are < mean + 1 standard deviation of all the correlation coefficients, and FDR < 0.05. The overlapped genes between two cell clusters are shown as each dot, and the overlapped significant genes highlighted in blue. The first quadrant corresponds to the inferred anterior aspect of each cluster, while the third quadrant corresponds to the inferred posterior aspect. On the right, gene expression of selected genes involved in Wnt signaling are plotted over PC1 of notochord (top) or PC1 of gut (bottom). Gene expression values were calculated from original UMI counts normalized to total UMIs per cell, followed by natural-log transformation. The line of gene expression was plotted by the *geom_smooth* function in ggplot2. **m,** On the left, the log-scaled fold-change of the average expression for the top highly variable genes between early vs. late NMPs (x-axis), and the Pearson correlation coefficient between gene expression for the top highly variable genes and PC2 of gut (y-axis). The first quadrant is associated with early somite counts for each cluster, while the third quadrant is associated with late somite counts. On the right, gene expression of selected genes (several Myc targets^50^, *Lin28a*, *Hsp90aa1*) are plotted between early vs. late NMPs (top) or over PC2 of gut (bottom). The highly differentially expressed genes between early vs. late NMPs were identified using the *FindMarkers* function of Seurat/v3^51^, after filtering out genes that are detected in < 10% of cells in both of the two populations. Significant genes were identified if their absolutely log-scaled fold-changes > 0.25, and adjusted p-values < 0.05.

Focusing first on the subset of cells annotated as either ‘NMPs & spinal cord progenitors’ or ‘mesodermal progenitors (*Tbx6*+)’ (cluster 1 in Fig. 2a), we performed PCA on highly variable genes. The top three PCs, which explain nearly half of transcriptional variation in these cells, appear to correspond to the differences between neuroectodermal vs. mesodermal fates (PC1), developmental stage (PC2), and the differentiation of bipotential NMPs towards either fate (PC3) (Fig. 2c-f; **Supplementary Table 6**). Assuming that PC3 tracks the progression of differentiation consistently between neuroectodermal and mesodermal fates, the data suggest that the state of being *T* (*Brachyury*)+, *Meis1*- (Fig. 2f-g) may be a better indicator of bi-potency than being *T+*, *Sox2*+^20^, consistent with two recent *in vivo* studies of the genetic dependencies of NMPs^21, 22^. In addition to *T*, other markers of NMPs that were strongly correlated with the undifferentiated state include *Cyp26a1* and *Wnt3a*^23^.

We observe striking contrasts in gene expression between NMPs derived from earlier (0-12 somites) vs. later (14-34 somites) embryos (Fig. 2c-g). Although we initially worried that this was a batch effect (as these embryos were processed in separate experiments), the observation is consistent with a study in which NMPs obtained from microdissected E8.5 and E9.5 embryos exhibited strong differences^24^, with many of the same genes exhibiting sharp differential expression here (*Cdx1* (early); *Hoxa10* (late); Fig. 2d; **Supplementary Table 7**). Overall, our results are consistent with these early vs. late NMPs underlying the “trunk-to-tail” transition^25^. Given that early and late somite count embryos were processed in different experiments, we profiled an additional 12 mouse embryos ranging in somite counts from 8 to 21 via sci-RNA-seq. This experiment validated and refined the estimated timing of this transition (Supplementary Fig. 5).

In addition to NMPs, another cell type marked by *T* is the notochord (cluster 2 in Fig. 2a). In the earliest embryos of this time course (0-12 somites), we observe two transcriptionally distinct subsets of cells within the notochord cluster, one more strongly *Noto*+ and the other more strongly *Shh*+ (Fig. 2h-i). As embryogenesis progresses, these subsets transition to a continuum, but the distinctions are preserved and reinforced, with the inferred derivatives of one early subpopulation marked by the expression of *Noto*, posterior *Hox* genes, and genes involved in Notch signaling, Wnt signaling and mesodermal differentiation (Supplementary Fig. 6a). Within this first early subpopulation, we also detect a highly distinct cluster of 60 cells that strongly express *Foxj1* and genes involved in motile ciliogenesis; these ciliated nodal cells are transient, peaking in terms of their contribution to the overall embryo at the 2-somite stage (Fig. 2h-i; Supplementary Fig. 6b).

In contrast, the inferred derivatives of second early subpopulation, marked by stronger *Shh* expression, is enriched for genes involved in neurogenesis and synaptogenesis, notably including *Sox10*, *Bmp3*, *Nrg1* and *Erbb4* (Supplementary Fig. 6c). One possibility is that the *Noto*+ subpopulation corresponds to the posterior notochord, which arises from the node, and the *Shh*++ subpopulation corresponds to the anterior mesendoderm (*i.e.* anterior head process and possibly the prechordal plate), which arises by condensation of dispersed mesenchyme and may contribute to forebrain patterning^26–29^. Performing PCA (after excluding ciliated nodal cells), we find that these presumably A-P differences are the predominant source of transcriptional heterogeneity in the notochord cluster (PC1; 28.7% of variation; **Supplementary Table 8**).

Turning to the gut (cluster 3 in Fig. 2a), we once again observe subsets of transcriptionally distinct progenitors at early somite stages that give rise to a continuum at later somite stages (Fig. 2j). A major axis of this continuum reflects A-P patterning, with subsets corresponding to lung (*Nkx2-1*+), hepatocytes (*Afp*+, *Hhex*+), pancreas (*Nkx2-2*+), foregut (*Sox2*+, *Gata4*+), midgut (*Gata4*+, *Onecut2*+) and hindgut (*Hoxc8*+, *Cdx2*+) progenitors (Fig. 2k; PC1; 19.6% of variation; **Supplementary Table 9**). As *T* expression is classically associated with the notochord and posterior mesoderm in the mouse literature, we were initially surprised to see strong *T* expression in the inferred posterior hindgut, coincident with the expression of posterior *Hox* genes (Supplementary Fig. 6d). However, this expression pattern is consistent with the ancestral role of *T* in the closing of the blastopore^30^ and hindgut defects in *Drosophila brachyenteron* and *Caenorhabditis elegans mab-9* mutants^31, 32^.

Finally, we sought to explore whether there are overlaps between genes associated with spatial patterning (A-P axis) or developmental progression (somite counts) across germ layers. In comparing genes whose expression patterns are highly correlated with the inferred A-P axis between notochord (PC1; n=591) and gut (PC1; n=502), we observe striking overlap and directional concordance (198 overlapping genes, 86% of which are consistently associated with the inferred anterior or posterior aspect of the notochord and gut; p<1e-28, χ2-test; Fig. 2l; **Supplementary Table 10**). Concordant, posterior-associated genes are highly enriched for genes involved in Wnt signaling as well as posterior *Hox* genes. One model to explain these overlaps between germ layers is that they are residual to the common origin of anterior mesendodermal derivatives (anterior head process, prechordal plate, anterior endoderm) from the early & mid-gastrula organizers vs. posterior mesendodermal derivatives (notochord and posterior endoderm) from the node^26^. Alternatively, these overlaps could be explained by physically coincident progenitors of these germ layers being exposed to similar patterns of Wnt signaling (*e.g.* strength, timing).

A second striking overlap between germ layers involves genes highly correlated with early vs. late somite counts in NMPs (n=257) vs. the gut (PC2; n=502). Once again, we observe striking overlap and directional concordance (82 overlapping genes, 70 (85%) of which are consistently associated with early or late somite counts; p<1e-15, χ2-test) (Fig. 2m; **Supplementary Table 11**). Given that early and late somite count embryos were processed in different experiments, we examined the additional 12 mouse embryos ranging in somite counts from 8 to 21. In this replication dataset, 77% of the initially overlapping, concordant genes replicated in terms of directionality-of-change between early (<13) vs. late (>13) somite count embryos in both NMPs and the gut (54/70; expected 25%; Supplementary Fig. 5). Genes reproducibly associated with early somite counts in both NMPs and the gut were strongly enriched for Myc targets, and also included *Lin28a*, a deeply conserved regulator of developmental timing^33, 34^, and multiple isoforms of Hsp90 (Fig. 2m; **Supplementary Table 11**), possibly reflecting greater proliferation.

## Diversification of the Intermediate and Lateral Plate Mesoderm

The mesoderm, which includes axial, paraxial, intermediate and lateral plate components, is a complex germ layer, with the ontogenesis of some derivatives (*e.g.* heart, blood) much better understood than others (*e.g.* the mesenchyme that plays a major role in patterning each organ). In the previous section, we considered aspects of the axial (notochord) and paraxial (somite-forming) mesoderm. In this section, we focus on the transition from intermediate mesoderm to the kidney, as well as on the transition from the lateral plate mesoderm (LPM) to organ-specific mesenchyme.

What is the continuum of transcriptional states that span the transition from intermediate mesoderm to a functional nephron? When we generate a new embedding from relevant cell types, we observe two major trajectories. The first corresponds to the maturation of the posterior intermediate mesoderm → metanephric mesenchyme → nephron progenitors → renal tubule; while the second corresponds to the maturation of the anterior intermediate mesoderm → ureteric bud → collecting duct (Fig. 3a-c). The specification of these posterior and anterior trajectories in late gastrulation is initiated by interactions between *Gdnf* and *Ret*^52^, followed by their progression to metanephric mesenchyme and the ureteric bud, respectively, around E10.25, and then to specific functional components of the nephron (Fig. 3a-c; Supplementary Fig. 7a-c). Even as their transcriptomes mature with time, the metanephric mesenchyme and ureteric bud persist through P0 (clusters 6 & 3 in Fig. 3a-b; Supplementary Fig. 7b), presumably providing an ongoing source of progenitors for nephrogenesis, which continues for a few days after birth^53^. The apparent bifurcation of the proximal tubule corresponds to major differences in the transcriptional state of cells from embryos obtained before birth (E18.75 or earlier) vs. after birth (P0) (cluster 9 in Fig. 3a-b; Supplementary Fig. 7d). We return to this observation in the final section of the manuscript.

**Figure 3.**
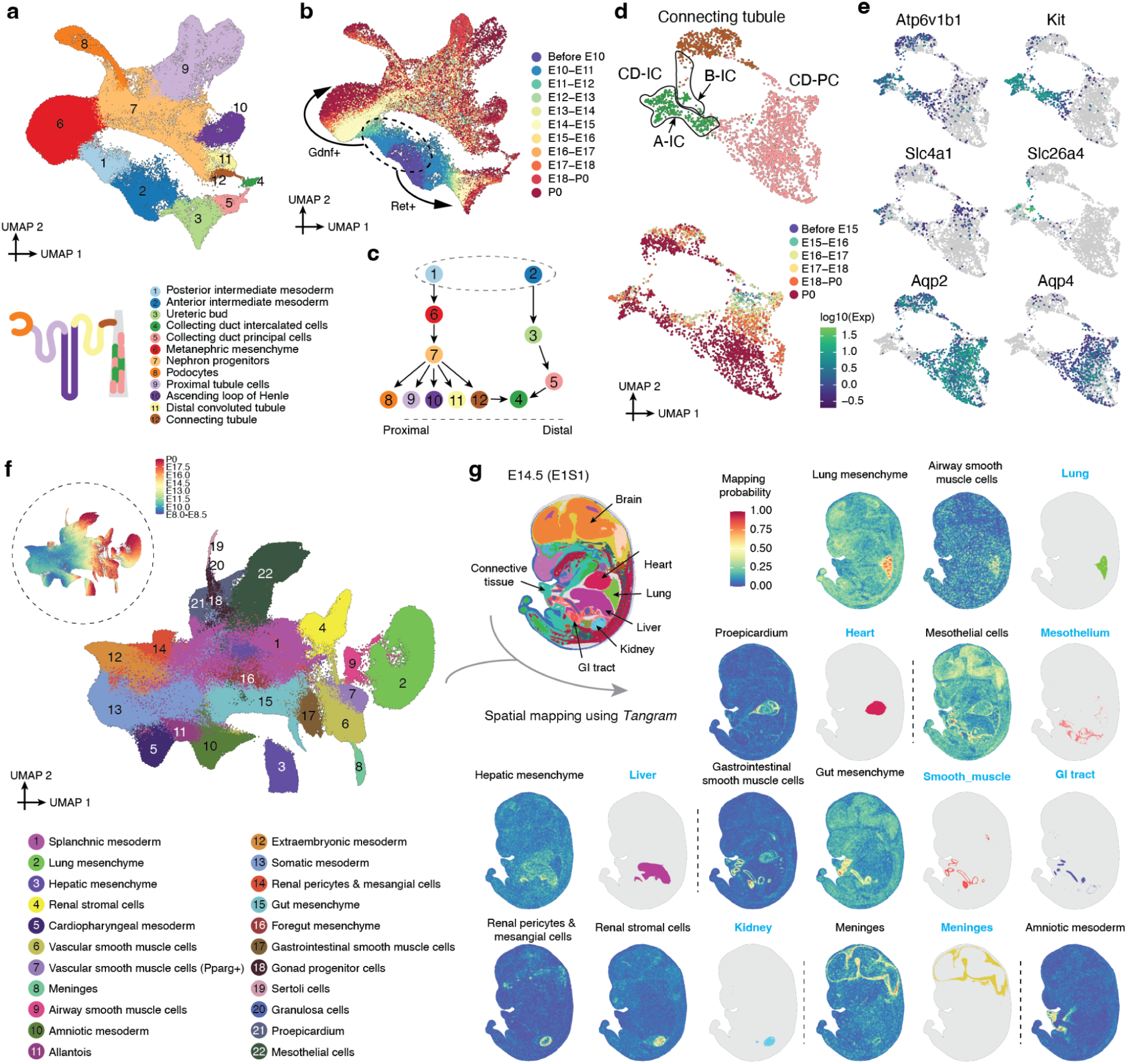
Diversification of the intermediate and lateral plate mesoderm. **a,** Re-embedded 2D UMAP of 95,226 cells corresponding to renal development. A schematic of a nephron is shown at the bottom left. **b,** The same UMAP as in panel a, colored by developmental stage (after downsampling to a uniform number of cells per time window). Dashed circle highlights posterior & anterior intermediate mesoderm, with arrows highlighting derivative trajectories expressing *Gdnf* and *Ret*, respectively. **c,** Manually inferred relationships between annotated renal cell types. Label annotations in panel a. Dashed circle highlights posterior & anterior intermediate mesoderm. Dashed line highlights the expected spatial ordering of annotated cell types from proximal (left) to distal (right) aspect of nephron. **d,** Re-embedded 2D UMAP of 2,894 cells from connecting tubule cells, collecting duct principal cells (CD-PC), and collecting duct intercalated cells (CD-IC). Cells are colored by either their initial annotations (top) or timepoint (bottom). Black circles highlight the cells which appear to be either type A (A-IC) or type B (B-IC) intercalated cells. **e,** The same UMAP as in panel d, colored by expression of marker genes specific to CD-IC (*Atp6v1b1*+), A-IC (*Kit*+, *Slc4a1*+), B-IC (*Slc26a4*+), CD-PC (*Aqp2*+, *Aqp4*+), and connecting tubule (*Aqp2*+, *Aqp4*-) ^63, 64^. **f,** Re-embedded 2D UMAP of 745,494 cells from lateral plate & intermediate mesoderm derivatives. Dashed circle at the top left highlights the same UMAP with cells colored by developmental stage. **g,** To infer the spatial origin of each lateral plate & intermediate mesoderm derivative shown in panel f, we leveraged a public dataset, *Mosta*, which profiles spatial transcriptomes for 53 sections of mouse embryos spanning 8 timepoints from E9.5 to E16.5^59^, together with our data and the *Tangram* algorithm^60^. In brief, for each timepoint of the *Mosta* data, we combined scRNA-seq data from three adjacent timepoints from our data (*e.g.* E16.25, E16.5, and E16.75 from scRNA-seq vs. E16.5 from *Mosta* data), and the total number of voxels within each section was randomly downsampled to 9,000 for computational efficiency. After applying *Tangram*, for each section, a cell-by-voxel matrix with mapping probabilities was returned. To reduce noise, we further smoothed the mapping probabilities for each voxel by averaging values of their *k* nearest neighboring voxels (*k* is calculated by natural-log scaled total number of voxels on that section) followed by scaling it to 0→1 across voxels of each section. An image of one selected section (E1S1) from E14.5 of the *Mosta* data is shown on the top left with major regions labeled. The spatial mapping probabilities across voxels on this section for selected subtypes within the lateral plate & intermediate mesoderm derivatives are shown (black titles), with the regional annotation appearing to best correspond to the inferred spatial pattern shown alongside (blue titles). The mapping results for the other sections are provided here.

In the anterior intermediate mesoderm, both tip and stalk cells are identified within the ureteric bud, the former of which gives rise to the collecting duct, and the latter to the ureter^54, 55^ (Supplementary Fig. 7e). Of note, we observe “convergence” of the posterior and anterior trajectories in collecting duct intercalated cells (cluster 4 in Fig. 3a-b). More detailed investigation suggests that the posterior intermediate mesoderm may also contribute to the collecting duct, although lineage analysis would be necessary to confirm this (Fig. 3d-e; Supplementary Fig. 7f).

The LPM is arguably much more complex than the axial, paraxial and intermediate mesoderms, giving rise to a remarkable diversity of cell types^56^. Although some LPM derivatives are intensely studied (*e.g.* heart), others remain poorly understood, in particular the mesoderm that lines the body wall and internal organs. Not only will this aspect of the LPM go on to give rise to important cell types and structures (*e.g.* fibroblasts, smooth muscle, mesothelium, pericardium, adrenal cortex, genital ridge, etc.), its interactions with other germ layers play a key role in organogenesis, *e.g.* reciprocal signaling between the mesoderm and endoderm serves as the basis of foregut patterning^57, 58^.

To annotate derivatives of the splanchnic mesoderm in particular (*i.e.* the visceral layer of the LPM), we leveraged published spatial transcriptome data from E9.5-E16.5 mice to impute spatial coordinates for cells in our data^59, 60^. Through a combination of spatial inference and marker gene analysis, we were able to assign annotations to 22 subtypes of the LPM & intermediate mesoderm major cell type (Fig. 3f-g; Supplementary Fig. 8; **Supplementary Table 12**). For example, we can define subsets of the splanchnic mesoderm that lines specific organs, including the heart (proepicardium), brain (meninges), lung, liver, foregut and gut, as well as organ-specific smooth muscle (airway vs. gastrointestinal vs. vascular) (Fig. 3g). We can also distinguish two subsets of LPM-derivatives mapping to the kidney, one to the inside and the other to the surface, which may correspond to renal stroma and the renal pericytes and mesangial cells, respectively (Fig. 3g).

The high resolution of our time course during early somitogenesis enables us to narrow the temporal windows during which many subtypes of organ-specific mesenchyme are specified (Supplementary Fig. 9a). We also applied a mutual nearest neighbors (MNN)-based approach to identify putative precursor cells of each subtype. We performed this analysis separately for subtypes first detected earlier (5-20 somites; Supplementary Fig. 9b-d) vs. later (25-34 somites; Supplementary Fig. 9e-f) in development. For example, we can identify distinct subsets of splanchnic mesoderm that are most highly related to foregut mesenchyme, hepatic mesenchyme and proepicardium, and may correspond to the ‘territories’ in which these organ-specific mesenchyme are induced (Supplementary Fig. 9b-d). These annotations may provide a starting point for deeper dives into each mesenchymal subset and how it patterns (and is reciprocally patterned by) the organ with which it interfaces, as has recently been done for the foregut^57^. For example, the hepatic and foregut mesenchyme are sharply distinguished from one another, as well as from the splanchnic territories from which they arise, by many genes including expected TFs (*e.g. Gata4* and *Barx1*, respectively^61, 62^). However, the inferred splanchnic territories of origin (labels 15 vs. 14 in Supplementary Fig. 9c) are also clearly distinct from one another, with hepatic mesenchymal progenitors expressing a program of epithelial-mesenchymal transition (EMT) and foregut mesenchymal progenitors expressing multiple guidance cue programs (e.g. semaphorins, ephrins, SLITs, netrins) (**Supplementary Table 13**).

### The Timing and Trajectories of Retinal Diversification

In the next two sections, we turn from the mesoderm to the neuroectoderm, first considering the retina and then neuronal diversification more broadly. In mouse development, neural epithelium arises as early as E9, giving rise to ciliary and pigment epithelium as well as multipotent retinal progenitor cells (RPCs) by E10. Through the remainder of fetal development and continuing postnatally, RPCs give rise to seven major types of retinal neurons in a conserved order^65^. Here we sought to leverage the depth and temporal resolution of these data to more precisely define the developmental intervals and rates at which retinal cell types emerge, proliferate and diversify.

We re-embedded and re-annotated 160,834 cells with relevant preliminary annotations across all timepoints (Fig. 4a; Supplementary Fig. 10a-b). We observe that eye field is already detectable in our earliest embryo (early head fold stage; 0 somite embryo in E8.5 bin; *Pax2*+, *n* = 782 cells), diversifying towards retinal progenitors (as early as E9.75) and retinal pigment epithelium (RPE) (as early as E10), as well as a third branch that appears as early as E9.5, sharply downregulates *Rax*, and ceases proliferating by E14.5, likely corresponding to the optic stalk (Supplementary Fig. 10c-d). This branch is undetectable in later time points, but pathway analysis suggests this is due to terminal differentiation in the context of a rapidly growing embryo, rather than apoptosis. Among retinal neurons, differentiation towards retinal ganglion cells (RGCs) begins as early as E11.75, and towards cone photoreceptors as early as E13. As development progresses, we observe a succession of retinal neuron types appearing in the expected order (Fig. 4b-c; Supplementary Fig. 10e), except for Müller glia, which emerge postnatally^66^. Among RPCs, the succession of sampled timepoints fills out a continuum of transcriptional states associated with diversification towards most major retinal neuron types (Fig. 4a-b)^67^. In contrast, the ciliary marginal zone (CMZ), identified as early as E11.25, remains most similar to a rapidly expanding pool of naive retinal progenitors. Strikingly, the CMZ appears to give rise to a second wave of pigment epithelium in the perinatal period, entirely separated in terms of both its transcriptional trajectory and timeframe from the branch leading to RPE, likely corresponding to the iris pigment epithelium (IPE; Fig. 4a; Supplementary Fig. 10f-g).

**Figure 4.**
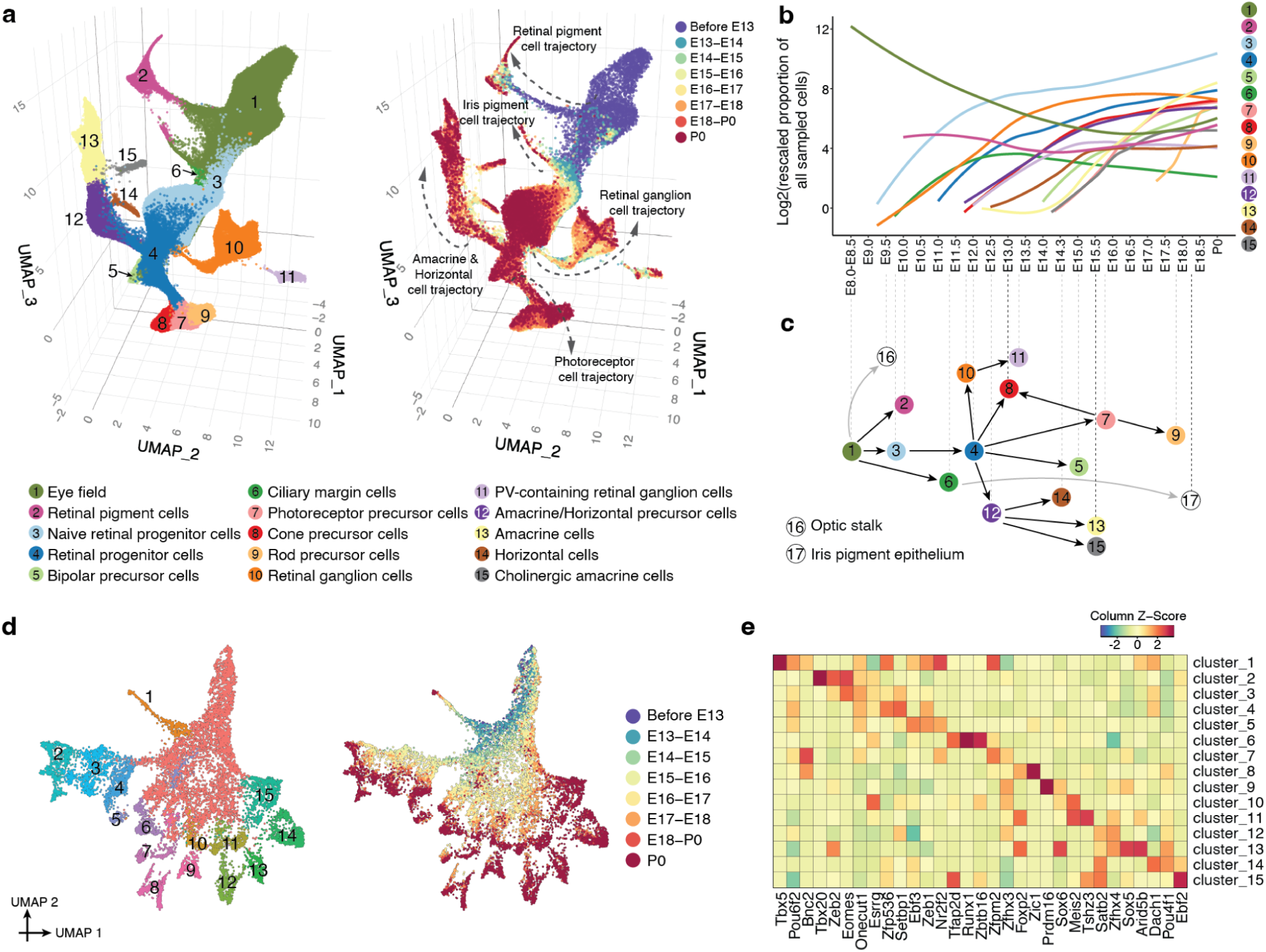
The timing and trajectories of retinal development. **a,** Re-embedded 3D UMAP of 160,834 cells corresponding to the retinal development from E8 to P0. Cells are colored by either their initial annotations (left) or timepoint (right, after downsampling to a uniform number of cells per time window). Arrows highlight five of the main trajectories observed. **b,** Rescaled proportion of profiled cells (log2; y-axis) for each cell type shown in panel a, as a function of developmental time (x-axis). For rescaling, the % of profiled cells in the entire embryo assigned a given annotation was multiplied by 100,000, prior to taking the log2. Line plotted with *geom_smooth* function in ggplot2. **c,** Schematic of retinal cell types emphasizing the timing at which they first appear and their inferred developmental relationships from E8-P0. The gray lines indicate subsets of the eye field and RPE subsequently annotated as the optic stalk (label 16) and iris pigment epithelium (label 17), respectively. Cell types are positioned along the x-axis at the timepoint at which they are first observed (Supplementary Fig. 10e). **d,** Re-embedded 2D UMAP of retinal ganglion cells. Cells are colored by either clusters (left; Leiden clustering followed by downselection to late-appearing clusters) or timepoint (right). **e,** The top 3 TF markers of the 15 clusters shown in panel d. Marker TFs were identified using the *FindAllMarkers* function of Seurat/v3^51^. Their mean gene expression values in each cluster are represented in the heatmap, calculated from original UMI counts normalized to total UMIs per cell, followed by natural-log transformation. The full list of significant TFs is provided in **Supplementary Table 14**.

Reanalyzing RGCs, we identify 15 clearly distinguishable subtypes, mainly diversifying in late gestation and well-defined by specific combinations of TFs (Fig. 4d-e). This extent of detected RGC diversity is on par with expectation for P0^68^, suggesting that the improved performance of sci-RNA-seq3 has substantially improved its ability to discriminate neuronal subtypes.

## The emergence of neuronal subtypes from the patterned neuroectoderm

The neuroectoderm and its derivatives comprise about 40% of cells profiled here (4.9M nuclei; Fig. 1e-f), of which the eye field and its derivatives are only about 3%. In this section, we describe the broader outlines of early neurogenesis in the post-gastrulation embryo, with deeper dives into the emergence of specific neuronal trajectories from the patterned neuroectoderm, as well as the transcriptional heterogeneity of spinal interneurons.

In the earliest embryos of this series (0-12 somite stage), we previously defined a continuum of cell states that correlated with anatomical patterning of the “pre-neurogenesis” neuroectoderm^11^. Performing a similar analysis here that extends through early organogenesis (E8-E13), we observe clusters corresponding to territories that will give rise to the major regions of the mammalian brain (Fig. 5a; Supplementary Fig. 11). As post-gastrulation development further unfolds, we observe numerous, distinct neurogenic trajectories arising from these territories (Fig. 5b; note that patterned neuroectoderm, direct neurogenesis and indirect neurogenesis largely correspond to the ‘neuroectoderm & glia’, ‘CNS neurons’ and ‘intermediate neuronal progenitors’ major cell cluster annotations, respectively)^73^. Of note, the extensive heterogeneity of the patterned neuroectoderm progressively diminishes as differentiating neurons become more similar with respect to their transcriptional states (Fig. 5c).

**Figure 5.**
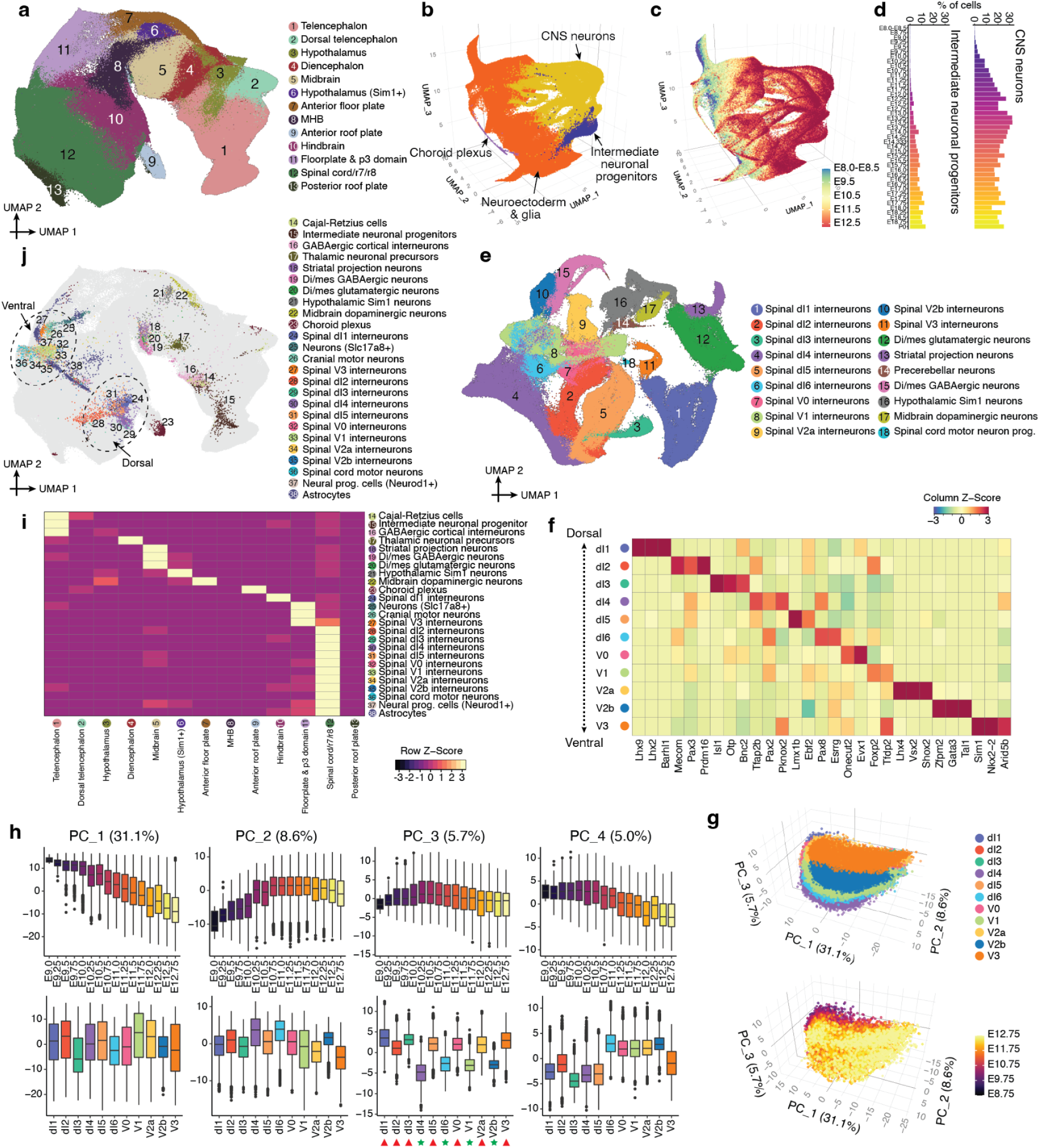
The emergence of neuronal subtypes from the patterned neuroectoderm. Please note panels are arranged in a clockwise manner. **a,** Re-embedded 2D UMAP of 1,185,052 cells, corresponding to different neuroectodermal territories, from neuroectoderm & glia major cell clusters from stages <E13. Derived cell types, *e.g.* eye field, glia, neurons, are excluded. **b,** Re-embedded 3D UMAP of 1,772,567 cells from neuroectodermal territories together with derived cell types, from stages <E13. Derived cell types are mainly from the CNS neuron, ependymal cell (more specifically, choroid plexus), and intermediate neuronal progenitor cell clusters. **c,** The same UMAP as in panel b, colored by timepoint. **d,** Composition of embryos from each 6-hr bin by intermediate neuronal progenitor (left) and CNS neuron (right) major cell clusters. **e,** Re-embedded 2D UMAP of 296,020 cells (glutamatergic neurons, GABAergic neurons, spinal cord dorsal progenitors, and spinal cord ventral progenitors) from stages <E13. **f,** The top 3 TF markers of the 11 spinal interneurons. Each spinal interneuron was first randomly downsampled to 10,000 cells and marker TFs were identified using the *FindAllMarkers* function of Seurat/v3^51^. Their mean gene expression values in each cluster are represented in the heatmap, calculated from original UMI counts normalized to total UMIs per cell, followed by natural-log transformation. The full list of significant TFs is provided in **Supplementary Table 15**. **g,** 3D visualization of the top three PCs of gene expression variation in 11 spinal interneurons (after randomly downsampling each spinal interneuron to 10,000 cells), calculated on the basis of the 2,500 most highly variable genes. Cells are colored by either cell types (top) or timepoints (bottom). **h,** Correlations between top four PCs and timepoints (top row) or cell types (bottom row). In the boxplots, the center lines show the medians; the box limits indicate the 25th and 75th percentiles. The red triangles and green stars highlight spinal cord interneurons which are glutamatergic vs. GABAergic identity, respectively. **i,** The number of mutual nearest neighbors (MNN) pairs between pairwise neuroectodermal territories (column) and their derivative cell types (row). The cell populations as shown in panel b, are first embedded into 30 dimensional PCA space, and then for individual derivative cell types, the MNN pairs (k = 10 used for k-NN) between their earliest 500 cells and the cells from neuroectodermal territories are identified. A handful of derivative cell types with <500 cells are excluded. Three derivative cell types with fewer than 50 MNN pairs are excluded but further analyzed by iterative mapping (Supplementary Fig. 14a). **j,** The same UMAP as in panel a, but with inferred progenitor cells colored by derivative cell type with the most frequent MNN pairs. Dotted circles highlight the dorsal and ventral spinal interneuron neurogenesis domains of the hindbrain & spinal cord.

Beginning as early as E8.75 (or more precisely, the 16 somite stage), most neuronal diversity in the prenatal mouse embryo is derived from direct neurogenesis (Fig. 5d), including motor neurons, cerebellar Purkinje cells, Cajal-Retzius cells, thalamic neuronal precursors and many other neuronal subtypes (see ‘CNS neurons’ sub-panel of Supplementary Fig. 4 for full list). Indirect neurogenesis^73^ has a slower start, with intermediate neuronal progenitors (*Eomes*+, *Pax6*+) first detected at E10.25, later giving rise to deep layer neurons, upper layer neurons, subplate neurons, and cortical interneurons (Fig. 5d; Supplementary Fig. 12a-b). Although many neuronal subtypes deriving from direct neurogenesis are easily distinguished, the majority of these cells (55%) could initially only be coarsely annotated as glutamatergic/GABAergic neurons or dorsal/ventral spinal cord progenitors (collectively 1.1M cells). To leverage the greater heterogeneity evident as these trajectories “launch” from the patterned neuroectoderm, we reanalyzed the subset of these cells derived from early embryos (stages earlier than E13), which greatly facilitated their more refined annotation (Fig. 5e; Supplementary Fig. 12c; **Supplementary Table 12**), while also highlighting the bases for components of this heterogeneity (*e.g.* anterior vs. posterior; excitatory vs. inhibitory; Supplementary Fig. 12d).

Among these more refined annotations are 11 subtypes of spinal interneurons. These assignments were based on subtype-specific TFs in these data (Fig. 5f; **Supplementary Table 15**) vs. the literature^74^. To systematically investigate transcriptional heterogeneity among spinal interneuron subtypes, we first applied the same PCA-based approach that we previously applied to NMPs (Fig. 2e-f). For differentiating spinal interneurons, PC1 and PC2 (together nearly 40% of variation) appear to correspond to neuronal differentiation (Fig. 5g-h; **Supplementary Table 16**), PC3 to glutamatergic vs. GABAergic identity, and PC4 to dorsal vs. ventral identity (Fig. 5h).

We next sought to infer the progenitor cells in the patterned neuroectoderm from which various neuronal and non-neuronal cell types derive. To this end, we took pre-E13 cells annotated as intermediate neuronal progenitors, astrocytes, choroid plexus, or any of the derivatives of direct neurogenesis, and co-embedded them with cells of the patterned neuroectoderm (Fig. 5a). Next, for each derivative cell type in this co-embedding, we selected 500 cells from the earliest stage embryos and identified their mutual nearest neighbor (MNN) cells in the patterned neuroectoderm. Finally, we located these inferred progenitors in our original embedding of the pre-E13 patterned neuroectoderm (Fig. 5i-j).

The resulting distribution of inferred progenitors in the patterned neuroectoderm is considerably more granular than our annotations of anatomical territories (Fig. 5j vs. Fig. 5a). The inferred progenitors of the choroid plexus are overwhelmingly in the anterior roof plate (91%), with a minor subset in the dorsal diencephalon (5%), although this balance is likely impacted by the temporal window in which this analysis is focused (E8-E13)^75^. Inferred astrocyte progenitors exhibit a more complex distribution, with inferred VA2 progenitors primarily assigned to the spinal cord/r7/r8 (83%) and hindbrain (16%), and inferred VA3 progenitors to the spinal cord/r7/r8 (57%) and floorplate & p3 domain (32%)^76^ (Supplementary Fig. 13). VA1 astrocytes appear to arise later than VA2 and VA3 astrocytes, and were not present in sufficient numbers for their progenitors to be inferred here.

The inferred progenitors of neuronal subtypes also largely fall within the expected territories, with additional sub-structure within those. For example, the inferred progenitors of dorsal and ventral spinal interneurons cluster distinctly (Fig. 5j). Of note, the progenitors of three neuronal subtypes (cerebellar Purkinje neurons, precerebellar neurons, spinal dI6 interneurons) were not clearly mappable. To address the possibility that this is because they share progenitors with other neuronal subtypes, we repeated the MNN analysis in an iterative fashion. Although the origins of precerebellar neurons and spinal dI6 interneurons remain ambiguous, this analysis suggests that cerebellar Purkinje neurons and dl2 spinal interneurons may have transcriptionally similar progenitors (Supplementary Fig. 14a).

How do neuronal subtypes’ identities arise, and how are they maintained^77^? As described above, we identified TFs that were specific to each of the 11 spinal interneuron subtypes (median 53 TFs per subtype; top 3: Fig. 5f; full list: **Supplementary Table 15**). These subtype-specific TFs exhibit diverse temporal dynamics (Supplementary Fig. 14b). Can we pinpoint which of these TFs might be involved in the initial specification of each subtype? Focusing on the dorsal spinal interneurons for which we successfully inferred progenitors (dl1-dl5), we identified TFs that are specific to the inferred progenitors of that subtype in the patterned neuroectoderm, relative to the inferred progenitors of other dorsal spinal interneurons (Supplementary Fig. 14c, top), most of which were bHLH or homeodomain TFs^78^. However, their expression was not maintained (Supplementary Fig. 14c, bottom), in line with the complex temporal dynamics of other subtype-specific TFs expressed after neuronal specification (Supplementary Fig. 14b)

Finally, we sought to systematically delineate the timing at which each of these neuronal subtypes initiates differentiation (Supplementary Fig.14d). This analysis suggests that the differentiation of each neuronal subtype from the patterned neuroectoderm is modestly asynchronous, but also clearly subtype-specific. For example, about 95% of inferred progenitors of dl2 spinal interneurons are from 20 somite to E11 stage embryos, while 95% of dl4 spinal interneurons inferred progenitors are from 27 somite to E11.75 stage embryos.

Taken together, these analyses are consistent with a model articulated by Sagner & Briscoe in which both spatial and temporal factors contribute to the specification of neuronal subtypes as they emerge from the patterned neuroectoderm^77^. Furthermore, they highlight the complexity of this process not only at the initiation of each neuronal subtype, but also over the course of their early maturation, *e.g.* at 6-hr resolution we can observe individual spinal interneuron subtypes expressing a dynamic succession of developmentally potent TFs (Supplementary Fig. 14b).

## A Rooted Tree of Cell Type Relationships Spanning E0 to P0

A primary objective of developmental biology is to delineate the lineage relationships among cell types. Transcriptional profiles of single cells do not explicitly contain lineage information. However, under the assumption that the transcriptional states of closely related cell types are more similar than those of distantly related cell types, one can envision a developmental tree based solely on scRNA-seq data^83^. Indeed, we and others have reconstructed scRNA-seq-based trees for portions of worm, fly, fish, frog and mouse development^1–7, 10^. Returning to the analogy of a time-lapse movie, the correspondence between the inferred and actual lineage relationships among cell types is anticipated to be a function of the quality and quantity of the snapshots taken (*e.g.* frame-rate of embryos sampled, number of cells sampled per time point, depth of sampling of each cell). As such, particularly for organogenesis & fetal development, the temporal resolution, breadth and depth of data reported here creates an opportunity to refine and extend a tree that relates cell types throughout mouse development.

In this section, we summarize our efforts to construct a rooted tree of cell types that spans mouse development from zygote to pup, *i.e.* E0 to P0, based on four published datasets^6, 84–86^ (110,000 cells spanning E0 to E8.5) and the main dataset described here (11.4M cells spanning E8 to P0) (**Supplementary Table 17**). As with our initial attempt at reconstructing such a tree for mouse development^11^, a major challenge is that these datasets are based on different scRNA-seq profiling technologies. Further challenges include the fact that cells’ transcriptional states are only loosely synchronized with developmental time^1–7, 10^, the multiple scenarios by which cell state manifolds may be misleading^83^, and finally, the sheer complexity of this mammalian organism.

To address these challenges, we adopted a heuristic approach, in which the tree reconstruction process was primarily data-driven, but with some degree of manual curation applied. First, based on data source, developmental window and cell type annotations, we split cells into fourteen subsystems which could be separately analyzed and subsequently integrated. The first two subsystems correspond to the pre-gastrulation and gastrulation phases of development and are based on external datasets^6, 84–86^. The remaining twelve subsystems derive from the data reported here, and collectively encompass organogenesis & fetal development (**Supplementary Tables 17-18**).

Second, dimensionality reduction was performed separately on cells from each of the fourteen subsystems. Manual reexamination of each subsystem led to some corrections or refinements of cell type annotations, ultimately resulting in 283 annotated cell type nodes, some with only a handful of cells (*e.g.* 60 ciliated nodal cells) and others with vastly more (*e.g.* 650,000 fibroblasts) (**Supplementary Table 19-20**). Of note, each of these annotated cell type nodes derives from one data source, such that there are some redundant annotations that facilitate “bridging” between datasets (Supplementary Fig. 15). In contrast with our previous strategy in which nodes were stage-specific^11^, each cell type node here is temporally asynchronous, and of course may also contain other kinds of heterogeneity (*e.g.* spatial, differentiation, cell cycle, etc.).

Third, we sought to draw edges between nodes (Fig. 6a-f). Within each subsystem, we identified pairs of cells that were mutual nearest neighbors (MNN) in 30-dimensional PCA space (k = 10 neighbors for pre-gastrulation and gastrulation subsystems, k = 15 for organogenesis & fetal development subsystems). Although the overwhelming majority of MNNs occurred within cell type nodes, some MNNs spanned nodes and are presumably enriched for *bona fide* cell type transitions. To approach this systematically, we calculated the total number of MNNs that spanned each possible pair of cell type nodes within a given subsystem, normalized by the total number of possible MNNs between those nodes, and ranked all possible intra-subsystem edges based on this metric (**Supplementary Table 21**). Of note, due to its complexity, this was done in two stages for the “Neuroectoderm & glia” subsystem, first applying the heuristic to the subset of cell types corresponding to the patterned neuroectoderm, and then again to identify edges between the patterned neuroectoderm and its derivatives (*i.e.* neurons, glial cells, etc.).

**Figure 6.**
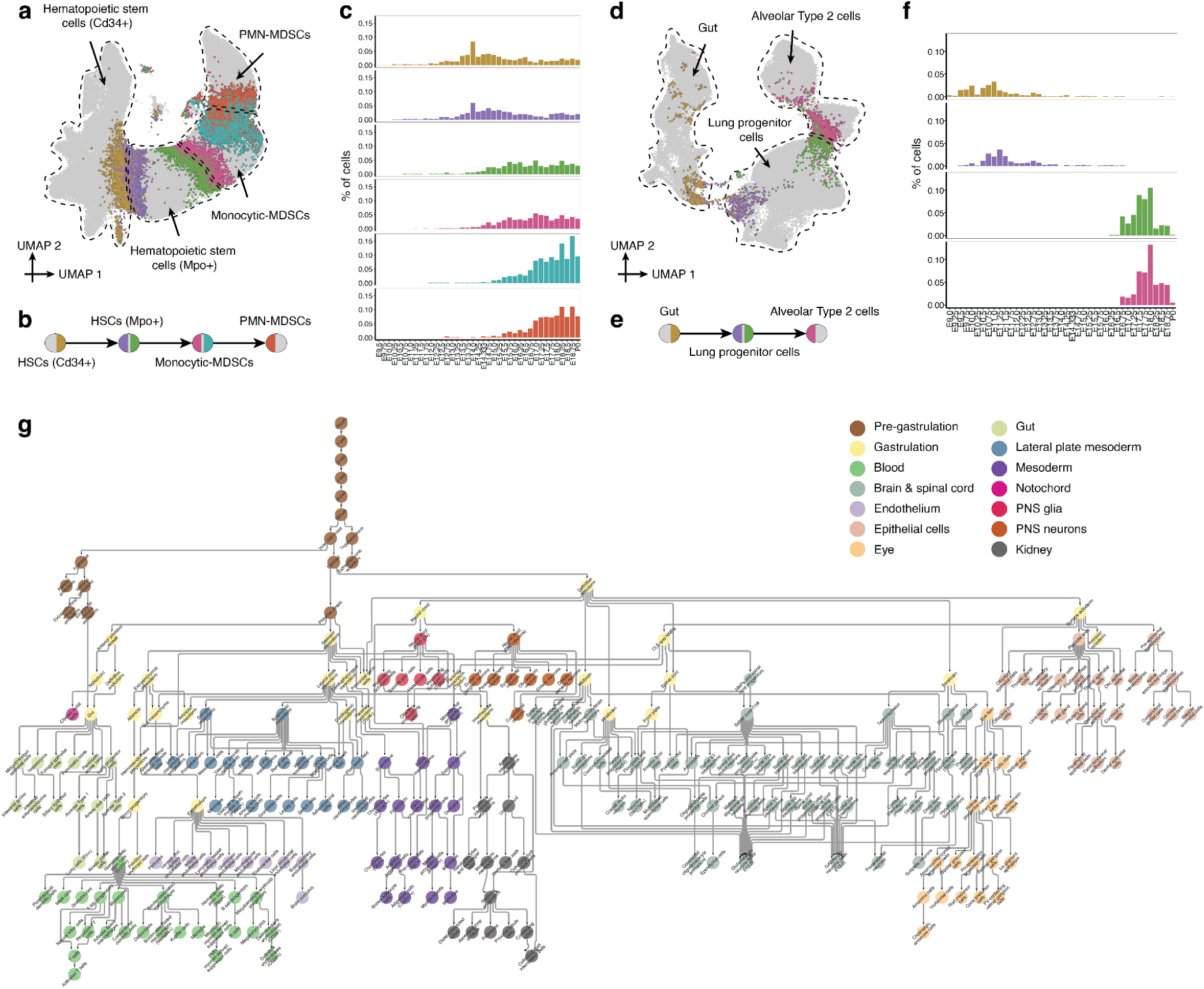
A data-driven tree relating cell types throughout mouse development, from zygote to pup. **a,** Illustration of basis for edge inference heuristic. Re-embedded 2D UMAP of 101,001 cells from hematopoietic stem cells (Cd34+), hematopoietic stem cells (Mpo+), monocytic myeloid-derived suppressor cells (MDSCs), and PMN myeloid-derived suppressor cells (MDSCs) within the “Blood” subsystem. Cells involved in MNN pairs that bridge cell types are coloured. For example, hematopoietic stem cells (Cd34+) and hematopoietic stem cells (Mpo+) involved in MNN pairs between these two cell types are coloured yellow and light purple, respectively. **b,** Inferred lineage relationships between annotated cell types in panel a, with corresponding color scheme. **c,** The % of inter-cell-type MNN cells (y-axis) over the total number of cells profiled from embryos from the corresponding time bin, with the same color scheme as panels a & b. **d,** Additional illustration of basis for edge inference heuristic. Re-embedded 2D UMAP of 71,718 cells from gut, lung progenitor cells, and alveolar Type 2 cells within the “Gut” subsystem. Cells involved in MNN pairs that bridge cell types are coloured. For example, gut and lung progenitor cells involved in MNN pairs between these two cell types are coloured yellow and light purple, respectively. **e,** Inferred lineage relationships between annotated cell types in panel d, with corresponding color scheme. **f,** The % of inter-cell-type MNN cells (y-axis) over the total number of cells profiled from embryos from the corresponding time bin, with the same color scheme as panels d & e. **g,** A rooted, directed graph corresponding to a mouse development, spanning E0 to P0 (yFiles Hiearchic layout in *Cytoscape*/v3.9.1). For presentation purposes, most ‘spatial continuity’ edges are removed (except for those between spinal cord dorsal & ventral progenitors (after E13.0) and GABAergic & glutamatergic neurons (after E13.0)). We also merged nodes with redundant labels derived from different datasets (i.e. “dataset equivalence” edges), resulting in a rooted graph composed of 262 cell type nodes and 338 edges. Nodes are colored and labeled by each of the 14 subsystems: Pre-gastrulation (*n* = 2,250 cells), Gastrulation (*n* = 108,857 cells), Blood (*n* = 1,576,789 cells), Brain & spinal cord (*n* = 4,434,234 cells), Endothelium (*n* = 312,029 cells), Epithelial cells (*n* = 398,373 cells), Eye (*n* = 166,852 cells), Gut (*n* = 453158 cells), Lateral plate mesoderm (*n* = 992,903 cells), Mesoderm (*n* = 2,717,903 cells), Notochord (*n* = 3,812 cells), PNS glia (*n* = 126,743 cells), PNS neurons (*n* = 146,825 cells), and Kidney (*n* = 111,786 cells).

Fourth, we manually reviewed the ranked list of 1,155 candidate edges for biological plausibility (those with a normalized MNN score > 1; Supplementary Fig. 15a), resulting in 452 edges which we manually annotated as more likely to correspond to either “developmental progression” or “spatial continuity” (**Supplementary Table 22**). Where nodes were connected to more than one other node, distinct subsets of cells were generally involved in each edge (Fig. 6a-b; Fig. 6d-e), and internode MNN pairs exhibited temporal coincidence (Fig. 6c,f). As only a handful of cells were profiled in the pre-gastrulation subsystem, those edges were added manually.

Finally, to bridge subsystems, we performed batch correction and co-embedding of selected timepoints from either the pre-gastrulation and gastrulation datasets, or the gastrulation and organogenesis & fetal development datasets, to identify equivalent cell type nodes, resulting in a third category of “dataset equivalence” edges (Supplementary Fig. 15b-e). Most of the 12 organogenesis & fetal development subsystems originate in cell type nodes for which equivalent nodes are already present at gastrulation (Supplementary Fig. 15e). The exceptions, presumably due to undersampling of this transition, were the “blood” and “PNS neuron” subsystems, for which we manually added edges to connect them with biologically plausible pseudo-ancestors. Altogether, we added 55 inter-subsystem edges.

The resulting developmental cell type tree, spanning E0 to P0 or zygote-to-pup, can be represented as a rooted, directed graph (Fig. 6g), in which dataset equivalence edges are oriented forward in time, developmental progression edges are manually oriented, and spatial continuity edges are left bidirectional. In practice, a small number of nodes in the tree have more than one parent, so the “tree” is formally a rooted, directed graph.

## Systematic Nomination of Transcription Factors and Other Genes for Cell Type Specification

We previously annotated the edges of a more limited cell type tree of mouse development by identifying TFs exhibiting sharp changes in gene expression at the developmental timepoint where a given cell type first appears, through comparisons of all cells of a new type to all cells of its inferred pseudoancestor as well as to any sister nodes^11^. Here, in the course of similarly annotating the zygote-to-pup cell type tree shown in Fig. 6g, we sought to take a more nuanced approach.

In particular, we stratified each cell type transition into four phases (Supplementary Fig. 16a). Given a directional edge between two nodes, A→B, we identified the subset of cells within each node that were either “inter-node” MNNs of the other cell type (groups 2 & 3 in Supplementary Fig. 16b) or “intra-node” MNNs of those cells (groups 1 & 4 in Supplementary Fig. 16b). If A→B, this approach effectively models the transition as group 1→2→3→4. Next, we identified differentially expressed TFs (DETFs) and genes (DEGs) across each portion of the modeled transition, *i.e.* early (1→2), inter-node (2→3), and late (3→4). In contrast with our previous heuristic^11^, this strategy highlights differences between cells that are most proximate to the cell type transition itself. Moreover, DETFs and DEGs identified in the early (1→2) and late (3→4) phases correspond to changes *within* node A or B, respectively, rather than between nodes A and B, which may facilitate the identification of early changes in gene expression, *i.e.* those that precede the cell type transition itself.

We applied this heuristic to 436 edges of the rooted tree shown in Fig. 6g, excluding only dataset equivalence edges and the pre-gastrulation subsystem. Of note, the directionality of many of these edges was not immediately obvious (*i.e.* those annotated as “spatial continuity” edges in **Supplementary Table 22**). In these cases the orientation of the “early” and “late” phases is arbitrary. For edges with a relatively small number of MNN pairs, we expanded each group to at least 200 cells by iteratively including their MNNs within the same cell type, to increase statistical power. Across all 436 edges, this heuristic resulted in the nomination of ranked lists of median 28 (IQR 12-51)DETFs and median 171 (IQR 76-389) DEGs per edge (mean), 5% (DETFs) and 7% (DEGs) of which were exclusively nominated in either the early or late phase of the transition, respectively. The most significant DETFs and DEGs for each edge are provided in **Supplementary Tables 23** and **24**, respectively. Most TFs and genes are only nominated in the context of one or a few edges, but there are a number of outliers that presumably have more general roles in cell type specification (Supplementary Fig. 16c-d).

On a cursory review, there are many instances in which the top-ranked upregulated DETF for the early phase of the transition corresponds to a well-established, early regulator of that cell type (*e.g. MITF* for melanocytes^87^; *Ebf1* & *Pax5* for B-cell progenitors^88^; *Lef1* for B-cells^89^; *Zfpm1* for megakaryocyte-erythroid progenitors^90^), but also many nominations of potentially novel regulatory relationships that may warrant further investigation (*e.g. Ltf* for monocytic myeloid-derived suppressor cells; *Tcf7l2* for Kupffer cells, *Esrrg* for dorsal telencephalon-derived choroid plexus; *Zfp536* for myelinating Schwann cells; *Rreb1* for adipocyte progenitors, etc.) (**Supplementary Table 23**).

Digging further into a well-studied transition, *Sox17* is the sole upregulated DETF during the early phase of the transition from anterior primitive streak to definitive endoderm (group 1→2), while a broader set of TFs (*Elf3, Sall4*, *Hesx1*, *Lin28a*, *Hmga1*, *Ovol2*, but notably not *Sox17*) are upregulated during the transition itself (group 2→3) (**Supplementary Table 23**). Non-TF DEGs specific to the early phase of the transition include *Cer1* (encoding Cerberus, a well-established signaling molecule and marker of definitive endoderm)^91^, *Slc25a4* (encoding an adenine nucleotide translocator specific to the inner mitochondrial membrane)^92^, and *Slc2a3* (encoding Glut3, a glucose transporter specific to cell types with high energy needs)^93^ (**Supplementary Table 24**). To examine this further, we extracted all cells participating in groups 1-4 of this transition and subjected them to conventional pseudotime analysis^7^. In brief, this analysis supported the upregulation of *Sox17* as preceding that of other nominated key TFs, and further highlighted *Cer1* as the only non-TF DEG with similar kinetics to *Sox17* (Supplementary Fig. 16e-f).

A more complex example involves hematopoietic stem cells (Cd34+), which in the developmental graph are the node-of-origin origin of a dozen cell types (Supplementary Fig. 16g). An important point is that although we treat hematopoietic stem cells (HSC) (Cd34+) as a single node for the purposes of the zygote-to-put developmental graph, the cells assigned to the node (as with most other nodes) are quite heterogeneous. For example, the cells participating in the MNNs that support the edges to various lymphoid, myeloid and erythroid derivatives are distinct subpopulations of HSCs (Supplementary Fig. 16h-i), and these subpopulations differentially express TFs nominated as early regulators, *e.g. Ebf1* for B cells^88^ and *Id2*^94^ and *Nfatc2* for conventional dendritic cells (Supplementary Fig. 16j). Establishing a level of resolution for a developmental tree that hits the right balance between the complexity of mammalian development and the desire for interpretability is an ongoing tension.

## Rapid Shifts in Transcriptional State Occur in a Restricted Subset of Cell Types Upon Birth

In the course of our analyses of this time-lapse, we anecdotally noted that for certain cell types, cells derived from P0 pups appeared very well separated from their fetal pseudoancestors, in sharp contrast with other cell types across the same temporal interval as well as with even these same cell types at all prior temporal intervals. The proximal tubule is one example of this phenomenon, discussed briefly above (cluster 9 in Fig. 3a-b; Supplementary Fig. 7d). However, a similar pattern was also noted for hepatocytes, adipocytes, and various cell types of the lungs and airways (Fig. 7a). As we worried that this could be due to batch effects or the well-documented pitfalls of over-interpreting UMAPs^95^, we conducted a time point correlation analysis. In brief, for each cell type, we performed dimensionality reduction (30 dimensions) on cells of that type from multiple, late-gestational timepoints (E16 to P0). We then identified k-nearest neighbors (k-NN, k = 10) to ask whether neighboring cells were derived from the same or different timepoints. In this framing, a low proportion of neighbors from different timepoints corresponds to a relatively abrupt change in transcriptional state. In general, cells from P0 pups were closer neighbors to other P0 cells than was the case for all other timepoints (Fig. 7b), which might be explained in part by a longer temporal interval between E18.75 and P0 than 6 hours. However, for many cell types, the difference was extreme, consistent with dramatic changes in a transcriptional state upon birth. These include not only the aforementioned cell types, but also various endothelial and blood lineages (Fig. 7b). In contrast, neuronal cell types did not exhibit such marked differences between animals collected shortly before vs. after birth (Fig. 7b).

**Figure 7.**
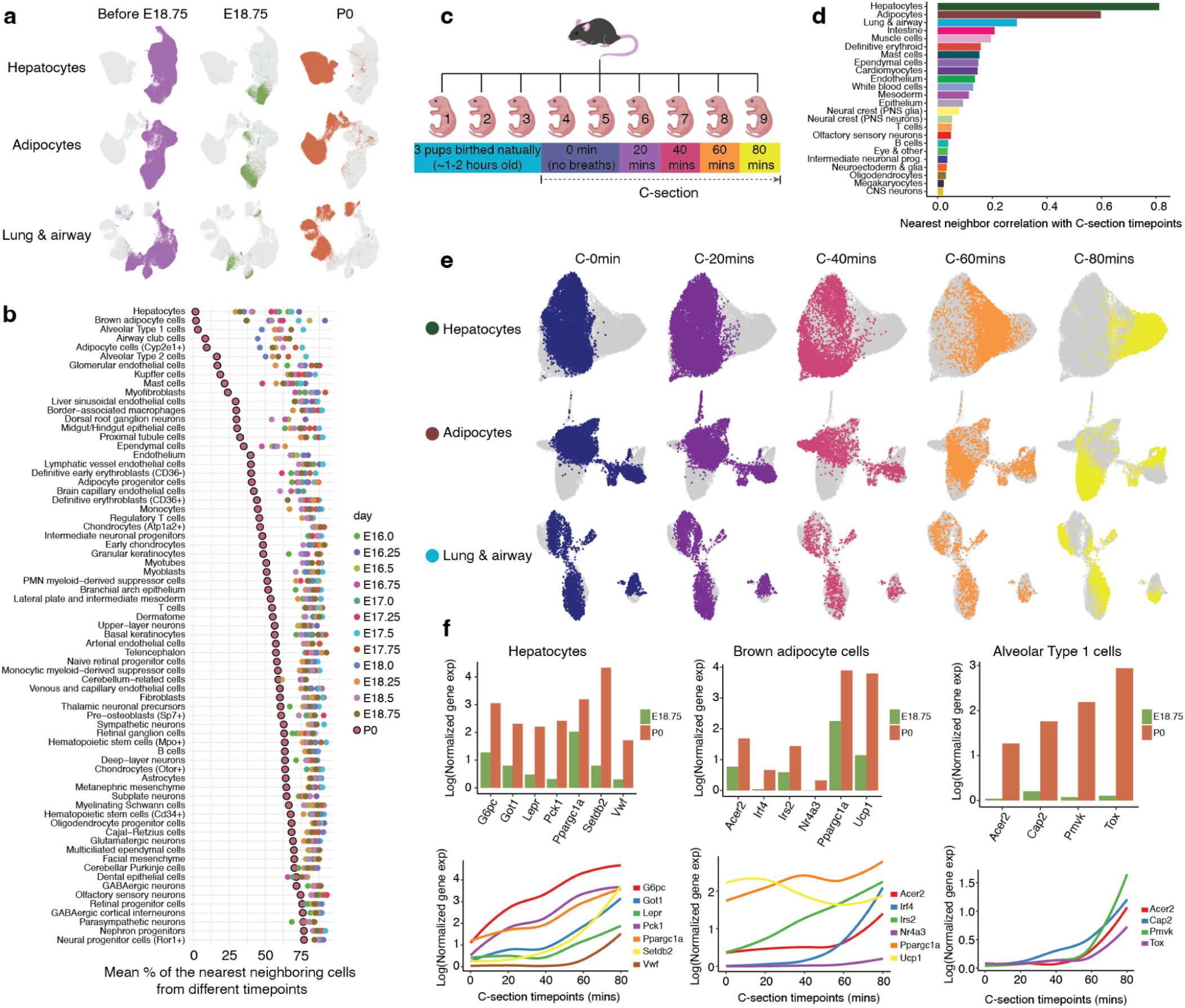
Rapid shifts in transcriptional state occur in a restricted subset of cell types upon birth. **a,** Re-embedded 2D UMAP of cells from three major cell clusters: hepatocytes (top row), adipocytes (middle row), and lung & airway (bottom row). For each row, the same UMAP is shown three times, with colors highlighting cells from before E18.75 (left), E18.75 (middle), or P0 (right) embryos. **b,** To systematically identify which cell types exhibit abrupt transcriptional changes before vs. after birth, for each of the 190 cell types, cells from animals harvested subsequent to E16 were combined, followed by PCA based on the top 2,500 highly variable genes. The 71 cell types with >= 200 cells from P0 and >= 200 cells from at least five timepoints prior to P0 were retained for the following analysis. First, timepoints with >= 200 cells were selected followed by downsampling cells from each timepoint to the median number of cells across those selected timepoints. Then, individual cells were searched for *k*-nearest neighbors (*k* was adjusted for different cell types, by taking log2-scaled median number of cells across the selected timepoints) in PCA space (*n* = 30 dimensions). Finally, for cells within each cell type, the average proportions of their nearest neighbor cells that are from a different timepoint were calculated. In this framing, a low proportion of neighbors from different timepoints corresponds to a relatively abrupt change in transcriptional state. Points corresponding to P0 are highlighted with black boundary. **c,** A new scRNA-seq dataset (hereafter referred to as “birth-series”) was generated from nuclei derived from nine individual pups from a single litter harvested shortly after delivery. This includes three vaginally birthed pups and six C-section delivered pups with 20 min increments between delivery and harvesting. **d,** For each major cell cluster in the birth-series dataset, we took cells from the six pups delivered by C-section and calculated a Pearson correlation coefficient between the timepoint of each cell and the average timepoints of its 10 nearest neighbors identified from the global PCA embedding (*n* = 30 dimensions). In this framing, high correlations are consistent with rapid, synchronized changes in transcriptional state. **e,** Re-embedded 2D UMAP of cells from three major cell clusters, based on cells from six pups delivered by C-section in birth series dataset: hepatocytes (*n* = 41,122 cells, top row), adipocytes (*n* = 19,696 cells, middle row), and lung & airway (*n* = 7,986 cells, bottom row). For each row, the same UMAP is shown multiple times, with colors highlighting cells from pups harvested after different intervals after delivery. **f,** Average normalized gene expression of selected genes are plotted between E18.75 vs. P0 in the original data (top), and normalized gene expression of the same genes are plotted as a function of C-section timepoints (bottom), for hepatocytes (left column), brown adipocyte cells (middle column), and alveolar type 1 cells (right column), respectively. Gene expression values were calculated from original UMI counts normalized to total UMIs per cell, followed by natural-log transformation. The line of gene expression was plotted by the *geom_smooth* function in ggplot2.

To validate and further resolve periparturition transcriptional dynamics, we collected nine pups that were part of a single litter. After 3 pups were delivered vaginally, the remaining six pups were delivered by Cesarean section (C-section) and sacrificed either immediately (0 min; 2 pups), or 20, 40, 60 or 80 min after C-section (1 pup each) (Fig. 7c; Supplementary Fig. 17a). Nuclei from these nine pups, together with residual nuclei from three samples from the original experiments, were subjected to a new sci-RNA-seq3 experiment, which yielded nearly 1M additional single cell profiles (Supplementary Fig. 17b; **Supplementary Table 1-2**).

Focusing first on the six pups delivered by C-section and 24 major cell clusters for which we were reasonably powered, we applied timepoint correlation analysis, as described above except treating the C-section timepoint as a continuous variable. Consistent with our earlier results (Fig. 7b), the outliers are the hepatocyte, adipocyte and lung & airway major cell clusters (Fig. 7d; Supplementary Fig. 17c-d). This experiment replicates our initial finding and also narrows the window in which these abrupt changes are first evident to the first hour of extrauterine life (Fig. 7e).

It is plausible that rapid changes in transcriptional programs might be physiologically necessary due to the profound differences between the placental and extrauterine environments. Moreover, one can imagine why these might be triggered by birth rather than developmentally timed, because of uncertainty with respect to precisely when birth will occur. Finally, we speculate that the proper development of certain organs might require the continuation of a precisely timed program (*e.g.* the brain), whereas for other organs, survival depends on an extremely rapid change in their function in the immediate wake of birth (*e.g.* the lungs & airways) (Fig. 7b). In examining DEGs of rapidly changing cell types, either between E18.75 and P0 embryos or across the 20 minute time-series of pups delivered by C-section, we see clues in regards to at least some of the specific physiological functionalities that these rapid changes might be serving (**Supplementary Tables 25-26**).

For example, in hepatocytes, genes involved in gluconeogenesis are sharply upregulated, including *Ppargc1a,* which encodes Pgc-1α, which in the liver serves as a master regulator of hepatic gluconeogenesis^96^, as well as *Pck1*, *G6Pc* and *Got1*, which encode key enzymes in this pathway (Fig. 7f). Aspects of these changes have previously been linked to changes in key nutritional hormones (rising glucagon, falling insulin) immediately after birth and presumably are necessary for maintaining normoglycemia in the wake of being abruptly cut off from maternal nutrients^97^. In brown adipocytes, we observe sharp upregulation of *Irf4*, a cold-induced master regulator of thermogenesis, and again of *Ppargc1a*, which in adipocytes plays a different role than in the liver, as Pgc-1α partners with Irf4 to drive the expression of *Ucp1* and uncoupled respiration^98^, presumably to maintain body temperature upon transition to the extrauterine environment^99^ (Fig. 7f).

The exact amount of time that lapsed between the birth of the vaginally birthed pups and their harvesting was not precisely captured in the replication experiments. However, on co-embedding cells derived from vaginally birthed pups with those delivered by C-section for the three most relevant major cell clusters, timepoint correlation analysis suggested that they were harvested within an hour of their birth (Supplementary Fig. 17e). However, this assumes similar kinetics for these rapid transcriptional changes in C-section vs. vaginally delivered pups. On more detailed inspection, the patterns are considerably more complex, with certain clusters (most notably, a subset of hepatocytes) appearing specific to vaginally birthed pups (Supplementary Fig. 17f; **Supplementary Table 27**).

What might explain this? The transition from the placenta to extrauterine life, as experienced by the neonate (and perhaps even by specific organs within the neonate), is very different between C-section vs. vaginal delivery, and various studies have shown that human babies delivered by each route have differences in physiology and health outcomes^100^. It is plausible that aspects of these postnatal phenotypic differences have their roots in how the massive, abrupt, cell type-specific changes documented here are influenced by mode of delivery.

## Discussion

We present a rich dataset in which we profiled the transcriptional states of 12.4 million nuclei sampled from 83 precisely staged embryos spanning late gastrulation (E8) to birth (P0), with 2-hr temporal resolution during somitogenesis, 6-hr resolution through to birth, and 20-min resolution during the immediately postpartum period. We note that despite its large scale, the entire project was driven by a small number of individuals. For example, the mouse embryos were staged by a single individual (IW), the vast majority of the sci-RNA-seq3 experiments were carried out by a single individual (BM) and the computational analyses were conducted by a single individual (CQ), with nearly all experiments and analyses completed within one year. We estimate that the direct costs of all reagents and labor invested in processing frozen embryos to sequencing libraries totaled to about $70,000, while the cost of sequencing totaled to about $300,000. Notably, our single dataset is equivalent to about 30% of the aggregated corpus of the Human Cell Atlas Data Portal as of March 2023.^101^

Three broad concepts supported our ability to generate, analyze and integrate such a large dataset with a small team at a modest cost: First, *multiplexing,* which fundamentally underlies the exponential scalability of single cell combinatorial indexing as well as that of massively parallel DNA sequencing. Second, *open science,* as we have taken abundant advantage of many freely released software packages for single cell data analysis released by the community^7, 18, 51, 60^. Third, our focus on *mouse development*, an eminently reproducible process through which we could access all mammalian cell types (or their predecessors) within a series of physically compact samples.

Our main goal in initiating this study was not to learn a specific piece of biology, but rather to lay a foundation for a comprehensive understanding of the development of a mammalian gastrula into a free-living pup (E8 → P0). By annotating hundreds of cell types and conducting deeper dives into selected developmental systems, we have sought to illustrate the early value of this foundation. Although the dataset is a rich source of hypotheses (*e.g.* the nomination of specific TFs as *in vivo* regulators of myriad cell types), the largest surprise to us was the discovery of rapid changes in transcriptional state in a restricted subset of cell types in the minutes to hours following birth. There is immense evolutionary pressure on the transition from placental to extrauterine life, which is arguably as fraught a moment as gastrulation in terms of physiological peril^102^. In specific cases, the genes sharply upregulated in specific cell types can be matched to specific adaptations (*e.g.* gluconeogenesis in hepatocytes, uncoupled respiration and thermogenesis in brown adipocytes). On the other hand, dozens of additional genes are sharply upregulated or downregulated in hepatic, adipose, and lung & airway tissues immediately after birth (**Supplementary Tables 25-26**). For these and many other tissues and cell types in which rapid periparturitional transcriptional changes are also documented (Fig. 7b,d), the adaptive functions served, not to mention the mechanisms underlying their rapid induction as well as differences shaped by the circumstances of delivery, warrant further exploration.

We recently proposed the concept of a “consensus ontogeny” of cell types, inclusive of both lineage histories and molecular states, as a potential structure for a “reference cell tree”103. The cell type tree constructed here, which spans mouse development, from a single cell zygote to the half-billion cells that make up a free-living pup (E0 → P0), represents a step in that direction. As with previous such steps by us and others^6, 7, 10, 11^, both cell type annotations and the tree structure itself almost certainly contain errors and are subject to corrections and refinement. Furthermore, as additional methods for single cell molecular profiling (*e.g.* chromatin accessibility, spatial mapping, etc) or organism-scale lineage tracing^104–108^ are applied to the mouse, we envision that this cell type tree (or other reductionist representations of development) will provide a framework for an increasingly rich, navigable roadmap of mouse development. Because it touches all aspects of prenatal development of the embryo proper from gastrulation to birth, the dataset may also be useful in ways that we did not anticipate at the project’s outset, *e.g.* as input for pre-training large language models of mammalian biology. Finally, just as Sulston reconstructed both the embryonic and post-embryonic lineages^109, 110^, we note that mouse development does not end at P0. We envision that the methods described here can also be applied to postnatal timepoints at the scale of the entire organism, with the long term goal of generating a time-lapse of the entire life cycle of a mammal at single cell resolution, from a single cell zygote to a natural death (E0 → D0).

## Methods

### Data reporting

For newly generated mouse embryo data, no statistical methods were used to predetermine sample size. Embryo collection and sci-RNA-seq3 data generation were performed by different researchers in different locations.

### Mouse embryos collection and staging

The details of collecting 12 mouse embryos around E8.5 with somites ranging from 0 to 12 were described in a previous publication^11^. Briefly, C57BL/6NJ (strain# 005304) mice were obtained at The Jackson Laboratory and mice were maintained via standard husbandry procedures. Timed matings were set in the afternoon and plugs were checked the following morning. Noon of the day a plug was found was defined as embryonic day (E) E0.5. On the morning of E8.5, individual decidua were removed and placed in ice cold PBS during the harvest. Individual embryos were dissected free of extraembryonic membranes, imaged, and the number of somites present were noted prior to snap freezing in liquid nitrogen (Supplementary Fig. 1). A portion of yolk sac from each embryo was collected for sex based genotyping and samples were stored at -80C until further processing.

For the newly generated mouse embryo data (ranging from E8.75 to P0), we employed a combination of staging methodologies depending on gestational age of harvest (Supplementary Fig. 2). To maximize temporal coherence, resolution, and accuracy, we sought to stage individual embryos based on well-defined morphological criteria, rather than by gestational day alone. Embryos harvested between E8.0 - E10.0 were staged based upon the number of somites counted at the time of harvest and further characterized by morphological features (Supplementary Fig. 1). For E10.25-E14.75 embryos, developmental age was determined using the embryonic Mouse Ontogenetic Staging System (eMOSS, https://limbstaging.embl.es/), which leverages dynamic changes in hindlimb bud morphology and landmark-free based morphometry to estimate the absolute developmental stage of a sample^15, 16^. A modified staging tool, implemented in python, with increased performance on E14.0-E15.0 samples was used to confirm staging of samples within this window (documentation and python scripts available at: https://github.com/marcomusy/welsh_embryo_stager). To distinguish samples staged via eMOSS, these samples are designated with “mE” for morphometric embryonic day (*e.g.* mE13.5, Supplementary Fig. 2). Due to the increased complexity of limb morphology at later stages automated staging beyond ∼E15.0 is not possible. As a consequence, harvests for all remaining embryonic samples (E15.0-E18.75) were performed precisely at 00:00, 06:00, 12:00, and 18:00 on the targeted day. From close inspection of limbs in this sample set we defined additional dynamics related to digit morphogenesis that allowed further binning of samples collected on Days 15 and 16 (Supplementary Fig. 2). Therefore, amongst samples profiled in this study only the E17.0-E18.75 samples were staged solely by gestational age. Lastly, P0 samples were harvested from litters at noon of the day of birth (parturition for C57BL/6NJ occurs between E18.75 and E19.0).

### Collecting mouse pups immediately after birth

Samples for analysis of periparturition transcriptional dynamics were collected from a plugged female that was monitored for signs of labor beginning at E18.75. Following the natural delivery of 3 pups the dam was euthanized and following removal from the uterus and extraembryonic membranes the remaining pups were either harvested immediately or placed in a warming chamber to monitor respiratory response and collected at 20 minute intervals. We collected nine new pups, to perform another sci-RNA-seq3 experiment. The first three pups are estimated to be between 1-2 hours old, although this was not precisely timed (samples 1-3 in Fig. 7c-d). None of these pups had nursed at the time of harvest. The next two pups were taken by C-section, decapitated and snap frozen immediately; no breaths were taken (samples 4-5 in Fig. 7c-d). The next four pups were taken by C-section and used for a “pink up” time course, harvesting one pup every 20 min (*i.e.* 20 mins, 40 mins, 60 mins, and 80 mins; samples 6-9 in Fig. 7c-d). During this time, all pups remained very active and working to establish a breathing rhythm. Pup 6 had not fully pinked up at time of harvest, but pups 7-9 had. Pups 8 and 9 had visible lungs in their chest cavities at 60 min. The last pup harvested at 80 min was fully pink with a reasonably stable breathing rhythm. No vocalization was heard from any pups during this collection. Of note, for additional quality control, we put nuclei from previously profiled E18.75 and P0 embryos into a small number of wells of the sci-RNA-seq3 experiment in which nuclei from this series were processed.

### Generating data using an optimized version of sci-RNA-seq3

Together with E8.5 data which has been reported before^11^, a total of 15 sci-RNA-seq3 experiments were performed on 75 individual mouse embryos. At least one sample was included for every ∼6 hour time interval from E8.0 to P0, and we also included a fine-scale sample set of embryos with distinct somite counts from 0-34 somites. Multiple samples were selected for a few timepoints (*e.g.* two samples for E13.0) to boost cell numbers. Meanwhile, we tried to ensure that both male and female mice roughly appear at adjacent timepoints (Figure. 1d). A detailed summary & images of individual embryos can be found in Supplementary Fig. 1-2 and **Supplementary Table. 1**.

To generate the dataset, we used the optimized sci-RNA-seq3 protocol^111^ as written, adjusting the volume and type of lysis buffer to the size of the embryos. Briefly, frozen embryos were pulverized on dry ice and cells were lysed with a phosphate-based, hypotonic lysis buffer containing magnesium chloride, Igepal, diethyl pyrocarbonate (DEPC) as an RNase inhibitor, and either sucrose or bovine serum albumin (BSA). Lysate was passed over a 20um filter, and the nuclei-containing flow-through was fixed with a mixture of methanol and dithiobis (succinimidyl propionate) (DSP). Nuclei were rehydrated and washed in a sucrose/PBS/triton/magnesium chloride buffer (SPBSTM), then counted and distributed into 96-well plates for reverse transcription with indexed oligo-dT primers.

Age-specific adaptations were as follows:

E10-CE13 embryos use 5mL BSA lysis buffer, E14 embryos use 10mL BSA lysis buffer, E15-E18 embryos use 20mL sucrose-based lysis buffer. Each of these samples were split over 48-96 wells for reverse transcription and the first round of indexing. A newborn P0 mouse requires 40mL of sucrose-based lysis buffer, and the lysate is divided into 4 fractions for filtration and fixing because of the amount of tissue involved. The two P0 mice were each processed as an individual experiment and were each split over 384 wells for reverse transcription.

For the mouse samples E8.0-E9.75, we used the "Tiny Sci" adaptation of the optimized sci-RNA-seq3^17^. Frozen embryos were gently resuspended in 100ul lysis buffer to free the nuclei, then 400ul of DSP-methanol fixative was added. In the same tube, fixed nuclei are rehydrated, washed, then put directly into 8-32 wells for reverse transcription.

After reverse transcription, nuclei were pooled, washed, and redistributed into fresh 96-well plates to attach a second index sequence by ligation. Then the nuclei were pooled again, washed and redistributed into the final plates. There, the nuclei undergo second-strand synthesis, extraction, tagmentation with Tn5 transposase and finally PCR to add the final indexes. The PCR products were pooled, size-selected, and then the library was sequenced on an Illumina NovaSeq. For some experiments, a second NovaSeq run was necessary to capture the extent of the library complexity, so we would add more sequencing reads until the PCR duplication rate met a threshold of 50% or the median UMI count per cell went over 2,500. The validation dataset (Supplementary Fig. 5) generated from 8-21 somites embryos was sequenced on an Illumina NextSeq.

### Processing of sci-RNA-seq3 sequencing reads

Data from each individual sci-RNA-seq3 experiment was processed independently. For each experiment, read alignment and gene count matrix generation was performed using the pipeline that we developed for sci-RNA-seq3^7^ (https://github.com/JunyueC/sci-RNA-seq3_pipeline) with minor modifications: base calls were converted to fastq format using Illumina’s *bcl2fastq*/v2.20 and demultiplexed based on PCR i5 and i7 barcodes using maximum likelihood demultiplexing package *deML*^112^ with default settings. Downstream sequence processing and single cell expression matrix generation were similar to sci-RNA-seq^8^ except that RT index was combined with hairpin adaptor index, and thus the mapped reads were split into constituent cellular indices by demultiplexing reads using both the RT index and ligation index (Levenshtein edit distance (ED) < 2, including insertions and deletions). Briefly, demultiplexed reads were filtered based on RT index and ligation index (ED < 2, including insertions and deletions) and adaptor-clipped using *trim_galore*/v0.6.5 (https://github.com/FelixKrueger/TrimGalore) with default settings. Trimmed reads were mapped to the mouse reference genome (mm10) for mouse embryo nuclei using *STAR*/v2.6.1d^113^ with default settings and gene annotations (GENCODE VM12 for mouse). Uniquely mapping reads were extracted, and duplicates were removed using the unique molecular identifier (UMI) sequence (ED < 2, including insertions and deletions), reverse transcription (RT) index, hairpin ligation adaptor index and read 2 end-coordinate (*i.e.* reads with UMI sequence less than 2 edit distance, RT index, ligation adaptor index and tagmentation site were considered duplicates). Finally, mapped reads were split into constituent cellular indices by further demultiplexing reads using the RT index and ligation hairpin (ED < 2, including insertions and deletions). To generate digital expression matrices, we calculated the number of strand-specific UMIs for each cell mapping to the exonic and intronic regions of each gene with *python*/v2.7.13 *HTseq* package^114^. For multi-mapping reads (*i.e.* those mapping to multiple genes), the read were assigned to the gene for which the distance between the mapped location and the 3’ end of that gene was smallest, except in cases where the read mapped to within 100 bp of the 3’ end of more than one gene, in which case the read was discarded. For most analyses we included both expected-strand intronic and exonic UMIs in per-gene single-cell expression matrices. After the single cell gene count matrix was generated, cells with low quality (UMI < 200 or detected genes < 100 or unmatched_rate (proportion of reads not mapping to any exon or intron) ≥ 0.4) were filtered out. Each cell was assigned to its originating mouse embryo on the basis of the reverse transcription barcode.

### Doublet removal

Here, we performed three steps with the aim of exhaustively detecting and removing potential doublets. Of note, all these analyses were performed separately on data from each experiment.

First, we used Scrublet to detect doublets directly. In this step, we first randomly split the dataset into multiple subsets (six for most of the experiments) in order to reduce the time and memory requirements. We then applied the *scrublet*/v0.1 pipeline^115^ to each subset with parameters (min_count = 3, min_cells = 3, vscore_percentile = 85, n_pc = 30, expected_doublet_rate = 0.06, sim_doublet_ratio = 2, n_neighbors = 30, scaling_method = ’log’) for doublet score calculation. Cells with doublet scores over 0.2 were annotated as detected doublets.

Second, we performed two rounds of clustering and used the doublet annotations to identify subclusters that are enriched in doublets. The clustering was performed based on *Scanpy*/v.1.6.0^18^. Briefly, gene counts mapping to sex chromosomes were removed, and genes with zero counts were filtered out. Each cell was normalized by the total UMI count per cell, and the top 3,000 genes with the highest variance were selected, followed by renormalizing the gene expression matrix. The data was log-transformed after adding a pseudocount, and scaled to unit variance and zero mean. The dimensionality of the data was reduced by PCA (30 components), followed by Louvain clustering with default parameters (resolution = 1). For the Louvain clustering, we first computed a neighborhood graph using a local neighborhood number of 50 using *scanpy.pp.neighbors*. We then clustered the cells into sub-groups using the Louvain algorithm implemented by the *scanpy.tl.louvain* function. For each cell cluster, we applied the same strategies to identify subclusters, except that we set resolution = 3 for Louvain clustering. Subclusters with a detected doublet ratio (by *Scrublet*) over 15% were annotated as doublet-derived subclusters. We then removed cells which are either labeled as doublets by *Scrublet* or that were included in doublet-derived subclusters. Altogether, 2.7% to 16.8% of cells in each experiment were removed by this procedure.

We found that the above *Scrublet* and iterative clustering based approach has difficulty identifying doublets in clusters derived from rare cell types (*e.g.* clusters comprising less than 1% of the total cell population), so we applied a third step to further detect and remove doublets. This step uses a different strategy to cluster and subcluster the data, and then looks for subclusters whose differentially expressed genes differ from those of their associated clusters. This step consists of a series of ten substeps. 1) We reduced each cell’s expression vector to retain only protein-coding genes, lincRNAs, and pseudogenes. 2) Genes expressed in fewer than 10 cells and cells in which fewer than 100 genes were detected were further filtered out. 3) The dimensionality of the data was reduced by PCA (50 components) first on the top 5,000 most highly dispersed genes and then with UMAP (max_components = 2, n_neighbors = 50, min_dist = 0.1, metric = ’cosine’) using *Monocle*/3-alpha^7^. 4) Cell clusters were identified in the UMAP 2D space using the Louvain algorithm implemented in *Monocle*/3 (res = 1e-06). 5) We took the cell clusters identified by *Monocle*/3, downsampled each cluster to 2,500 cells, and computed differentially expressed genes across cell clusters with the *top_markers* function of *Monocle*/3 (reference_cells=1000). 6) We selected a gene set combining the top ten gene markers for each cell cluster (filtering out genes with fraction_expressing < 0.1 and then ordering by pseudo_R2). 7) Cells from each main cell cluster were subjected to dimensionality reduction by PCA (10 components) on the selected set of top cluster-specific gene markers. 8) Each cell cluster was further reduced to 2D using UMAP (max_components = 2, n_neighbors = 50, min_dist = 0.1, metric = ’cosine’). 9) The cells within each cluster were further subclustered using the Louvain algorithm implemented in *Monocle*/3 (res = 1e-04 for most clustering analysis). 10) Subclusters showing low expression of the differentially expressed genes identified in step 5 and enriched expression of specific markers of a different cluster were annotated as doublet-derived subclusters and filtered out. This procedure eliminated 0.5-13.2% of the cells in each experiment.

### Cell clustering and cell-type annotations

For data from individual experiments, after removing the potential doublets detected by the above three steps, we further filtered out the potential low-quality cells by investigating the numbers of UMIs and the proportion of reads mapping to the exonic regions per cell (Supplementary Fig. 3). Then, we merged cells from individual experiments, to generate the final dataset, which included 15 sci-RNA-seq3 experiments with 21 runs of NovaSeq. When we took a first round of cell-embedding, we noticed that one mouse embryo at E14.5 had a grossly reduced proportion of neuronal cells. This particular sample had been divided during pulverization, and we suspect that large portions of the frozen embryo did not make it into the experiment. We removed cells from this E14.5 embryo, and we further filtered out cells from the whole dataset with doublet score (by *Scrublet*) > 0.15 (∼ 0.3% of the whole dataset), as well as cells with either the percentage of reads mapping to ribosomal chromosome (Ribo%) > 5 or the percentage of reads mapping to mitochondrial chromosome (Mito%) > 10 (∼0.1% of the whole dataset). Finally, 11,441,407 cells from 74 embryos were retained, of which the median UMI count per cell is 2,700 and median gene count detected per cell is 1,574. For the final dataset, the number of cells recovered by each embryo and the basic quality information for cells from each sci-RNA-seq3 experiment was summarized in the **Supplementary Table. 1 & 2**. For sex separation and confirmation of embryos with or without sex genotyping, we counted reads mapping to a female-specific non-coding RNA (*Xist*) or chrY genes (except *Erdr1* which is in both chrX and chrY). Embryos were readily separated into females (more reads mapping to *Xist* than chrY genes) and males (more reads mapping to chrY genes than *Xist*).

We then applied *Scanpy*/v.1.6.0^18^ to this final dataset, performing conventional single-cell RNA-seq data processing: 1) retaining protein-coding genes, lincRNA, and pseudogenes for each cell and removing gene counts mapping to sex chromosomes; 2) normalizing the UMI counts by the total count per cell followed by log-transformation; 3) selecting the 2,500 most highly variable genes and scaling the expression of each to zero mean and unit variance; 4) applying PCA and then using the top 30 PCs to calculate a neighborhood graph (n_neighbors = 50), followed by leiden clustering (resolution = 1); 4) performing UMAP visualization in 2D or 3D space (min.dist = 0.1). For cell clustering, we manually adjusted the resolution parameter towards modest overclustering, and then manually merged adjacent clusters if they had a limited number of differentially expressed genes (DEGs) relative to one another or if they both highly expressed the same literature-nominated marker genes. For each of the 26 major cell clusters identified by the global embedding, we further performed a sub-clustering with the similar strategies, except setting n_neighbors = 30 when calculating the neighbor graph and min_dist = 0.3 when performing the UMAP. Subsequently, we annotated individual cell clusters identified by the sub-clustering analysis using at least two literature-nominated marker genes per cell type label (**Supplementary Table 5**).

To be clear, we have hierarchically nominated three levels of cell-type annotations in the manuscript.

a. In the global embedding involving all 11.4 M cells we identified <u>26 major cell clusters</u> (Fig 1**.e-f; Supplementary Table 4**).
b. For individual major cell clusters, we performed sub-clustering, resulting in <u>190 cell types</u> (Supplementary Fig. 4; **Supplementary Table 5**).
c. For a handful of cell types, in specific parts of the manuscript, we performed further sub-clustering, to identify <u>cell subtypes</u>. For example: 1) we re-embedded 745,494 cells from the lateral plate & intermediate mesoderm derivatives, identifying 22 subtypes, most of which are corresponding to different types of mesenchymal cells (Fig. 3f; **Supplementary Table 12**). 2) we re-embedded 296,020 cells (glutamatergic neurons, GABAergic neurons, spinal cord dorsal progenitors, and spinal cord ventral progenitors) from stages <E13, identifying 18 different neuron subtypes (Fig. 5e; **Supplementary Table 12**).

Of note, we processed and analyzed the “birth-series” dataset (*n* = 962,697 nuclei after removing low quality cells and potential doublets cells) and the “early vs. late somites” data (*n* = 104,671 nuclei after removing low quality cells and potential doublets cells) using exactly the same strategy, except without performing sub-clustering on each major cell cluster.

### Quantitatively estimating cell number for individual mouse embryo at any stage during organogenesis

To estimate the cell number of individual embryos, we selected a representative embryo from 12 timepoints at 1 hour increments, from E8.5 to P0 (roughly considered as E19.5). Each embryo was digested with proteinase K overnight, and total genomic DNA was isolated with a Qiagen puregene tissue kit (Qiagen cat. no. 158063). DNA was quantitated and cell number was estimated by taking the total ng recovered, and assuming 2.5 billion base pairs per mouse genome (times two for a diploid cell), 650g per mole of a base pair. Estimating cell number this way does not include any losses due to the DNA preparation, and does not count non-nucleated cells.

Based on the experimentally estimated cell numbers of those 12 embryos, we applied polynomial regression (degree = 3) to fix a curve across embryos between the embryonic day and log2-scaled cell number (adjusted R^2^ > 0.98) (Supplementary Fig. 18a). P0 was treated as E19.5 in the model. Then, the total cell number of a whole mouse embryo at any day between E8.5 and P0 is predicted using the below formula:

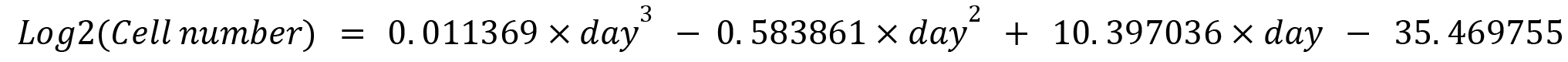

To further estimate the “doubling time” of the total cell number in a whole mouse embryo, at a given timepoint (day), we took the derivatives from the above formula as the log2-scaled proliferation rate *p*(day), and then calculated 24 × 2 / 2^*p*(*day*)^, resulting in an estimate of the number of hours required for the mouse embryo to duplicate its total cell number (Supplementary Fig. 18b).

## Acknowledgments

We thank the members of the Shendure lab for helpful discussions. This work was supported by the Brotman Baty Institute for Precision Medicine, a grant from Paul G. Allen Frontiers Group (Allen Discovery Center for Cell Lineage Tracing to J.S.) and the National Institutes of Health (1UM1HG011586 to W.N.S, J.S and C.M.D..; R01HG010632 to J.S. and C.T.). I.W. and S.A.M were supported by N.I.H. grant UM1OD023222 and the JAX Director’s Innovation Fund. J.S. is an Investigator of the Howard Hughes Medical Institute.

## Data availability

All data used here have been made freely available via http://mouse.gs.washington.edu. The data generated in this study can be downloaded in raw and processed forms from the NCBI Gene Expression Omnibus under accession number GSE186069 and GSE228590. The code used here has been provided from https://github.com/ChengxiangQiu/JAX_code. A website on which the tree of mouse development can be navigated in association with the underlying scRNA-seq data is under development.

## Author Contributions

J.S., M.S., C.Q. designed the research.

I.C.W. collected and staged the mouse embryos, and wrote the corresponding parts of the manuscript.

B.K.M. developed the optimized sci-RNA-seq3 protocol and generated the data (with assistance from R.M.D., T-M.L., E.N., M.T., O.F., D.R.O., A.R.G., S.I.), and wrote the corresponding parts of the manuscript.

C.Q. performed all computational analyses. X.H., S.S., W.S.N., and C.T. assisted with data analysis.

I.C.W., E.N., X.D., C.M.D., N.H., J.C., C.B.M., D.K., A.F.S., M.S., S.A.M., C.T. assisted with results interpretation.

C.Q. and J.S. collaboratively explored and annotated the data, and wrote the manuscript except for sections corresponding to mouse collection/staging and data generation.

J.S. supervised the project.

---

**Supplementary Figure 1.**
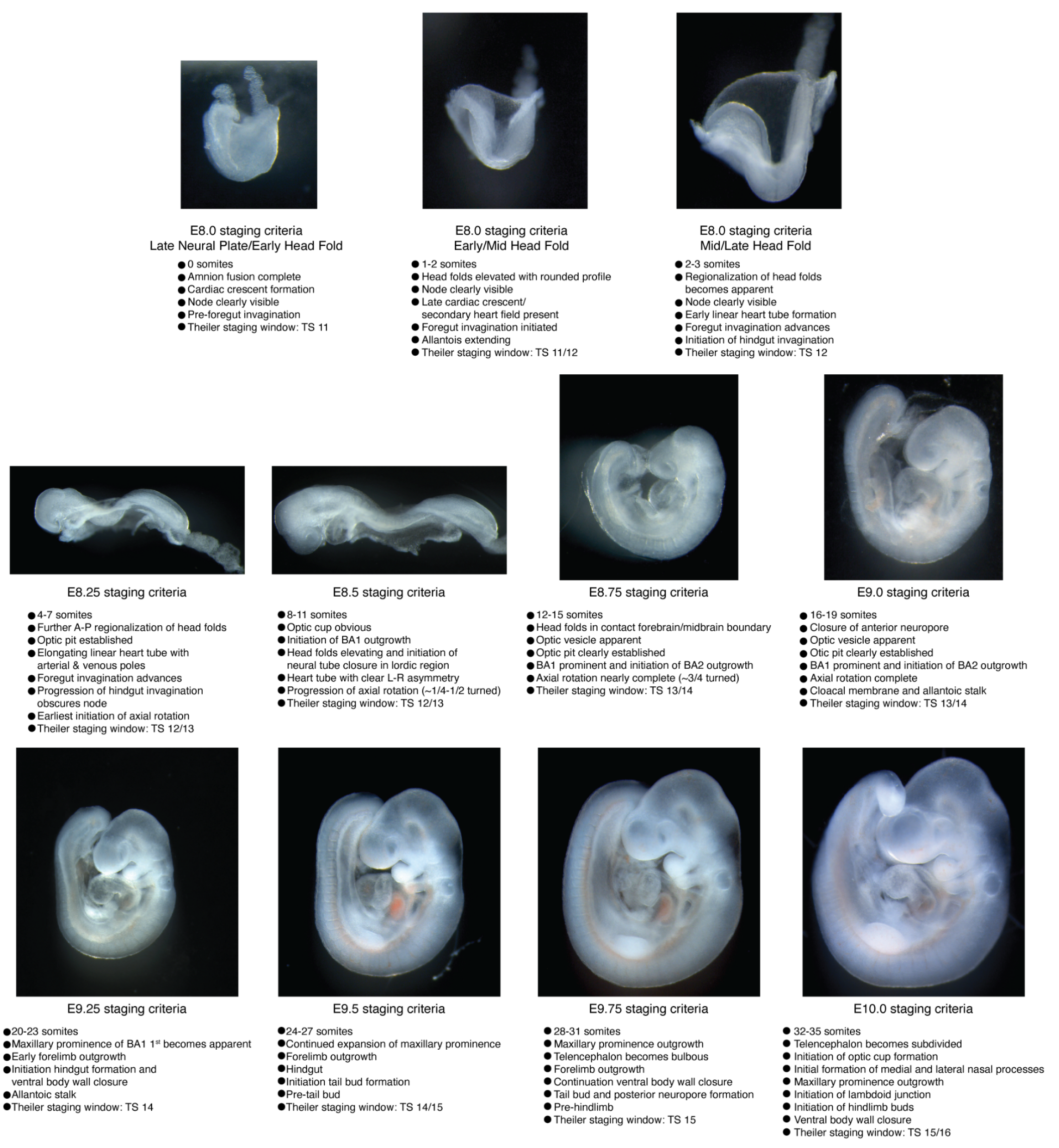
Embryos harvested between E8 and E10 were precisely staged based upon somite counting. Harvested embryos were grouped into bins based on somite counting and further characterized based upon morphological features. Stage-representative images are shown with details of the main staging criteria for each coarse temporal bin listed. The approximately overlapping Theiler Stage (TS) is also noted for reference.

**Supplementary Figure 2.**
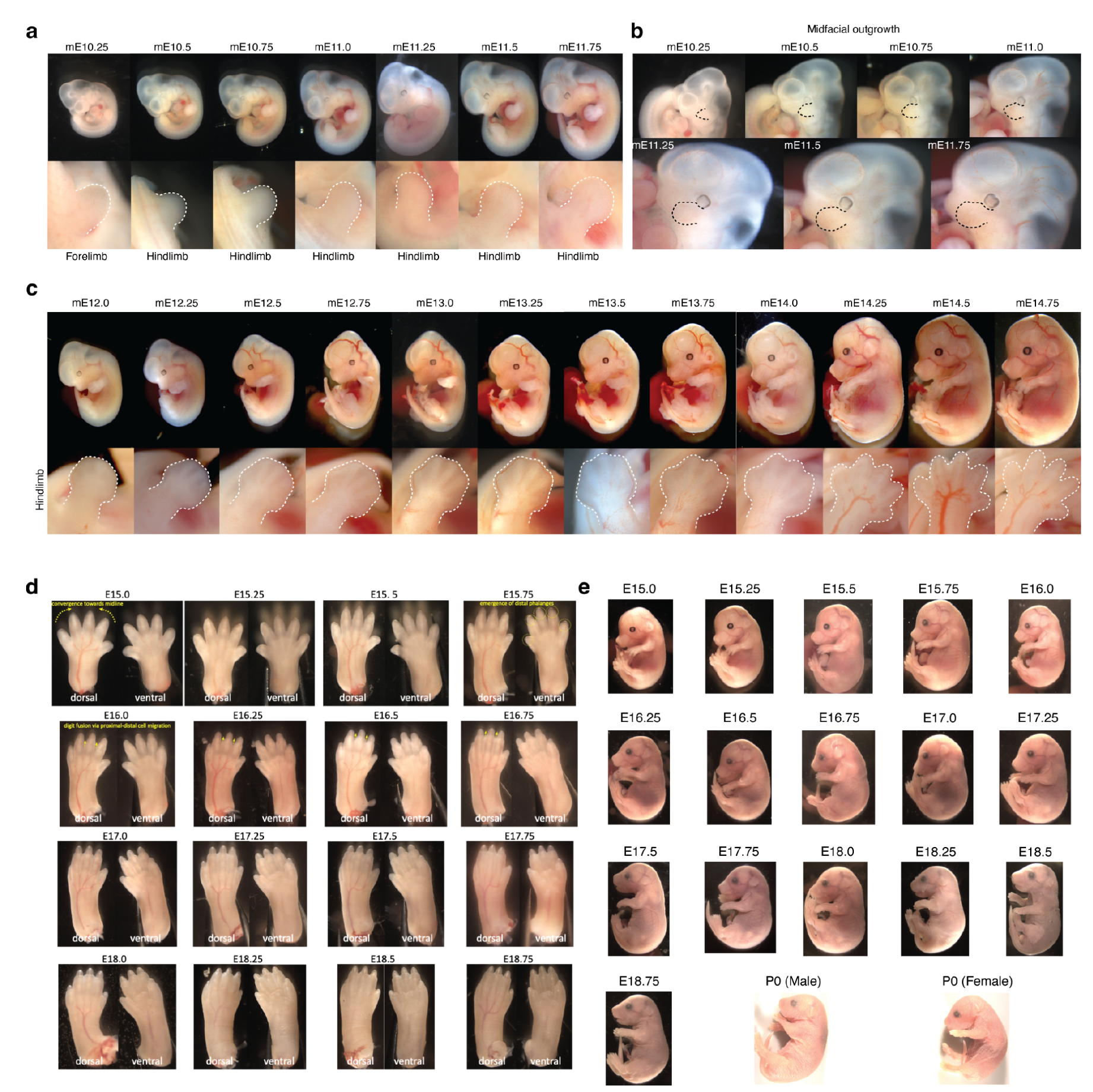
After E10, embryos were precisely staged based on morphological features. This was mainly done using the embryonic mouse ontogenetic staging system (eMOSS), an automated process that leverages limb bud geometry to infer developmental stage^15, 16^. Staging results derived from eMOSS are designated with “mE” for morphometric embryonic day. **a**, For each temporal bin at 6-hr increments from E10.25-E11.75, an image of a stage-representative embryo is shown in the top row. Images of each embryo’s limb bud (white dashed outline) used for staging are shown in the bottom row. **b**, View of the craniofacial region of embryos shown in panel **a** demonstrates that limb bud staging also recreates the ordered ontogenetic progression of craniofacial morphogenesis, including development of the brain, eye, and outgrowth of facial prominences (black dashed line highlights maxillary process). **c**, For each temporal bin at 6-hr increments from E12.0-E14.25, an image of a randomly selected embryo is shown in the top row. The subview of its hindlimb is shown in the bottom row. **d**, eMOSS is able to stage E10.25-E4.75, after which limb morphology becomes too complex. We defined additional dynamics related to digit formation to stage E15.0-E16.75 embryos. However, the remaining timepoints (E17.0-E18.75) were staged based upon gestational age. For each temporal bin at 6-hr increments from E15.0-E18.75, an image of the hindlimbs of a randomly selected embryo is shown. **e**, For each temporal bin at 6-hr increments from E15.0-P0, an image of a stage-representative embryo is shown.

**Supplementary Figure 3.**
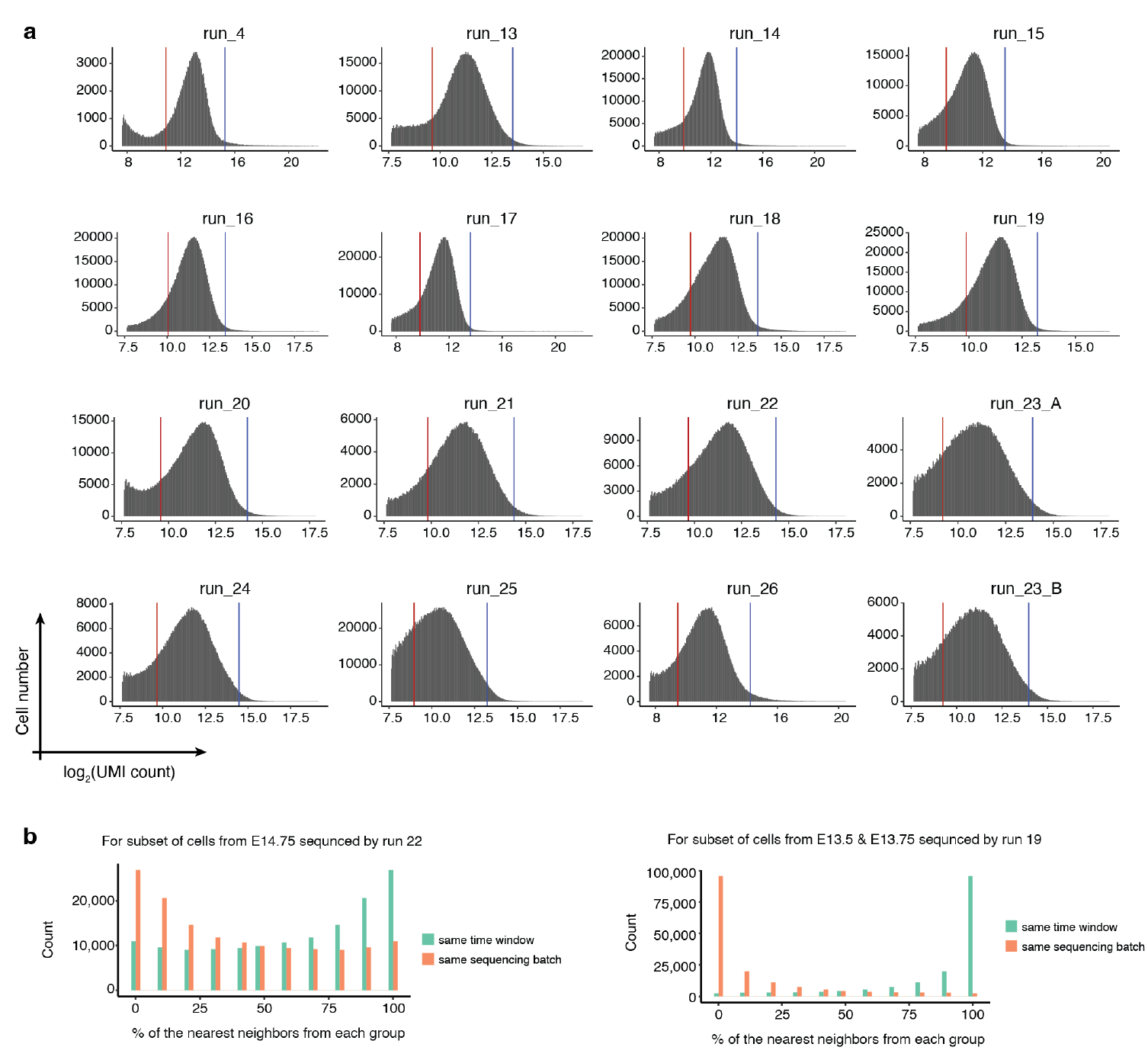
Higher quality sci-RNA-seq3 data as generated by an optimized protocol. **a**, Histograms of log_2_(UMI count) per single nucleus for each of 15 sci-RNA-seq3 experiments. For the 14 newly performed experiments (run_13 to run_26), upper (blue line) and lower (red line) thresholds used for quality filtering correspond to the mean plus 2 standard deviations and mean minus 1 standard deviation of log2-scaled values, respectively, after excluding cells with >85% of reads mapping to exonic regions (except for the lower bound of 500, which was manually assigned for run_25), are shown with vertical lines. The data of run_4, which was reported previously^11^, was subjected to the same thresholds used in the original study, *i.e.* the mean +/- 2 standard deviations of log2-scaled values (blue and red vertical lines, respectively), after excluding cells with >85% of reads mapping to exonic regions. Run_23_A & B were from the same sci-RNA-seq3 experiment, but with nuclei which were sequenced separately. **b**, Although most of the embryos from the same approximate stage (*e.g.* E14.0-E14.75) were included in the same sci-RNA-seq3 experiment (**Supplementary Table 1**), we profiled extra nuclei in some experiments for a handful of timepoints to ensure sufficient coverage. Here we sought to leverage those instances to check for potential batch effects across experiments. For this, on the embedding learned from all of the data, we asked whether these cells’ profiles are more similar to cells from the same experiment or, alternatively, cells from the same time window. Left: for a random subset of cells from E14.75 which were profiled in experiment run_22 (primarily E17.0-E17.75), we performed a *k*-nearest neighbors (*k*NN, *k* = 10) approach in the global 3D UMAP to find the nearest neighboring cells either from the same experiment (red) or the same time window (E14.0-E14.75) but different experiment (blue). The percentages of the nearest neighboring cells from the two groups for individual cells are presented in the histogram. Right: for a random subset of cells from E13.5 & E13.75 which were profiled in experiment run_19 (primarily E10.5-E11), we performed a *k*-nearest neighbors (*k*NN, *k* = 10) approach in the global 3D UMAP to find the nearest neighboring cells either from the same experiment (red) or the same time window (E13.0-E13.75) but a different experiment (blue). The percentages of the nearest neighboring cells from the two groups for individual cells are presented in the histogram. In both examples, we observe that nearest neighbors are overwhelmingly cells from a different experiment (but the same time window), rather than cells from the same experiment (but a different time window).

**Supplementary Figure 4.**
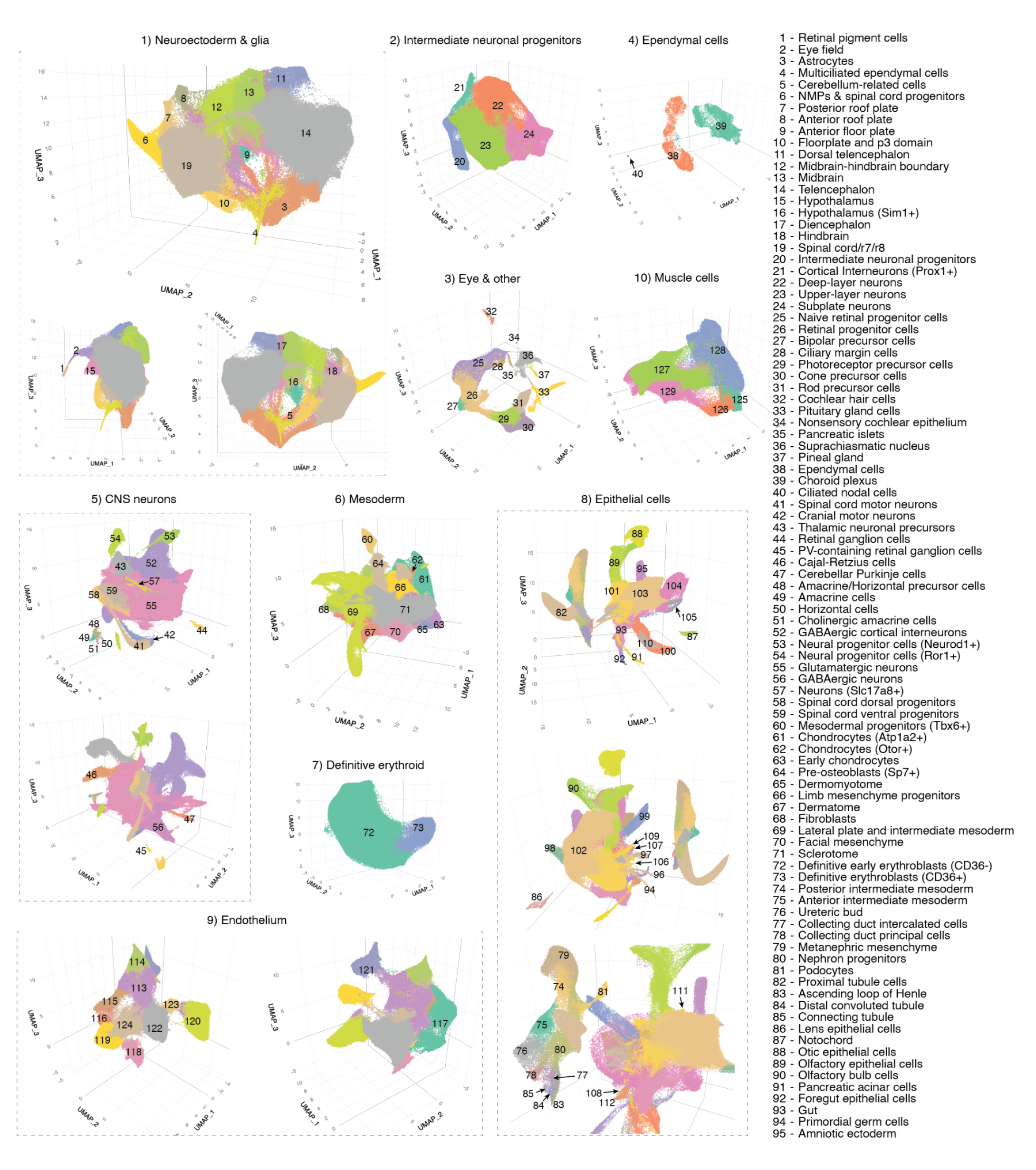
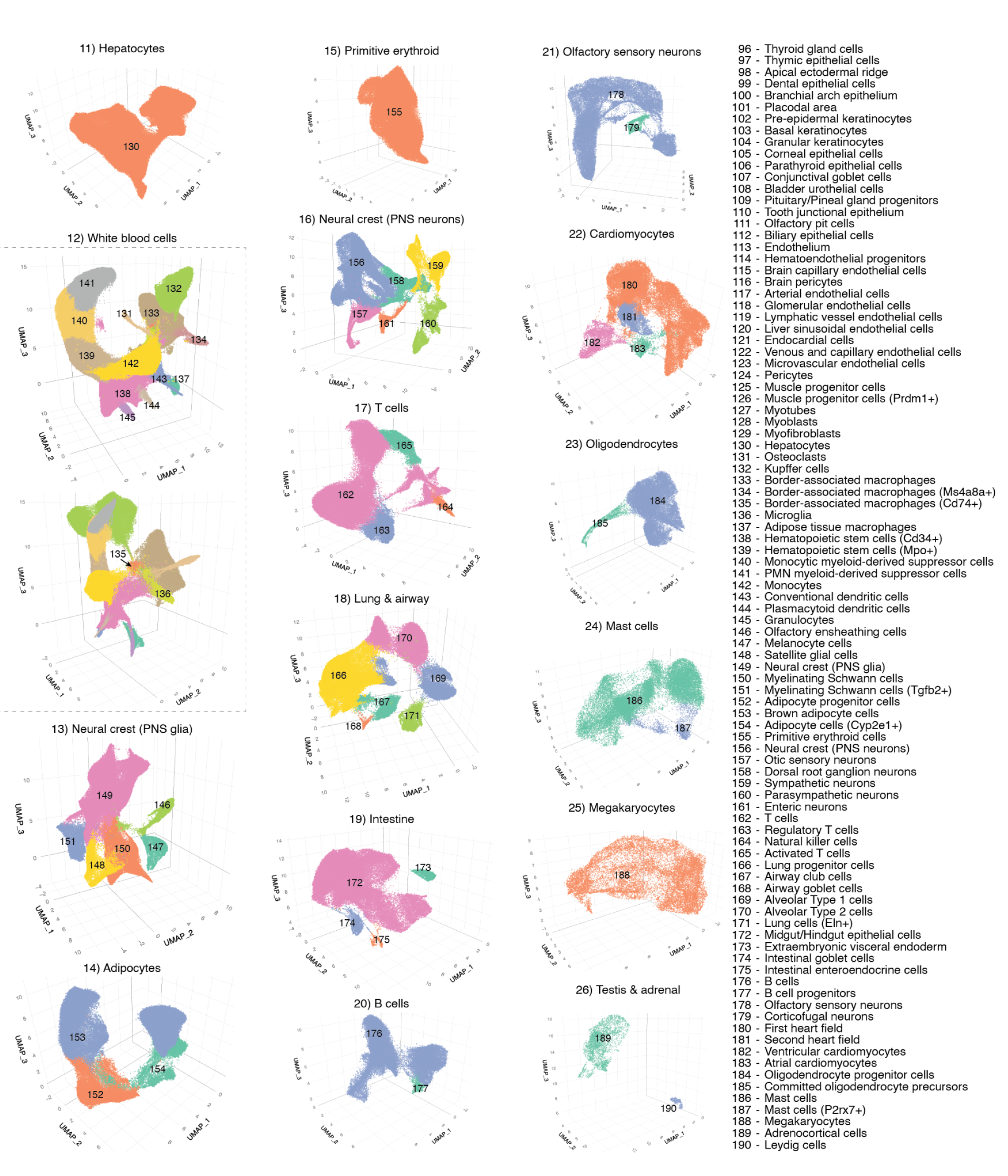
Cell type annotations. For each of the 26 major cell clusters, we performed subclustering and then annotated each of 190 subclusters using at least two literature-nominated marker genes per cell type label (**Supplementary Table 5**).

**Supplementary Figure 5.**
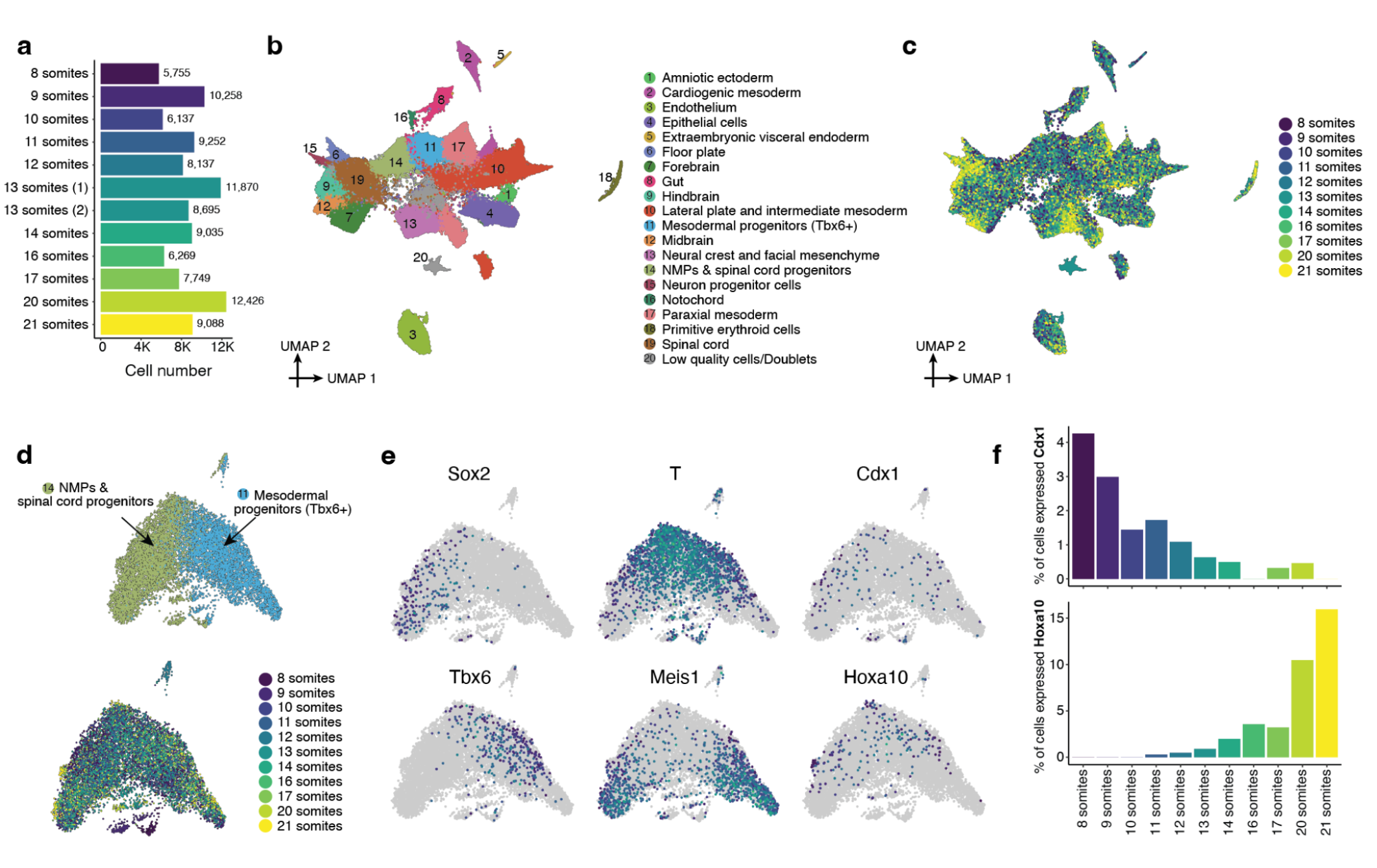
A validation sci-RNA-seq3 dataset of mouse embryos from somites 8 to 21. To validate findings related to differences between embryos staged with early vs. late somite counts, particularly in NMPs, we profiled another 12 precisely staged mouse embryos, ranging from 8 to 21 somites, in an independent sci-RNA-seq3 experiment. The resulting library was sequenced on an Illumina NextSeq 2000, resulting in 104,671 cells in total, with a median UMI count of 513 and a median gene count of 446 per cell. **a**, The number of cells profiled from each embryo. **b**, 2D UMAP visualization of the validation dataset (all cell types). **c**, The same UMAP as in panel b, with cells colored by somite count of the originating embryo. **d**, Re-embedded 2D UMAP of 9,686 cells from NMPs & spinal cord progenitors (cluster 11) and mesodermal progenitors (*Tbx6*+) (cluster 14) in panel b. Cells are colored by either the original annotation (top) or somite count (bottom). **e**, The same UMAP as in panel d, colored by gene expression of marker genes which appear specific to different subpopulations of NMPs: column 1: differences between neuroectodermal (*Sox2*+) vs. mesodermal (*Tbx6*+) fates^35^; column 2: the differentiation of bipotential NMPs (*T*+, *Meis1*-) towards either fate^36, 37^; column 3: earlier (*Cdx1*+) vs. later (*Hoxa10*+) NMPs^24^. **f**, Within the cells shown in panel d, the proportion of cells (y-axis) which express either *Cdx1* (top) or *Hoxa10* (bottom) are plotted as a function of somite count of the originating embryo.

**Supplementary Figure 6.**
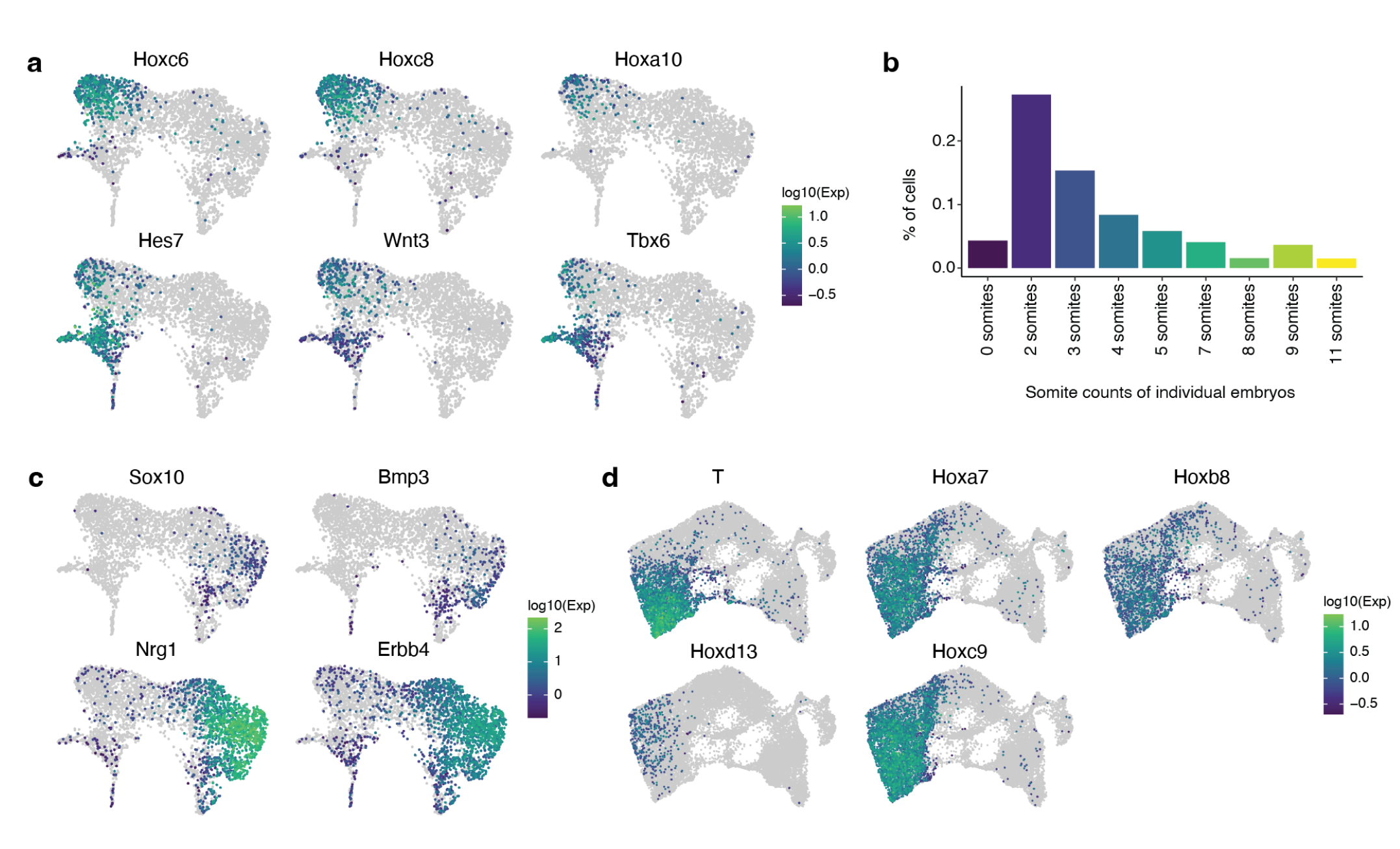
Transcriptional heterogeneity in the posterior embryo during the early somitogenesis. **a**, The same UMAP as in Fig. 2h, colored by gene expression of marker genes which appear specific to the subpopulation of notochord cluster that is *Noto*+, including posterior *Hox* genes (*Hoxc6*, *Hoxc8*, *Hoxa10*), and genes involved in Notch signaling (*Hes7*), Wnt signaling (*Wnt3*) and mesodermal differentiation (*Tbx6*). **b**, Cell proportions falling into the ciliated nodal cell cluster for embryos with different somite counts. **c**, The same UMAP as in Fig. 2h, colored by gene expression of marker genes which appear specific to the subpopulation of the notochord *Noto*- and more strongly *Shh*+, including *Sox10*, *Bmp3*, *Nrg1*, and *Erbb4*. **d**, The same UMAP as in Fig. 2j, colored by gene expression of marker genes which appear specific to the posterior gut endoderm, including *T*, *Hoxa7*, *Hoxb8*, *Hoxd13*, and *Hoxc9*.

**Supplementary Figure 7.**
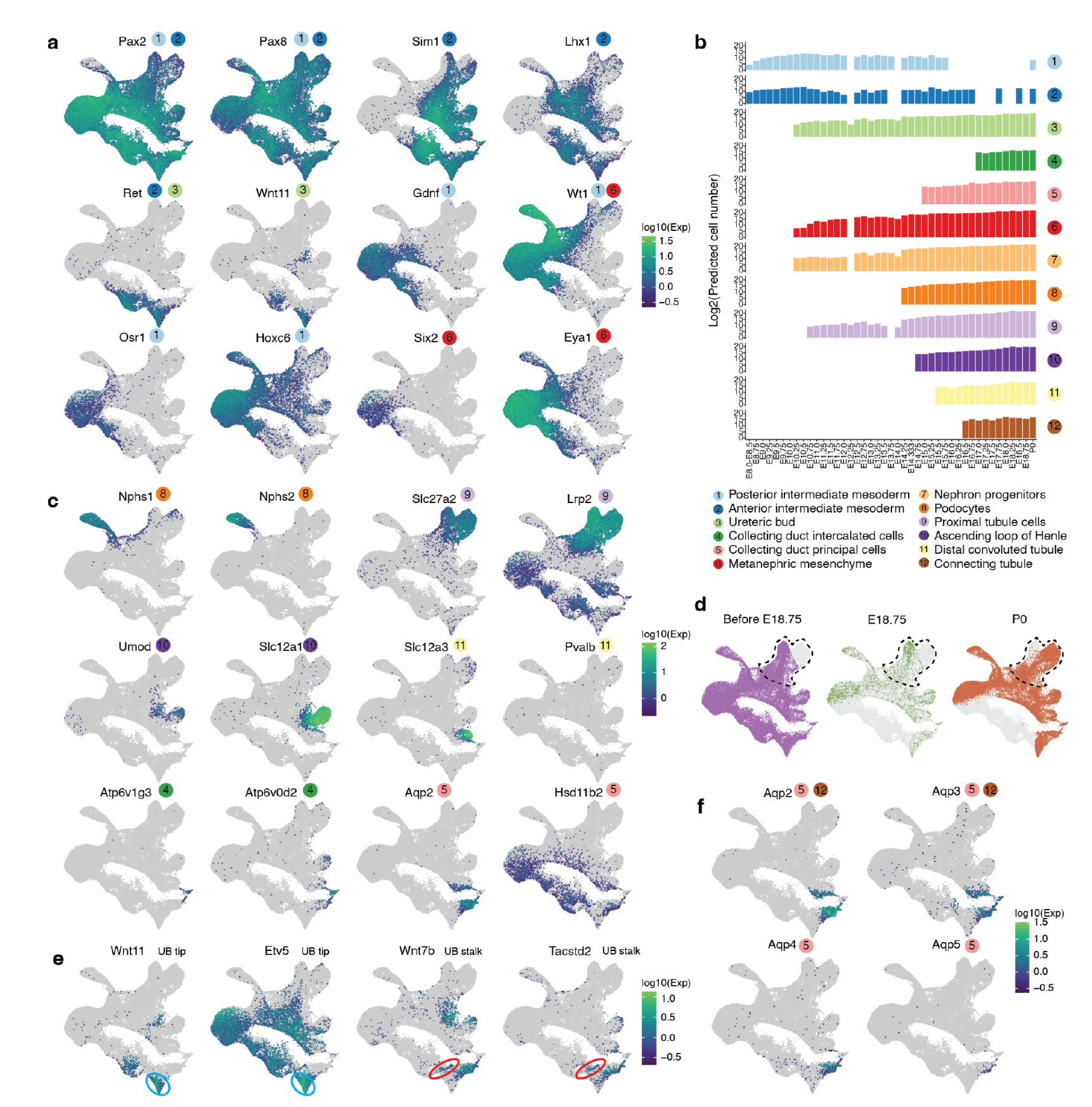
Transcriptional heterogeneity in renal development. **a**, The same UMAP as in Fig. 3a, colored by expression of marker genes which appear specific to anterior intermediate mesoderm (*Pax2*+, *Pax8*+, *Sim1*+, *Lhx1*+, *Ret*+), posterior intermediate mesoderm (*Pax2*+, *Pax8*+, *Gdnf1*+, *Wt1*+, *Osr1*+, *Hoxc6*+), ureteric bud (*Ret*+, *Wnt11*+) or metanephric mesenchyme (*Wt1*+, *Six2*+, *Eya1*+). References for marker genes are provided in **Supplementary Table 5**. **b**, The predicted absolute number (log2 scale) of cells of each renal cell type at each timepoint. The predicted absolute number was calculated by the product of its sampling fraction in the overall embryo and the predicted total number of cells in the whole embryo at the corresponding timepoint (Fig. 1e). For each row, the first timepoint with at least 10 cells assigned that cell type annotation is labeled, and all observations prior to that timepoint are discarded. **c**, The same UMAP as in Fig. 3a, colored by expression of marker genes which appear specific to podocytes (*Nphs1*+, *Nphs2*+), proximal tubule cells (*Slc27a2*+, *Lrp2*+), ascending loop of Henle (*Umod*+, *Slc12a1*+), distal convoluted tubule (*Slc12a3*+, *Pvalb*+), collecting duct intercalated cells (*Atp6v1g3*+, *Atp6v0d2*+) or collecting duct principal cells (*Aqp2*+, *Hsd11b2*+). References for marker genes are provided in **Supplementary Table 5**. **d**, The same UMAP as Fig. 3a is shown three times, with colors highlighting cells from before E18.75 (left), E18.75 (middle), or P0 (right). Dotted cycles highlight cells which appear to correspond to the proximal tubule. **e**, The same UMAP as in Fig. 3a, colored by expression of marker genes which appear specific to the ureteric bud tip (*Wnt11*+, *Ret*+, *Etv5*+) or stalk (*Wnt7b*+, *Tacstd2*+)^54^. Ureteric bud tip and stalk are highlighted by blue and red circles, respectively. **f**, The same UMAP as in Fig. 3a, colored by expression of marker genes which appear specific to connecting tubule cells (*Aqp2*+, *Aqp3*+, *Aqp4*-) or collecting duct cells (*Aqp2*+, *Aqp3*+, *Aqp4*+, *Aqp5*+)^64^.

**Supplementary Figure 8.**
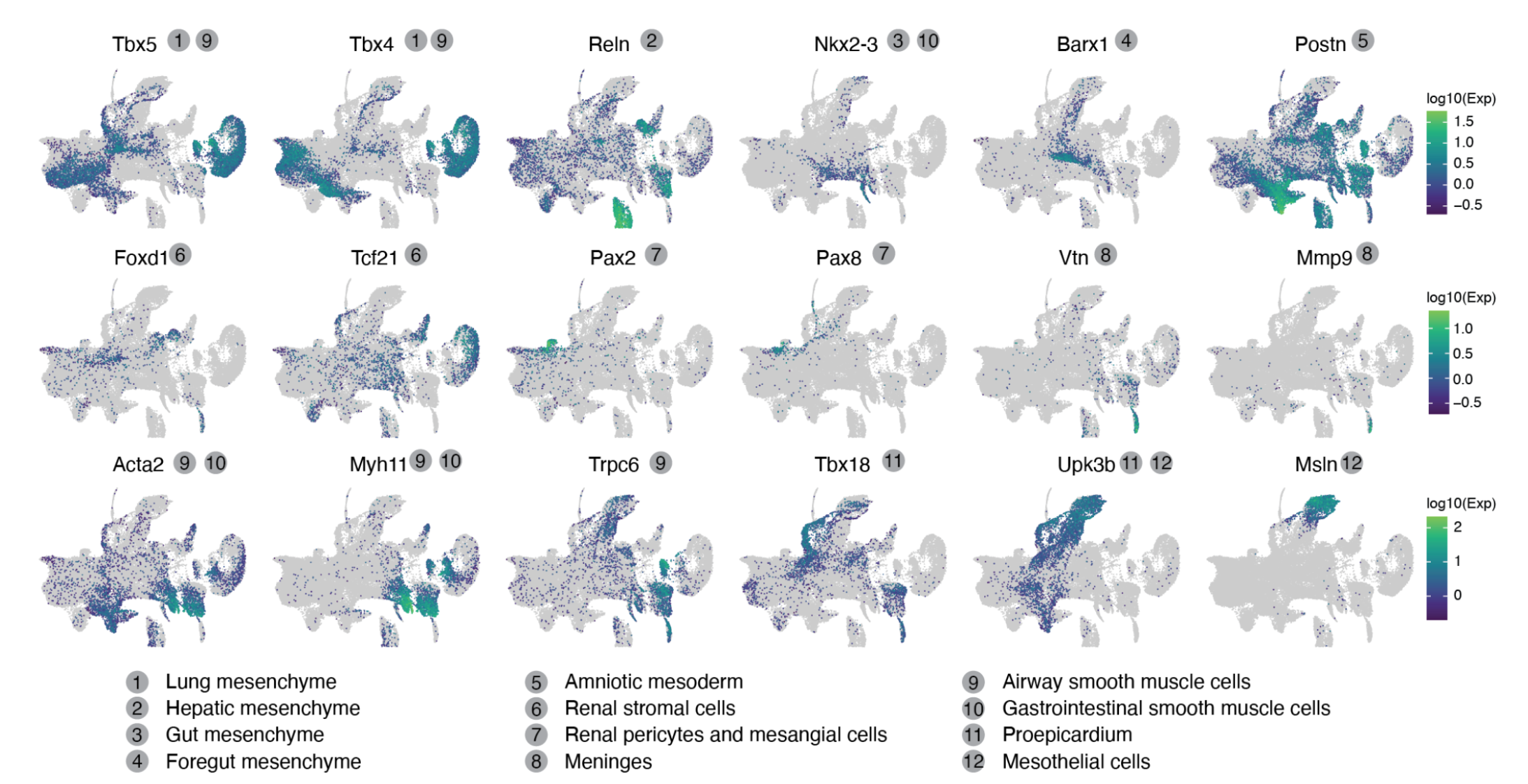
Transcriptional heterogeneity in mesenchyme. The same UMAP as in Fig. 3f, colored by expression of marker genes which appear specific to lung mesenchyme (*Tbx5*+, *Tbx4*+), hepatic mesenchyme (*Reln*+), gut mesenchyme (*Nkx2-3*+), foregut mesenchyme (*Barx1*+), amniotic mesoderm (*Postn*+), renal stromal cells (*Foxd1*+, *Tcf21*+), renal pericytes and mesangial cells (*Pax2*+, *Pax8*+), meninges (*Vtn*+), airway smooth muscle cells (*Trpc6*+, *Tbx5*+), gastrointestinal smooth muscle cells (*Nkx2-3*+), proepicardium or mesothelium (*Msln*+). References for marker genes are provided in **Supplementary Table 12**.

**Supplementary Figure 9.**
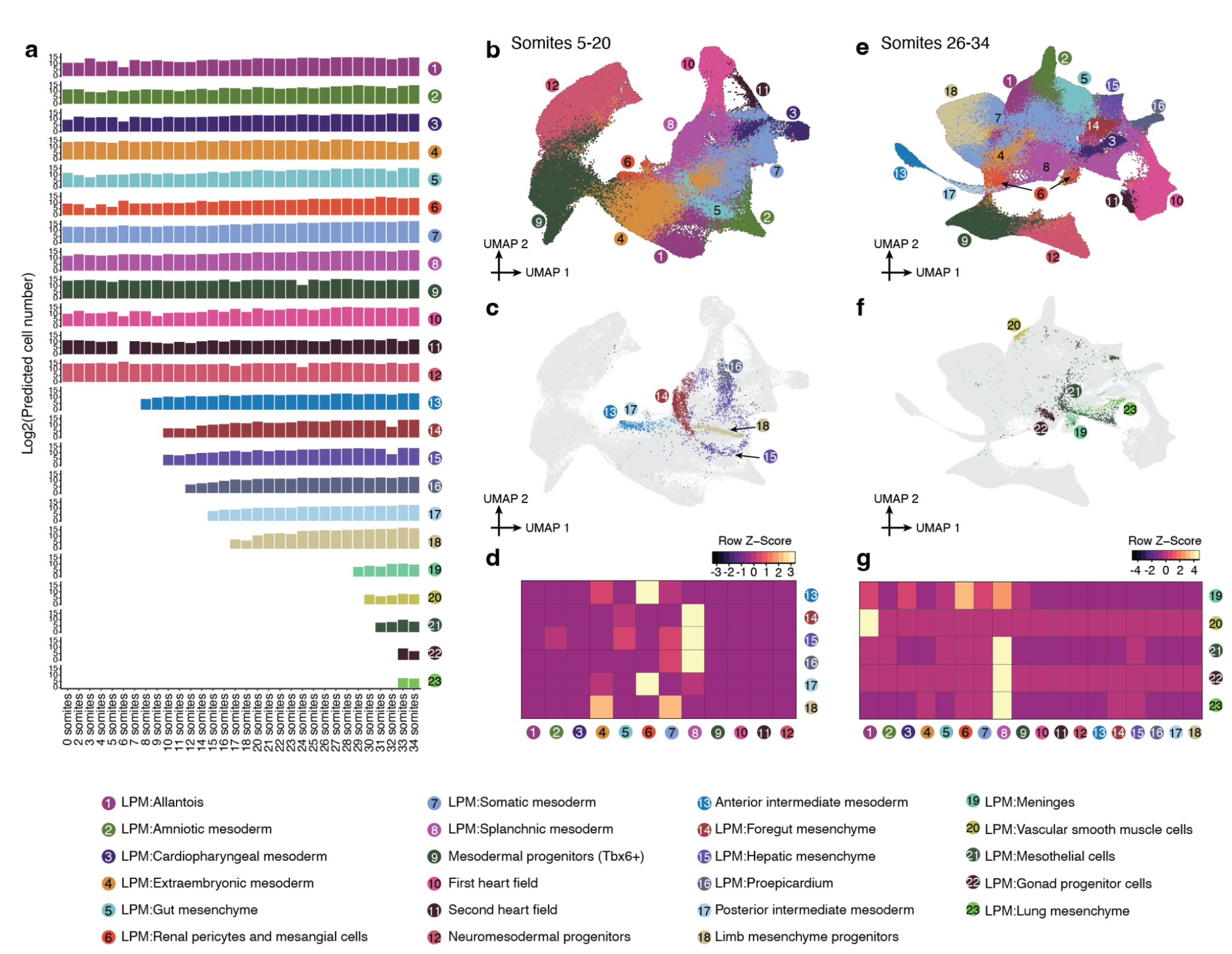
The emergence of mesenchymal subtypes from the patterned mesoderm. **a**, The predicted absolute number (log2 scale) of cells of each mesoderm cell type at each somite count. The predicted absolute number was calculated by the product of its sampling fraction in the overall embryo and the predicted total number of cells in the whole embryo at the corresponding timepoint. Because cell numbers were only predicted for the broader bins (Fig. 1e), rather than individual somite counts, these were used for roughly corresponding sets (0-12 somite stage: E8.5; 14-15 somite stage: E8.75; 16-18 somite stage: E9.0; 20-23 somite stage: E9.25; 24-26 somite stage: E9.5; 27-31 somite stage: E9.75; 32-34 somite stage: E10.0). For each row, the first somite count with at least 10 cells assigned that cell type annotation is labeled, and all observations prior to that somite count are discarded. **b**, Re-embedded 2D UMAP of 110,753 cells from the selected cell types of mesoderm (clusters 1-12 as listed in panel a) from 5-20 somite stage embryos. **c**, The same UMAP as in panel b, but with inferred progenitor cells colored by derivative cell type with the highest mutual nearest neighbors (MNN) pairing score. **d**, Normalized MNN pairing score between mesodermal territories (column) and their inferred derivative cell types (row) from 5-20 somite stage embryos. The selected cell populations are first embedded into 30 dimensional PCA space, and then for individual derivative cell types, MNN pairs (k = 10 used for k-NN) between their earliest 500 cells (in absolute time) and cells from mesodermal territories are identified. **e**, Re-embedded 2D UMAP of 275,000 cells from the selected cell types of mesoderm (clusters 1-12 as listed in panel a) from 26-34 somite stage embryos. **f**, The same UMAP as in panel e, but with inferred progenitor cells colored by derivative cell type with the highest MNN pairing score. **g**, Normalized MNN pairing score between mesodermal territories (column) and their inferred derivative cell types (row) from 26-34 somite stage embryos. The selected cell populations are first embedded into 30 dimensional PCA space, and then for individual derivative cell types, MNN pairs (k = 10 used for k-NN) between their earliest 500 cells (in absolute time) and cells from mesodermal territories are identified.

**Supplementary Figure 10.**
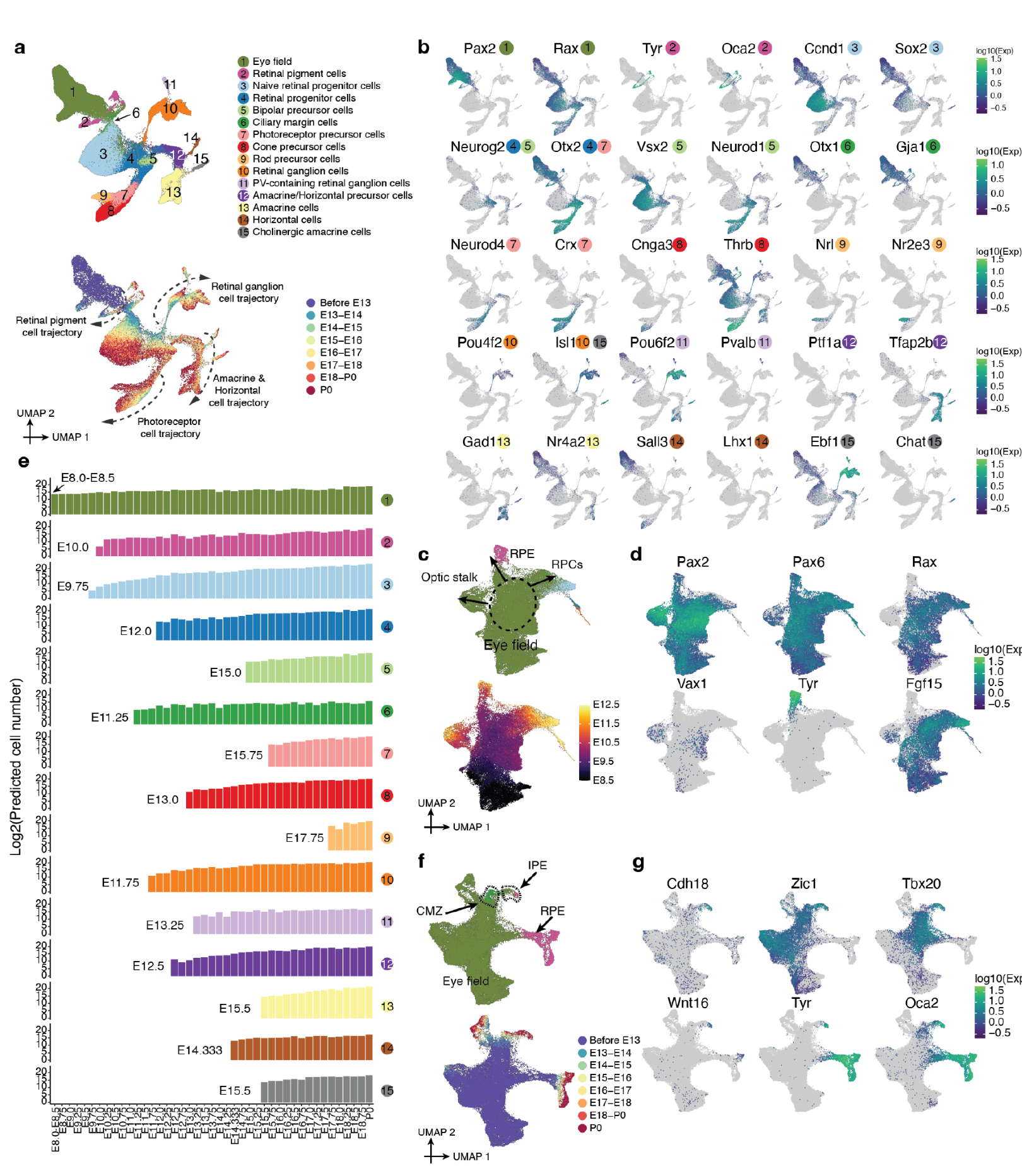
The timing and trajectories of retinal development. **a**, Re-embedded 2D UMAP of 160,834 cells corresponding to the retinal development from E8 to P0. Cells are colored by either their initial annotations (top) or timepoint (bottom, after downsampling to a uniform number of cells per time window). Arrows highlight four of the main trajectories observed. Same as Fig. 4a except 2D instead of 3D projection. **b**, The same UMAP as in panel a, colored by gene expression of marker genes for each annotated retinal cell type. References for marker genes are provided in **Supplementary Table 5**. **c**, Re-embedded 2D UMAP of the subset of cells in panel a from stages &lt;= E12.5. Cells are colored by either their initial annotations (top) or timepoint (bottom). **d**, The same UMAP as in panel c, colored by gene expression of markers of retinal progenitor cells RPCs (*Pax2*+, *Pax6*+, *Rax*+, *Fgf15*+)^69^, RPE (*Tyr*+)^70^, and the optic stalk (*Pax2*+, *Vax1*+, *Rax*-)^71^. **e**, The predicted absolute number (log2 scale) of cells of each retinal cell type at each timepoint. The predicted absolute number was calculated by the product of its sampling fraction in the overall embryo and the predicted total number of cells in the whole embryo at the corresponding timepoint (Fig. 1e). For each row, the first timepoint with at least 10 cells assigned that cell type annotation is labeled, and all observations prior to that timepoint are discarded. **f**, Re-embedded 2D UMAP of a subset of cells in panel a corresponding to eye field, RPE and CMZ. Cells are colored by either their initial annotations (top) or timepoint (bottom). **g**, The same UMAP as in panel f, colored by gene expression of marker genes for IPE^72^ or pigment epithelium more generally (*Tyr* & *Oca2*). RPE: retinal pigment epithelium. CMZ: ciliary marginal zone. RPCs: retinal progenitor cells. IPE: iris pigment epithelium.

**Supplementary Figure 11.**
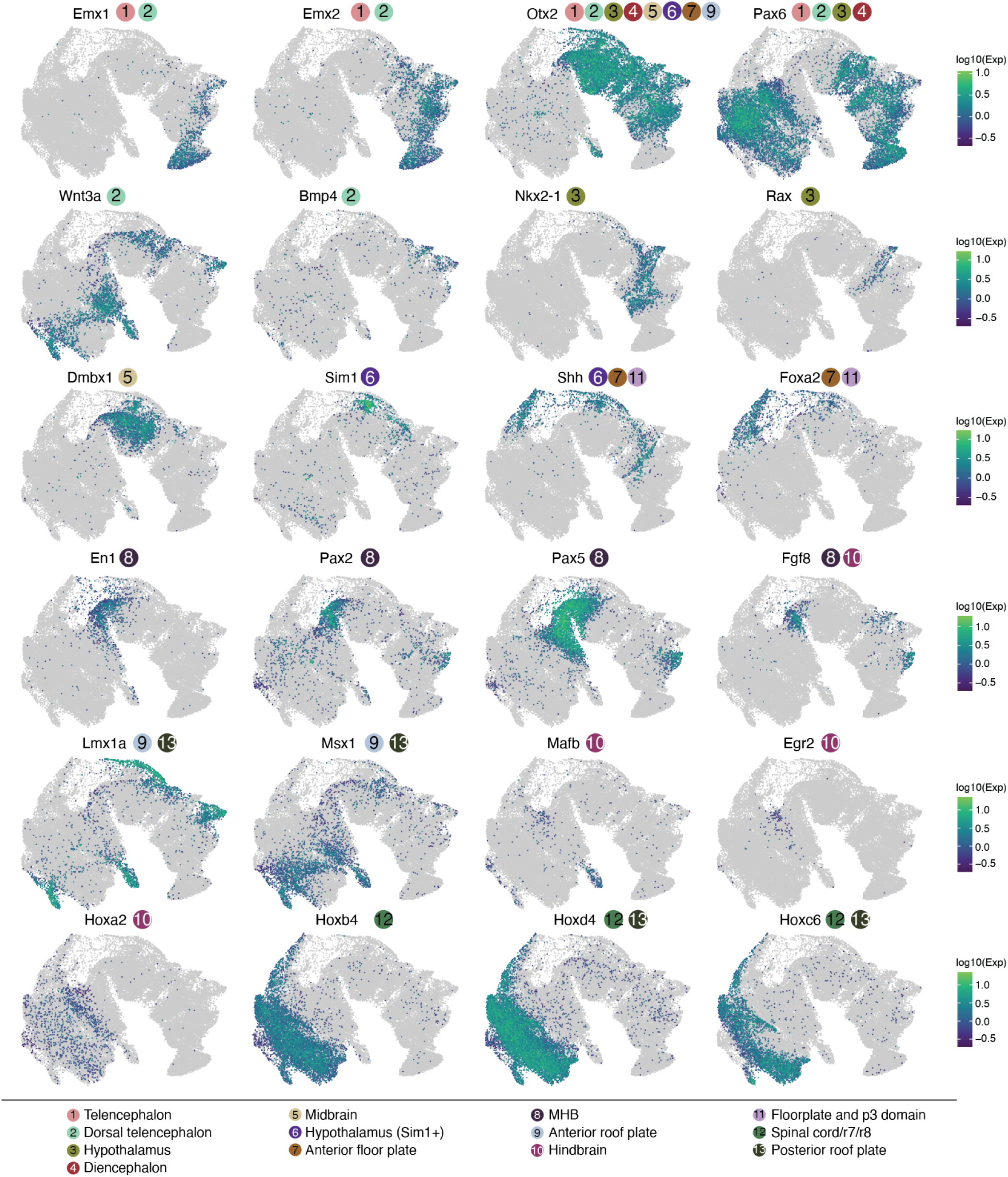
Marker gene expression for different neuroectodermal territories. The same UMAP as in Fig. 5a, colored by gene expression of marker genes for each neuroectodermal territory. References for marker genes are provided in **Supplementary Table 5**.

**Supplementary Figure 12.**
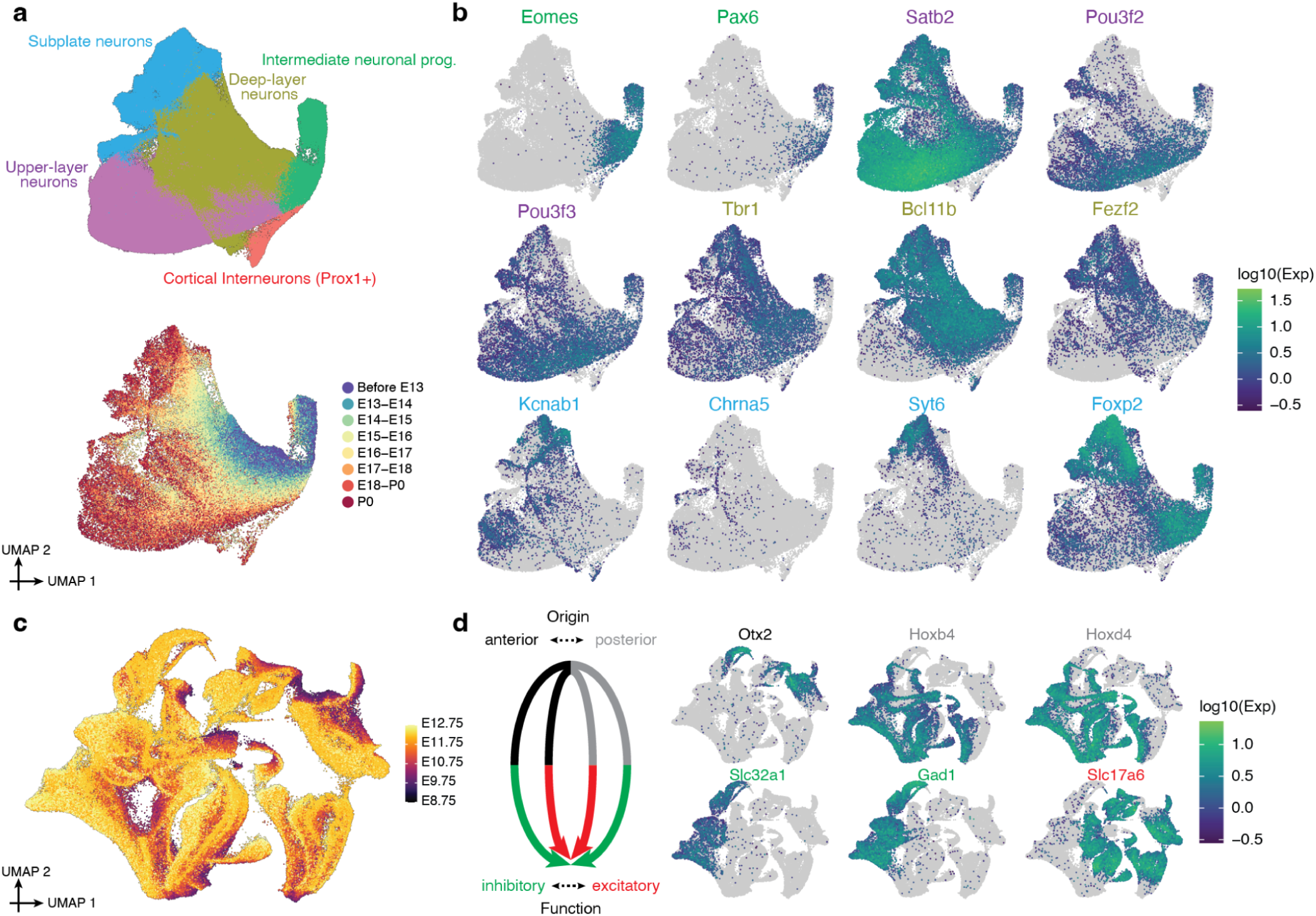
Subtypes of intermediate neuronal progenitors, glutamatergic & GABAergic neurons. **a**, Re-embedded 2D UMAP of 628,251 cells within the intermediate neuronal progenitors major cell cluster, colored by either cell type (top) or developmental stage (bottom, after downsampling to a uniform number of cells per time window). **b**, The same UMAP as in panel a, colored by gene expression of marker genes which appear specific to intermediate neuronal progenitors (*Eomes*+, *Pax6*+), upper-layer neurons (*Satb2*+, *Pou3f2*+, *Pou3f3*+), deep-layer neurons (*Tbr1*+, *Bcl11b*+, *Fezf2*+), or subplate neurons (*Kcnab1*+, *Chrna5*+, *Syt6*+, *Foxp2*+). References for marker genes are provided in **Supplementary Table 5**. **c**, The same UMAP as in Fig. 5e, with cells colored by timepoints. **d**, Left: Neuronal subtypes shown in Fig. 5e originate from anterior vs. posterior of neuroectoderm, and then subsequently display inhibitory vs. excitatory functions after differentiation. Right: The same UMAP as in Fig. 5e, colored by gene expression of marker genes which appear specific to anterior (*Otx2*+)^79^ vs. posterior (*Hoxb4*+, *Hoxd4*+)^80^ origins, or inhibitory (*Slc32a1*+, *Gad1+*)^81^ vs. excitatory (*Slc17a6*+)^82^ functions.

**Supplementary Figure 13.**
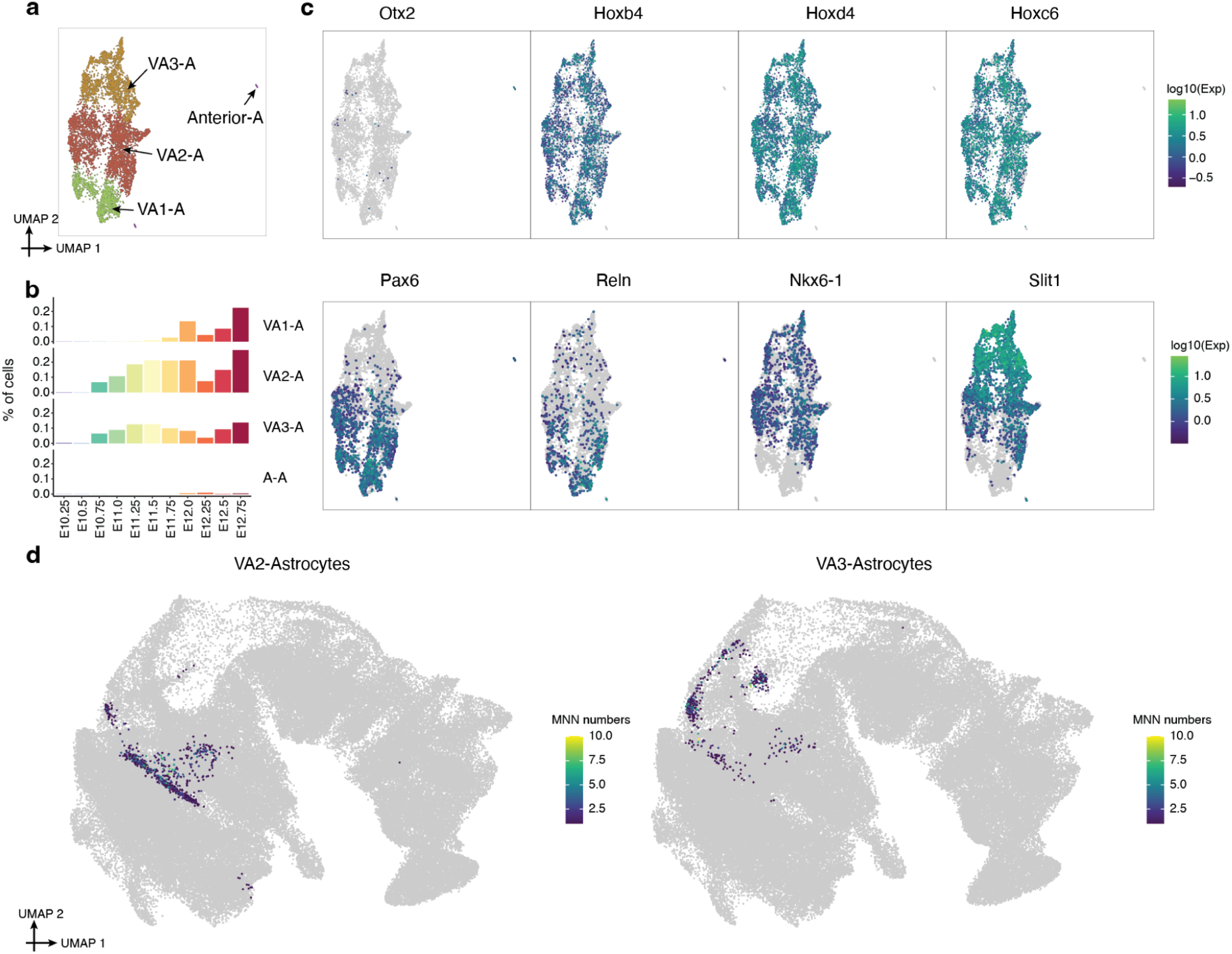
Subtypes of early astrocytes and their inferred progenitors. **a**, Re-embedded 2D UMAP of 5,928 cells within the astrocytes from stages <E13. **b**, Composition of embryos from each 6-hr bin by different subpopulations of astrocytes. **c**, The same UMAP as in panel a, colored by gene expression of marker genes which appear specific to anterior (*Otx2*+) or posterior *(Hoxb4*+, *Hoxd4*, *Hoxc6*+) astrocytes, VA1-astrocytes (*Pax6*+, *Reln*+), VA2-astrocytes (*Pax6*+, *Reln*+, *Nkx6-1*+, *Slit1*+), and VA3-astrocytes (*Nkx6-1*+, *Slit1*+)^76^. **d**, The same UMAP of the patterned neuroectoderm as in Fig. 5a, with inferred progenitor cells of astrocytes colored by the frequency that has been identified as a MNN with either VA2-astrocytes (left) or VA3-astrocytes (right).

**Supplementary Figure 14.**
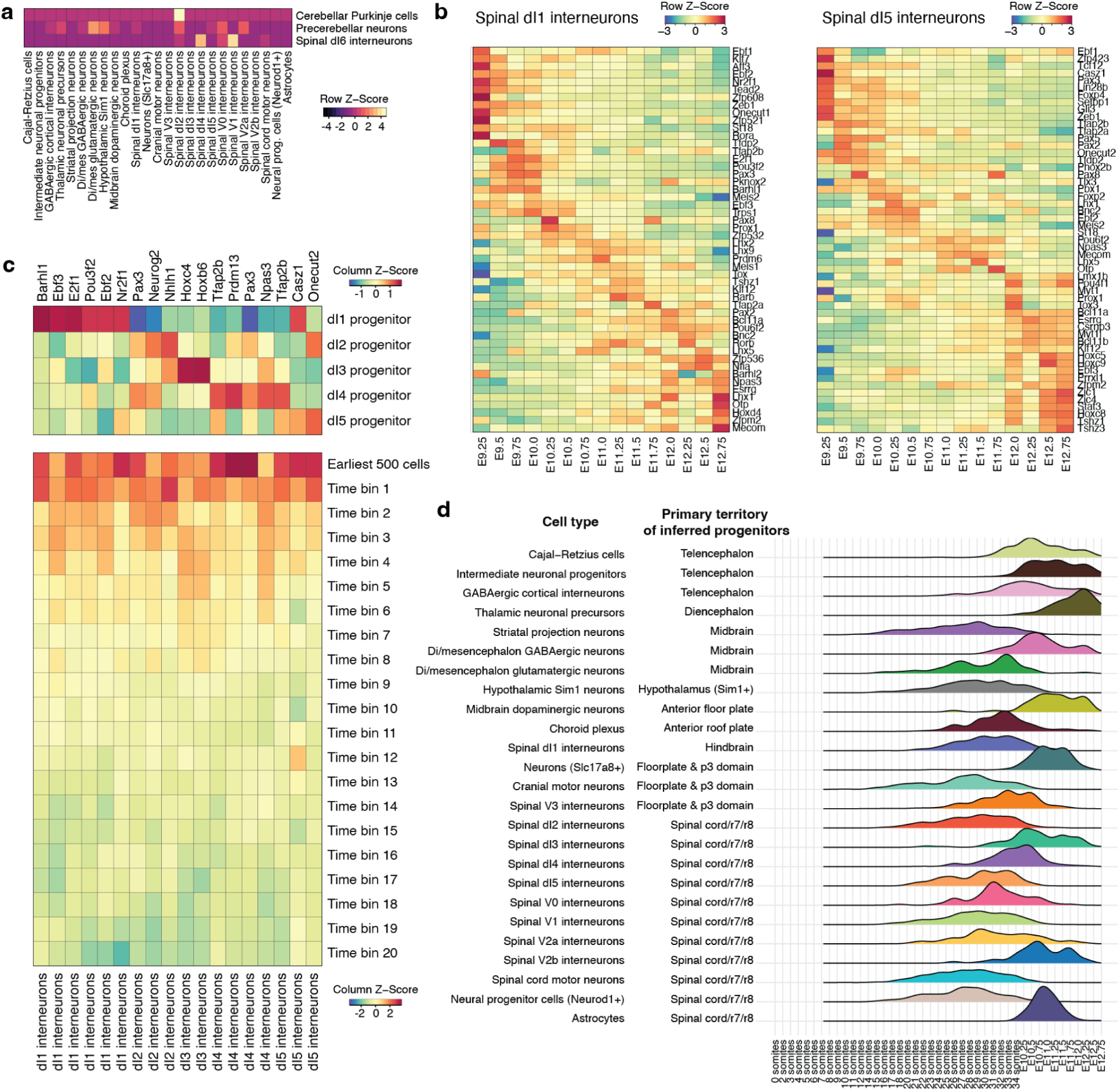
The timing of neuronal subtype differentiation from the patterned neuroectoderm. **a**, For those three cell types (cerebellar Purkinje cells, precerebellar neurons, spinal dI6 interneurons) which were excluded in Fig. 5d-e due to having fewer than 50 MNN pairs, we performed a recursive mapping to identify whether they might share progenitors with another derived cell type, essentially repeating the analysis but attempting to map the earliest cells of these cell types to other derivative cell types rather than the patterned neuroectoderm. The heatmap shows the number of MNN pairs between pairwise cell types. In brief, this analysis suggests that spinal dl2 interneurons and cerebellar astrocytes share progenitors, while the progenitors of the other two re-analyzed cell types remain ambiguous. **b**, Gene expression across timepoints, for the specific TF markers of spinal dI1 (left) or spinal dI5 (right) interneurons. **c**, Top: gene expression for 18 selected TFs, across progenitor cells of dI1-5 from the neuroectodermal territories. Bottom, gene expression for 18 selected TFs across 21 time bins for dl1-5 spinal interneurons in which the TF has been nominated as marker TF. For individual spinal interneurons (each column), the first time bin involves the earliest 500 cells, then the left cells break into 20 bins ordered by their timepoints and with the same number of cells in each bin. Only cells from stages &lt;E13 are included. **d**, For each neuronal subtype in Fig. 5i-j, we selected the annotation in the patterned neuroectoderm to which the most inferred progenitors had been assigned, and plotted the distribution of timepoints for that subset of inferred progenitors.

**Supplementary Figure 15.**
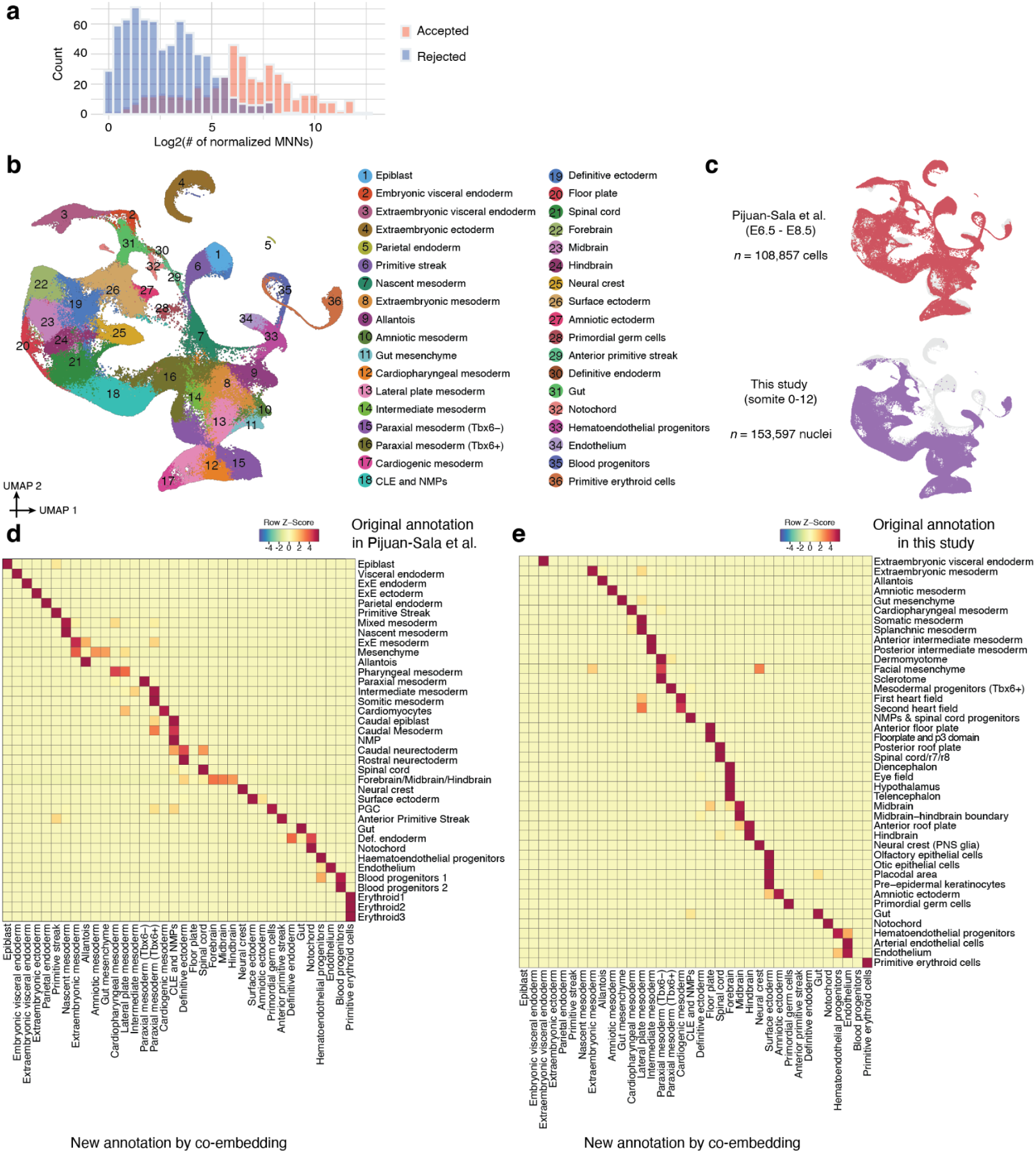
Integration of scRNA-seq profiles from gastrulation and early somitogenesis to identify equivalent cell type nodes across datasets generated by distinct technologies. **a**, 1,155 edges with the number of normalized MNNs > 1 were manually reviewed for biological plausibility. Histogram of edges that were accepted or rejected as a function of normalized MNN score. **b**, 2D UMAP visualization of co-embedded cells, derived both from a gastrulation dataset based on cells from E6.5 to E8.5 generated on the 10x Genomics platform^6^ (*n* = 108,857 cells) and the earliest ∼1% of this dataset (0-12 somite stage embryos) generated by sci-RNA-seq3 (*n* = 153,597 nuclei), after batch correction^51^. This is essentially an updated version of an analysis that we have done previously^11^. We performed clustering and cell type annotation on the integrated co-embedding, as shown. **c**, The same UMAP as in panel b is shown twice, with colors highlighting cells/nuclei from Pijuan-Sala’s dataset^6^ (top) or early somitogenesis^11^ (bottom). **d**, For cells from the original Pijuan-Sala’s dataset^6^, we quantify and display the overlap between the original annotations and the new annotations shown in panel b. For each row, the proportions of cells that are distributed across each column are transformed to z-score. **e**, For nuclei from the early somitogenesis embryos^11^, we quantify and display the overlap between the original annotations and the new annotations shown in panel b. These mappings were the basis for dataset equivalence edges between the “gastrulation” and 12 “organogenesis & fetal development” subsystems. For each row, the proportions of cells that are distributed across each column are transformed to z-score. CLE: Caudal lateral epiblast. NMPs: Neuromesodermal progenitors.

**Supplementary Figure 16.**
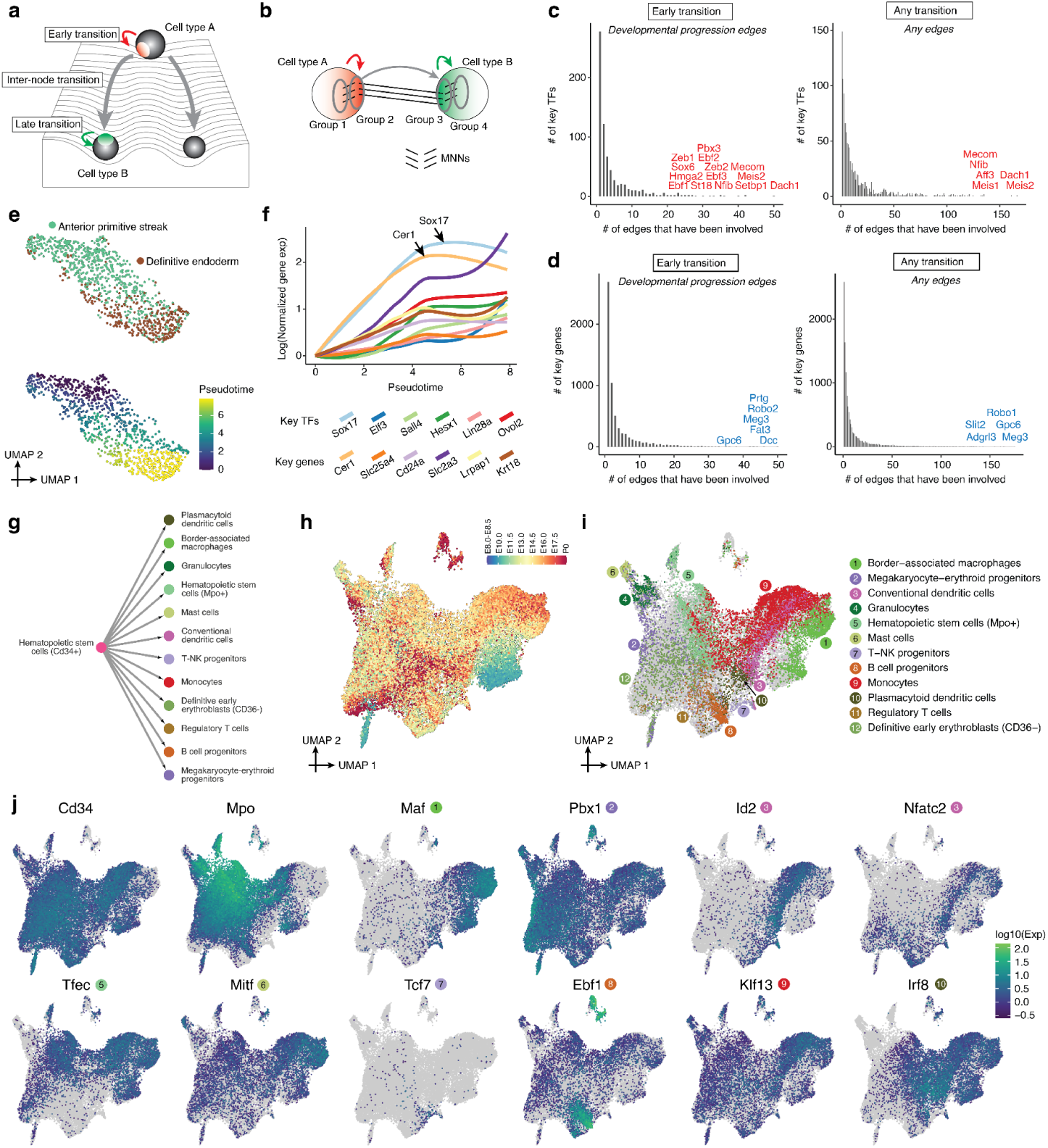
Systematic nomination of TFs and other genes for cell type specification. **a**, A Waddington landscape cartoon illustrating how a cell type transition might be broken into three phases. **b**, Given a directional edge between two nodes, A→B, we identified the subset of cells within each node that were either MNNs of the other cell type (inter-node; groups 2 & 3) or MNNs of those cells (intra-node; groups 1 & 4). If A→B, this effectively models the transition as group 1→2→3→4. **c**, Histograms of the number of edges in which TFs are differentially expressed. The left histogram counts only genes when they are differentially expressed across the early phase of an developmental progression edge, while the right histogram counts genes when they are differentially expressed in any phase of all edges. **d**, Same as panel c, but for all genes rather than only TFs. **e**, Re-embedded 2D UMAP of 988 cells participating in groups 1-4 of the transition from anterior primitive streak → definitive endoderm. Cells are colored by either cell type annotations (top) or estimated pseudotime (bottom) using *Monocle*3^7^. **f**, For cells in panel e, normalized gene expression of selected genes are plotted as a function of estimated pseudotime. Gene expression values were calculated from original UMI counts normalized to total UMIs per cell, followed by natural-log transformation. The line of gene expression was plotted by the *geom_smooth* function in ggplot2. We manually added an offset based on their expression at pseudotime = 0 to the y-axis for individual genes. **g**, A sub-graph of Fig. 6g, including hematopoietic stem cells (Cd34+) and 12 cell type nodes which appear derived from it. **h**, Re-embedded 2D UMAP of 37,750 cells from hematopoietic stem cells (Cd34+), colored by developmental stage (after downsampling to a uniform number of cells per stage). **i**, The same UMAP as in panel h, but with inferred progenitor cells (the cells participating in the MNNs that support the edges) colored by derivative cell type with the most frequent MNN pairs. **j**, The same UMAP as in panel h, colored by gene expression of selected top key TFs which were upregulated during the “early transition” for each derivative.

**Supplementary Figure 17.**
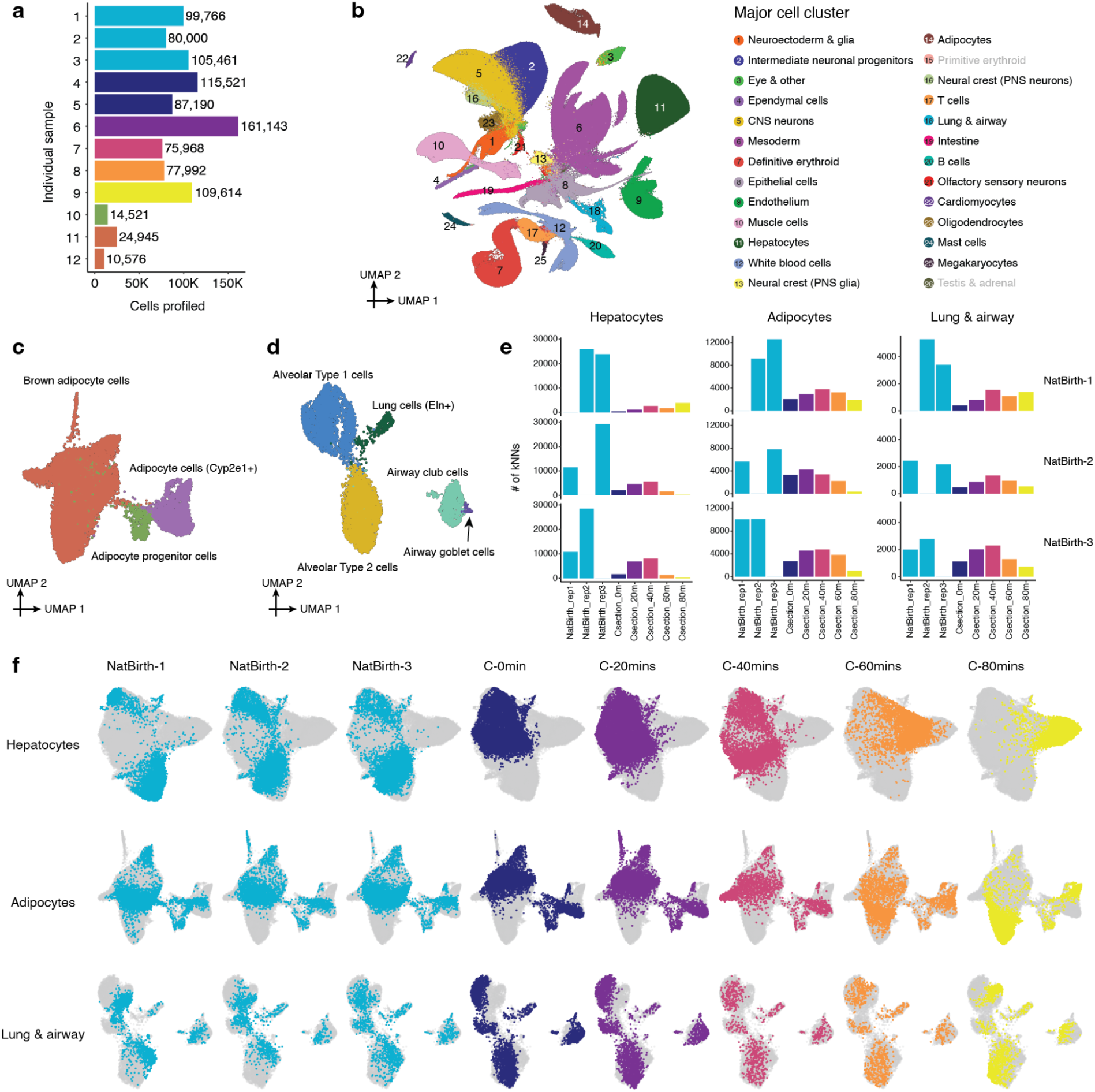
Rapid shifts in transcriptional state occur in a restricted subset of cell types upon birth, and differ between vaginally and C-section delivered pups. **a,** The number of nuclei profiled for each animal shown in Fig. 7c. A small number of nuclei from additional fetal samples from the original set of experiments were also profiled for quality control (samples 10-12). **b**, 2D UMAP visualization of the birth-series dataset (*n* = 962,697 cells). Colors correspond to 26 major cell cluster annotations (Fig. 1e). Two major cell clusters (the primitive erythroid and testis & adrenal major cell clusters) shown in the original dataset but missed here are highlighted in gray. Primitive erythroid cells are not present at these timepoints and testis & adrenal cells are collapsed to the epithelial cells major cell cluster due to their low numbers. **c**, Re-embedded 2D UMAP of 19,696 cells of the adipocyte major cell cluster. **d**, Re-embedded 2D UMAP of 7,986 cells of the lung & airway major cell cluster. **e**, For these three major cell clusters, we co-embedded cells from three vaginally delivered pups (samples 1-3 in Fig. 7c) and six pups delivered by C-section (samples 4-9 in Fig. 7c), followed by subsetting a uniform number of cells per sample. For cells from each of the three vaginally delivered pups, we calculated the number of their 10 nearest neighbors in the PCA embedding (*n* = 30 dimensions) from other samples. **f**, Re-embedded 2D UMAP of cells from these three major cell clusters, based on cells from three vaginally delivered pups and six pups delivered by C-section. For each row, the same UMAP is shown multiple times, with colors highlighting cells from individual pups (or two pups, in the case of the 0-min C-section timepoint).

**Supplementary Figure 18.**
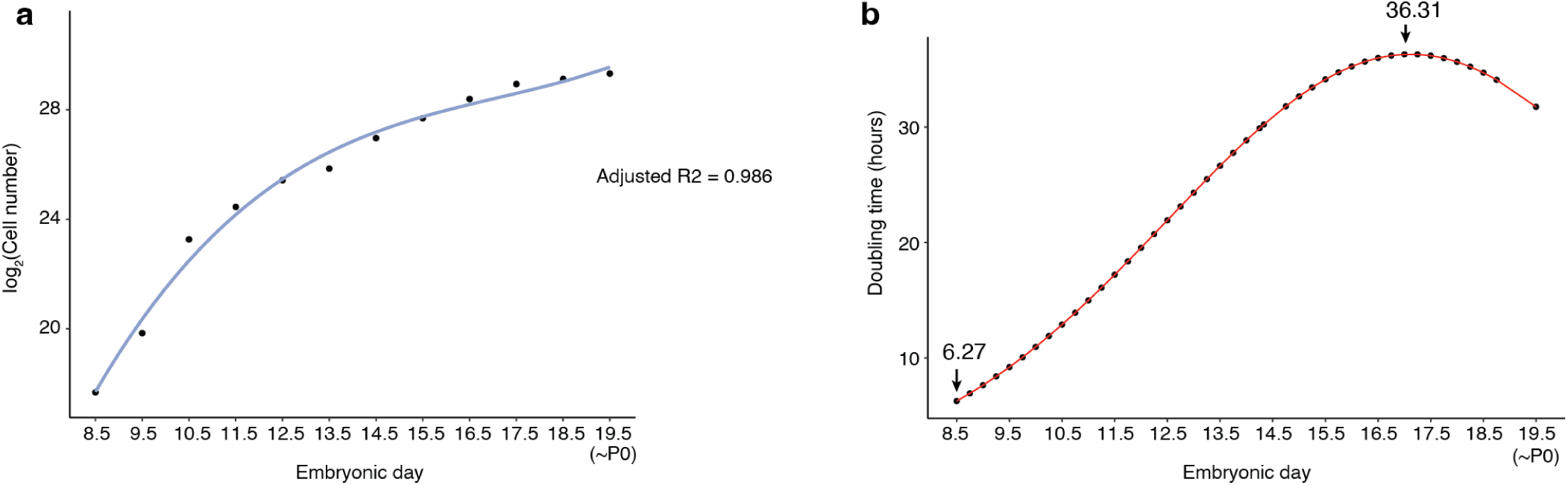
Quantitatively estimating cell number for individual mouse embryos as a function of developmental stage. **a,** Based on the experimentally estimated cell numbers of the 12 embryos (ranging from E8.5 to P0), we applied polynomial regression (degree = 3) to fix a curve across embryos between the embryonic day and log2-scaled cell number. P0 was treated as E19.5 in the model. **b**, The estimated “doubling time” of the total cell number in a whole mouse embryo are plotted as a function of timepoints. The timepoints with the longest (E17.0) and shortest (E8.5) estimated “doubling times” are highlighted.

